# CDK4 phosphorylation status and rational use for combining CDK4/6 and BRAF/MEK inhibition in advanced thyroid carcinomas

**DOI:** 10.1101/2023.06.05.543656

**Authors:** Jaime M. Pita, Eric Raspé, Katia Coulonval, Myriam Decaussin-Petrucci, Maxime Tarabichi, Geneviève Dom, Frederick Libert, Ligia Craciun, Guy Andry, Laurence Wicquart, Emmanuelle Leteurtre, Christophe Trésallet, Laura A. Marlow, John A. Copland, Cosimo Durante, Carine Maenhaut, Branca M. Cavaco, Jacques E. Dumont, Giuseppe Costante, Pierre P. Roger

**Affiliations:** Institut de Recherche Interdisciplinaire en Biologie Humaine et Moléculaire (IRIBHM) and ULB-Cancer Research Center (U-CRC), Université Libre de Bruxelles (ULB), Brussels, Belgium; Department of Pathology, Lyon Sud Hospital, Claude Bernard Lyon 1 University, Lyon, France; BRIGHTCore, ULB, Brussels, Belgium; Tumor Bank of the Institut Jules Bordet Comprehensive Cancer Center – Hôpital Universitaire de Bruxelles, ULB, Brussels, Belgium; Department of Head & Neck and Thoracic Surgery, Institut Jules Bordet Comprehensive Cancer Center – Hôpital Universitaire de Bruxelles, ULB, Brussels, Belgium; Tumorothèque du Groupement de Coopération Sanitaire-Centre Régional de Référence en Cancérologie (C2RC) de Lille, Lille, France; Department of Pathology, Univ. Lille, CNRS, Inserm, CHU Lille, UMR9020-U1277-CANTHER-Cancer Heterogeneity, Plasticity and Resistance to therapies, F-59000 Lille, France; Department of General and Endocrine Surgery – Pitié-Salpêtrière Hospital, Sorbonne University and Department of Digestive, Bariatric and Endocrine Surgery – Avicenne University Hospital, Paris Nord – Sorbonne University, Assistance Publique des Hôpitaux de Paris, Bobigny France, Paris, France; Department of Cancer Biology, Mayo Clinic, Jacksonville, United States of America; Department of Translational and Precision Medicine, Sapienza University of Rome, Rome, Italy; Molecular Endocrinology Group, Unidade de Investigação em Patobiologia Molecular (UIPM), Instituto Português de Oncologia de Lisboa Francisco Gentil (IPOLFG), Lisbon, Portugal; Departments of Endocrinology and Medical Oncology, Institut Jules Bordet Comprehensive Cancer Center – Hôpital Universitaire de Bruxelles, ULB, Brussels, Belgium

**Author notes:** In memoriam (Jacques E. Dumont). J.M. Pita and P.P. Roger are co-corresponding authors. Address for all correspondence related to this submission to: P.P. Roger, IRIBHM, Université Libre de Bruxelles, Campus Erasme, 808 route de Lennik, B-1070 Brussels, Belgium.

**Keywords:** ATC, PDTC, CDK4 Thr172-phosphorylation, palbociclib, trametinib, dabrafenib

## Abstract

Despite overall good prognosis associated to thyroid cancer (TC), poorly differentiated carcinomas (PDTC) and anaplastic carcinomas (ATC, one of the most lethal human malignancies) represent major clinical challenges. We have shown that the presence of active T172-phosphorylated CDK4 predicts sensitivity to CDK4/6 inhibitory drugs (CDK4/6i) including palbociclib. Here, CDK4 phosphorylation was detected in all well-differentiated TC (n=29), 19/20 PDTC, 16/23 ATC, and 18/21 TC cell lines including 11 ATC-derived ones. The cell lines lacking CDK4 phosphorylation were insensitive to CDK4/6i. RNA-sequencing and immunohistochemistry revealed that tumors and cell lines without phosphorylated CDK4 presented very high p16*^CDKN2A^* levels that were associated with proliferative activity. No *RB1* mutations were found in 5 of these 7 tumors. p16/KI67 immunohistochemistry and a previously developed 11-gene signature identified the likely insensitive tumors lacking CDK4 phosphorylation. In cell lines, palbociclib synergized with dabrafenib/trametinib, completely and irreversibly arresting proliferation. The combined drugs prevented resistance mechanisms induced by palbociclib, most notably Cyclin E1-CDK2 activation and a paradoxical stabilization of phosphorylated CDK4 complexes. Our study supports the evaluation of CDK4/6i for ATC/PDTC treatment, including in combination with MEK/BRAF inhibitors.

## 1. Introduction

Despite overall good prognosis associated to thyroid cancer, the management of patients with advanced thyroid tumors represents a major clinical challenge. Indeed, up to 10% of well-differentiated thyroid carcinomas (WDTC), initially cured with surgery followed by radioactive iodine treatment (RAI), may develop locally advanced or metastatic disease. Sixty to seventy percent of these tumors eventually become RAI-refractory [1], with a dramatic impact on survival. On the other hand, poorly differentiated carcinomas (PDTC) and anaplastic carcinomas (ATC), composed by dedifferentiated cells, primarily present as highly aggressive tumors. ATC has been claimed as one of the most lethal tumor types, contributing up to 50% of the deaths attributable to thyroid cancer, with patients displaying a median survival of 3 to 5 months [2,3]. In the majority of cases, tumor resection is not possible and response to chemo– or radio-therapy is poor [2,4,5]. PDTC and differentiated high grade carcinomas exhibit intermediate behavior and prognosis between WDTC and ATC [6,7].

In the last decade, targeted therapies have improved the management of RAI-refractory recurrent/metastatic WDTC and PDTC, but the toxicity associated to these treatments greatly impairs their clinical use [8]. Moreover, the new therapeutic strategies, including targeted therapy and immunotherapy, did not produce major improvement in terms of survival for most ATC patients [5,9,10].

Aberrant cell proliferation due to dysregulated cell division is a peculiar trait of cancer. The complexes between cyclins and cyclin-dependent kinases (CDK) play a pivotal role in the control of cell division and modulation of transcription in response to mitogenic factors. Particularly, CDK4 and CDK6 represent the master node of regulation for the G1 phase restriction point [11], by phosphorylating and initiating the inactivation of the retinoblastoma tumor-suppressor protein (RB). This process is subsequently maintained by a positive feedback loop linking RB to E2F-dependent transcription of *CCNE1* (cyclin E1), which activates CDK2. CDK4/6 assembly with a D-type cyclin is required for its activity and this binding can be counteracted by proteins of the INK4 family (such as p16 and p15), or can be facilitated by CIP/KIP family members (p21 and p27) [12,13]. Due to its deregulation, many cancer cells are addicted to CDK4/6 activity [14]. Selective inhibitors of CDK4/6 are being tested in numerous clinical trials against various types of cancers [15,16]. The CDK4/6 inhibitors (CDK4/6i) palbociclib, ribociclib and abemaciclib have become the standard of care for the treatment of Estrogen Receptor-positive advanced or metastatic breast tumors in combination with endocrine therapy, providing major improvements in progression-free and overall survival [17–20]. Still, indication for these drugs is limited by the lack of suitable biomarker of potential sensitivity. We have extensively demonstrated that phosphorylation on amino acid T172 is an absolute requirement for the activation of CDK4 and is finely regulated, determining the cell cycle commitment [11,13,21–24]. By contrast, the homologous T177 phosphorylation on CDK6 is not regulated and is generally absent or weak [13,25]. In our previous studies in breast tumors [26] and pleural mesotheliomas [27], we observed a variable presence of CDK4 T172-phosphorylation in most tumors. Its absence however, was also observed in some highly proliferative tumors, in association with the main mechanisms of resistance to CDK4/6i, including RB defects or inactivation, and high expressions of *CDKN2A* (p16), *CCNE1* (cyclin E1) and *E2F1* [26–29]. Hence, CDK4 T172-phosphorylation might be the most relevant biomarkers of potential tumor sensitivity to CDK4/6i, by identifying or predicting the presence of active CDK4, which is the actual target of inhibitory drugs. Furthermore, we showed that the CDK4 modification profile of breast tumors and cell lines can be predicted using the expression values of 11 genes [26].

PDTC and ATC are highly proliferative cancers associated with increased cell cycle progression and chromosomal instability [30–34]. Several studies from our group indicated CDK4 as a critical regulator of physiological and cancerous thyrocyte proliferation [13,21,35,36]. Moreover, MAPK/ERK, EGFR/ERBB and PI3K/AKT, widely recognized as the main deregulated signaling pathways in thyroid tumors [37,38], have CDK4/6 as major integrating node. It could therefore be speculated that inhibition of CDK4/6 might effectively represent a therapeutic approach for advanced thyroid tumors.

The aims of this work were to define if the CDK4 phosphorylation can be detected in thyroid cancer, whether it is variable and indicative of sensitivity to CDK4/6i and predictable on a biomarker based upon protein and/or mRNA expression. Furthermore, we aimed to identify drug combinations including CDK4/6i, which actively block the growth of advanced thyroid cancers, providing the rationale for testing in clinical trials.

## 2. Materials and methods

### 2.1 Human thyroid tissue samples

Thyroid tumors and normal thyroid tissues from flash-frozen samples or optimal cutting temperature (OCT) compound-embedded samples were obtained from different institutions. This study was performed in accordance with the Declaration of Helsinki and collection of patient tissues and associated data was done in agreement with the Ethics Committees of Jules Bordet Institute (CE1978, CE2970) with informed consent of patients, Sapienza University of Rome, Pitié-Salpêtrière Hospital and with the Mayo Clinic Institutional Review Board protocol. Samples and associated data from the Institute – Instituto Português de Oncologia de Lisboa Francisco Gentil (IPOLFG) – were obtained in compliance with all applicable laws, including a written consent, and the study was approved by the institute’s Ethics Committee. Written informed consent was obtained from all patients of the tumor tissue bank – Tumorothèque Centre de Ressources Biologiques des Hospices Civil de Lyon. Samples and associated data were obtained from the tumor tissue bank – Tumorothèque ALLIANCE CANCER de Lille – that operates under the authorization AC-2018-3110 granted by the French ministry of research. Prior to scientific use, patients were appropriately informed and asked to consent in compliance with the French regulations.

All ATC and PDTC diagnoses were histologically re-evaluated and confirmed by an expert thyroid pathologist (MD-P), by examination of corresponding FFPE slides or, whenever possible, of serial sections of OCT slides. PDTC were defined following the Turin proposal [39].

A total of 140 samples were collected for two-dimensional gel (2D-gel) electrophoresis analysis (42 ATC, 30 PDTC, 8 follicular thyroid carcinoma (FTC), 9 oncocytic FTC, 23 papillary thyroid carcinoma (PTC) (1 metastasis), 7 lymph node metastases and 21 normal thyroid tissues). Twenty-one samples were discarded due to signs of wide necrotic tissue (more than 80% of tissue area; 2 ATC), misclassification (1 ATC and 1 oncocytic FTC)), bad RNA quality (RNA integrity number lower than 3)/protein quality (3 PTC, 8 ATC, 1 normal and 3 oncocytic FTC) or insufficient protein quantity (1 PTC and 1 PDTC). Eighty-four of 119 samples were selected for RNA-seq, and following an exploratory analysis, 27 cases were further excluded from analysis due to: gross contamination/misclassification (3 normal with neoplastic cells, 1 PDTC with no neoplastic signs, 2 ATC with more than 70% of PTC cells, 1 ATC with 60% of stroma area); sample duplicate (1 normal); low RNA quality (RNA integrity number lower than 4.5; 1 ATC and 2 PDTC); high proportion of PCR duplicates (more than 0.5% of the library; 1 FTC and 2 oncocytic FTC); or very low/undetectable 2D-gel signal (4 ATC, 6 PDTC and 2 PTC). In summary, 98 samples were included in the 2D-gel proteomic analysis and 57 samples were included in the transcriptomic analysis.

### 2.2 Thyroid cancer cell lines

#### 2.2.1 Cell culture and inhibitors

Human thyroid carcinoma cell lines were maintained in a humidified atmosphere (5% CO_2_) at 37°C and cultured as described in Supplementary Table S1. The origin, authentication, and main features of each cell line are also provided. Cells were passaged for fewer than 2 months or within 20 passages. Cell culture reagents were obtained from Gibco, (Carlsbad, CA, USA). Palbociclib (PD0332991; S1116), abemaciclib (LY2835219; S7158), trametinib (GSK1120212; S2673) and dabrafenib (GSK118436; S2807) were purchased from SelleckChem (Houston, TX, USA) and dissolved in DMSO. Controls were treated with the same concentration of DMSO.

#### 2.2.2 DNA synthesis and cell growth assays

For DNA synthesis assay, cells seeded in triplicates in 96-well plates were incubated for at least 16 h before being challenged with the indicated serial dilutions of inhibitors for 24 h. One hour before fixation with methanol, 100 µM 5-bromo-2′-deoxyuridine (BrdU, Sigma-Aldrich, St Louis, MO, USA) and 4 µM 5-fluoro-2′-deoxyuridine (FldU, Sigma-Aldrich) were added to the cells. Immunodetection of DNA-incorporated BrdU was done as described [26,41], imaged and analyzed semi-automatically with a custom-made ImageJ macro as described previously [26].

The effect of inhibitory drugs on cellular growth was also evaluated using the SRB assay and MTT assay, as described previously [26]. Cells seeded in triplicates in 96-well plates were allowed to attach for at least 16 h before being incubated with serial dilutions of CDK4/6i for 144 h (SRB) or 48 h (MTT). Serial dilutions of puromycin were used as positive controls.

#### 2.2.3 Long-term cell treatments

To evaluate the effect of a prolonged treatment on DNA synthesis, cells seeded in duplicates in 3-cm dishes were incubated for at least 16 h before being treated with 1 µM palbociclib for 2 d or 10 d. After 10 d, cells were either fixed or washed twice with phosphate-buffered saline (PBS) and allowed to grow without drug for 1 d. Media and drugs were replenished every 3 d. One hour before fixation with methanol and permeabilization with 0.1% Triton X-100 (Sigma-Aldrich), 40 µM 5-ethynyl-2’-deoxyuridine (EdU, Invitrogen, Waltham, MA, USA) was added to the cells. DNA-incorporated EdU was detected following the Click-iT assay protocol (Invitrogen). Briefly, fixed and permeabilized cells were incubated for 30 min with a Tris-buffered saline buffer supplemented in the following order, with 1 mM copper(II) sulfate (Sigma-Aldrich), 100 mM L-ascorbic acid (Sigma-Aldrich) and 5 µM Alexa Fluor 594 Azide (Thermo Fischer Scientific). DAPI (1 µg/ml) was used as nuclear counterstain and round coverslips were mounted in each dish with ProLong Gold Antifade mountant. Cells were observed under a microscope and, for each condition, a total of at least 500 cells was manually counted.

#### 2.2.4 Clonogenic assays

Equal amounts of cells (5×10^2^ to 2×10^3^/well depending on the cell line) were seeded in 6-well plates and allowed to attach for at least 16 h, before being treated with the indicated drugs. After 10 d, cells were either stopped or washed twice with PBS and allowed to grow without drugs for the same amount of time. Media and drugs were replenished every 3 d. Cells were then fixed with 4% paraformaldehyde (Sigma-Aldrich) and stained with 0.05% crystal violet solution (in distilled water, Sigma-Aldrich) for 30 min. After washing and air-drying, the plates were photographed.

#### 2.2.5 Senescence-associated β-galactosidase activity staining

Cells seeded in 6-well plates, at densities adapted for each condition (5×10^2^ to 2×10^4^/well), were incubated for at least 16 h before being treated with drugs. After 6 or 8 d (media and drugs replenished every 3 d), β-galactosidase activity in cells was detected using the Senescence β-galactosidase staining kit (Cell Signaling Technology, Danvers, MA, USA) at pH 6.0, following the manufacturer’s protocol. After incubation with the staining solution for 16-18 h at 37°C without CO_2_, the proportion of cells with developed blue color was quantified. For each condition, a total of at least 500 cells was counted manually using a light-field microscope.

#### 2.2.6 Combination index calculation

Analysis of drug synergy was done using the CompuSyn software (www.combosyn.com) which is based on Chou-Talalay’s combination index theorem [42]. The software uses a median-effect method that determines if the drug combination produces greater effects together than expected from the summation of their individual effects. The combination index (CI) values were calculated for the different concentration-effect plots (for each of the serial dilutions) based on the parameters derived from the median-effect plots of the individual drugs or drug combinations at the fixed ratios. The CI was calculated based on the assumption of mutually nonexclusive drug interactions. Definition of the degree of synergism in accordance to the CI value was based as described [43].

### 2.3 Protein analyses

The antibodies used in this work are listed in Supplementary Table S2. Equal amounts of whole-cell extract proteins or immunoprecipitates were separated by SDS-PAGE and immunodetected. For 2D-gel electrophoresis, cells were lysed in a buffer containing 7 M urea and 2 M thiourea. Ground tissues obtained from cryogrinding of flash-frozen tumor tissues and frozen tumor slides embedded in OCT (at least, 10 sections of 10 µm per sample) were solubilized as described [26]. Proteins were separated by isoelectric focusing on immobilized linear pH gradient strips (pH 5 to 8, Bio-Rad, Hercules, CA, USA) before separation by SDS-PAGE. Chemiluminescence images of the samples were acquired on films or with a Vilber-Lourmat Solo7S camera and quantified using the Bio1D software (Vilber-Lourmat, Marne-la-Vallée, France). The profile of CDK4 separated by 2D-gel electrophoresis has been characterized previously [13,26]. Quantification of the spot 2 and spot 3 volumes (corresponding to the two main modified forms of CDK4) were done from 16-bit scans of the 2D-gel immunoblots, with unsaturated signals. After linear correction of the background, the volume ratio (spot3/spot2) was used to define three types of CDK4 modification profiles: a profile A (for absent) was attributed when the ratio was below 0.02; a profile L (for low) was attributed when the ratio lied between 0.02 and 0.50; and a profile H (for high) was given when the ratio was equal or above 0.50.

Co-immunoprecipitations were performed as described [13,21]. RB-kinase activity of immunoprecipitated CDK complexes was measured by *in vitro* incubation with ATP and a fragment of RB, as described [21,36].

### 2.4 RNA-sequencing

#### 2.4.1 Library preparation and sequencing

Total RNA was isolated from cell lines using the RNeasy Mini Kit (Qiagen, Hilden, Germany). Total RNA from ground tissues (obtained from cryogrinding of flash-frozen tumor pieces) and from OCT-embedded frozen tissue slides (at least, 10 sections of 10 µm per sample) were first extracted with TRI Reagent Solution (Invitrogen) using a Potter-Elvehjem homogenizer with a motorized PTFE pestle, and then were purified with the RNeasy Mini kit (Qiagen) and on-column DNase digestion (RNase-free DNase Set, Qiagen) according to the manufacturer’s protocol. Alternatively, RNA and DNA were extracted with Buffer RLT Plus using the AllPrep DNA/RNA mini kit (Qiagen) according to the manufacturer’s protocol. RNA yield and purity were assessed using a Fragment Analyzer 5200 (Agilent Technologies, Massy, France). 10ng to 100 ng of RNA was used for cDNA libraries and sequences production, as previously described [27]. Homo_sapiens.GRCh38.90.gtf annotations and Homo_sapiens.GRCh38.90.dna.primary-assembly.fa sequence files downloaded from ftp.Ensembl.org were used for reads alignments. Transcript level counts were calculated with HTSeq and normalized to library size to obtain counts per 20 million reads (CP20M). Integrative Genomics Viewer software A. [44] (IGV version 2.12.3) was used for visualization and screening of mutations.

#### 2.4.2 Analysis of RNA-sequencing data

Principal component analysis (PCA) was performed in R (version 4.2.3) using the libraries FactoMineR [45] and factoextra. The gene expression profiles of all samples tested in the study were used. Genes were first filtered by removing those with a null mean expression. Next, gene expression values were scaled by dividing each expression value by the standard deviation of the expression values of the considered gene with the sweep function. The PCA function was used to compute the principal components explaining decreasing proportions of the variance. The two first recorded principal components were plotted with symbols corresponding to CDK4 profiles and colors to types using the ggplot2 package. As in the exploratory plot, several samples were not clustering with others of the same group, the outlier samples were removed prior to define 95% confidence ellipses. These include two normal samples clustering with PDTC or ATC, one ATC clustering with PTC and two PDTC clustering with ATC. Finally, exclusion from the selection was extended to the samples with bad quality proteomic profiles or RNA (as detailed in section 2.1). The final sample selection used to define the confidence ellipses included 52 tumors out of the 84 tumors analyzed. A new plot was drawn with the excluded samples and the ellipses described above.

The thyroid differentiation score (TDS) was defined as the mean z-score of the 16 genes previously described [46]. An epithelial-mesenchymal transition (EMT) score was obtained as previously described [47] (with the minor difference of including *OCLN* gene), by subtracting the mean z-score of 4 epithelial marker genes from the mean z-score of 13 mesenchymal marker genes. To obtain a gene z-score in each sample, gene expression in CP20M was first mean-centered and then divided by the corresponding standard deviation of the gene expression across all samples.

#### 2.4.3 Calculation of *CDKN2A* exon 1α expression

The pileup function of the R package Rsamtools was used to extract the coverage values at each position of the *CDKN2A* locus from each sample BAM file. Genomic coordinates of the whole locus and all introns and exons coding for p14 and p16 were extracted from ftp.Ensembl.org. A linear regression between the coverage at each intronic position and their gene coordinates was used to correct the coverage at each exon positions for background. When background estimates were higher than the observed coverage, the value was set to zero. As the best fitting between exon 2 coverage and the sum of the exons 1α and 1β coverage was obtained using maximum coverage values, this parameter was used. To be able to compare *CDKN2A* gene and exon 1α expression levels in each sample, exon 1α expression was calculated as a fraction of *CDKN2A* gene expression. This fraction was defined by the ratio of maximum coverage values between exon 1α and the sum of exons 1α and 1β in each of the samples.

#### 2.4.4 Prediction of the CDK4 modification profile

As previously detailed and characterized [26], a centroid method was used to predict the CDK4 profile whereby the expression profile of 11 genes (including *CDKN2A*) of an unknown sample was compared to three references built by computing, for each 11 genes, the average of their expression among prototype tumors with A, H or L CDK4 profiles. The predicted profile was the one corresponding to the A, H or L centroid with the highest correlation coefficient. As they were initially profiled with the Affymetrix platform [26], the breast tumor references used to predict the CDK4 modification profile were first adapted by using the RNA-seq expression data acquired with RNA extracted from the same samples. Raw thyroid RNA-seq CP20M expression values were compared to these adapted references by Spearman correlation. For the *CDKN2A* gene, these values were also corrected, as detailed in section 2.4.3, by calculating the contribution of the p16-specific exon 1α to the expression of the whole *CDKN2A* locus expression. In this case, all samples were scaled by the same factor such as the average expression *CDKN2A* exon 1α of profile A thyroid tumors was equal to the *CDKN2A* value of the reference for profile A breast tumors.

### 2.5 Targeted DNA-sequencing

#### 2.5.1 Preparation and sequencing

Genomic DNA was extracted with Buffer RLT Plus using the AllPrep DNA/RNA mini kit (Qiagen) or was purified from the remaining interphase/organic phase left from RNA extraction by performing an alkaline phenol-chloroform extraction. Briefly, back extraction buffer (4M guanidine-thiocyanate; 50 mM sodium citrate; 1 M Tris, pH 8.0) and 200 µl of chloroform were added, the mixture was agitated vigorously and was let to sit for 10 min. After centrifugation at 12000 g for 15 min at 4°C, the upper phase was kept and an equal volume of chloroform was added. After vigorously agitation and incubation for 3 min, samples were centrifuged as before. After precipitation by addition of equal volume of chilled isopropanol to the upper phase and washing 3 times with 70% ethanol, the pelleted DNA was dried at 55°C for 15 min and was resuspended in EB buffer (10 mM Tris-HCl, pH 8.5) with agitation at 60°C. DNA was quantified, and quality checked with the Quant-iT PicoGreen dsDNA Assay Kit (Thermo Fisher Scientific). Massive parallel sequencing was performed using targeted-capturing of the 165 genes included in the ‘Solid and Haematological tumours’ panel (BRIGHTCore, Brussels, Belgium). 150 ng of genomic DNA was fragmented and processed to construct libraries with barcodes, which were hybridized with the DNA panel. The libraries were sequenced on Illumina NovaSeq 6000 with a coverage of 1500×.

#### 2.5.2 DNA-sequencing analyses

We derived genome-wide copy number profiles using the off-target reads from our targeted panel by running ASCAT.sc (https://github.com/VanLoo-lab/ASCAT.sc). Briefly, ASCAT.sc first removes the on-target regions from the bed definition of the target with padding of 1000 bp upstream and downstream. Then it bins the rest of the genome in 30 kb bins and count the number of reads with MAPQ > 30 falling in each bin to obtain a read-count track genome-wide. We used a normal diploid sample (L2), previously profiled with the same panel [27], to normalize the read counts. The number of reads in each bin is divided by the average number to get a ratio r, which can be expressed as a function of the purity ρ, the average number of copies (or average ploidy ψ) and the local number of DNA copies nT: r = (n_T ρ + 2(1−ρ))/ψ. We take the log of this track and segment it with circular binary segmentation [48]. Then for each value of the ploidy ψ∈[1.5,5] by 0.01 and purity ρ∈[.2,1] by 0.01 we fit n_T of each segment from the first equation n_T = (ψr−2(1−ρ))/ρ. We calculate the sum of Euclidean distances between n_T and the closest integer values, that is, round (n_T), and then select the combination of values of ψ and ρ that minimizes this distance.

### 2.6 Immunohistochemical analysis

Hematoxylin/eosin (HE), KI67 and p16 immunohistochemistry (IHC) was performed with FFPE tissue sections on a fully automated BenchMark Ultra IHC system (Ventana, Roche Diagnostics, Basel, Switzerland) with the UltraView Universal DAB Detection Kit (Roche Diagnostics), using standard routine protocols. Otherwise, FFPE sections from Mayo Clinic center were processed as described [49] and probed with a different set of antibodies (Supplementary Table S2). Pictures were acquired with a NanoZoomer digital scanner (Hamamatsu Photonics, Hamamatsu, Japan) at 40× magnification or in an Aperio AT2 scanner (Leica Biosystems, Wetzlar, Germany) at 40x magnification (samples from IPOLFG institute).

The QuPath software [50] (version 0.4.0) was used for immunohistochemistry scoring analysis. For each section, the most appropriate tumor areas (without necrosis or fibrosis) were delimited and the positive cell detection command was applied using the sum of optical density, and the nuclear minimum area for detection was set between 10-25 µm^2^. A 0.2 intensity threshold for the nuclear mean DAB optical density was applied for KI67 positive detection. Three intensity thresholds were set for the cellular mean DAB optical density of p16 staining, categorizing cells into weak (0.2 ≤ threshold < 0.4), moderate (0.4 ≤ threshold < 0.6) and strong (threshold ≥ 0.6) positive detection. The H-score was calculated by the software, which is a weighted sum (1 for weak, 2 for moderate and 3 for strong) of the percentage of cells in each intensity class, ranging from 0 to 300.

### 2.7 Response curves and statistical analysis

Response curves and statistical analyses were performed using GraphPad Prism version 6.0 (GraphPad Software, La Jolla, CA, USA). Half-maximal inhibitory concentrations of the cell proliferation (GI50) and 95% confidence intervals were estimated by fitting the data to the logarithm of concentration vs. normalized response model, with standard slope (Hill slope of –1.0) and using a least-squares fitting method. The two-sided unpaired Student’s t-test was used for comparison between two groups. Multiple group comparisons were done using the Kruskal–Wallis test followed by Dunn’s multiple comparison tests. Correlations were evaluated with the Pearson correlation (two-sided). Comparison of Kaplan–Meier survival curves was performed using the Mantel–Cox log-rank test. The Fisher’s exact test (two-sided) was used for the analysis of contingency tables. A p-value ≤ 0.05 was considered statistically significant.

## 3. Results

### 3.1 CDK4 phosphorylation is detected in most thyroid tumors and its absence is associated to higher expressions of *CCNE1*, *E2F1* and p16*^CDKN2A^*together with lower *RB1* levels

To evaluate the proportion of patient thyroid tumors that would be potentially responsive or intrinsically resistant to CDK4/6 inhibition, we determined the CDK4 modification profile and the presence of activated T172-phosphorylated CDK4 from a cohort of fresh-frozen samples (n=98) representative of the different subtypes of thyroid tumors (Fig. 1A, Supplementary Fig. S1 and Table S3). CDK4 was detected by immunoblotting from whole protein extracts separated by 2D-gel electrophoresis. CDK4 was thus resolved by its charge into three main forms as first characterized in thyroid primary cultures [13,21,51] and in other tumor types [26,27]. The most basic form (spot 1) is the native CDK4. The most acidic form (spot 3) increases in response to *in vitro* proliferation stimulus of various cells including thyrocytes [13,21,24,35,51–53] and has been identified as the highly regulated T172-phosphorylated CDK4 form, using several approaches including T172-phosphospecific antibodies [11,22,26,54]. To evaluate the relative level of CDK4 activation, we compared the presence of this T172-phosphorylated form to another modified CDK4 form (spot 2). This latter form does not incorporate [32P] phosphate and binds to p16 but only weakly to cyclins D [13]. The abundance ratio of the T172-phosphorylated form (spot 3) over the form separated in spot 2 allowed us to define three types of profiles (L for low, H for high and A for absent phosphorylated CDK4), as previously reported [26,27]. All WDTC exhibited the T172-phosphorylated form of CDK4 (n=29). This form was also present in all lymph node metastases (n=7) and in one available PTC distant metastasis, with higher proportion of profiles H in these groups of samples (50%), than in primary WDTC (25%). In advanced dedifferentiated tumors, phosphorylated CDK4 was present in 95% of PDTC (n=20) but only in 70% of ATC (n=23). On the other hand, as expected with quiescent tissue, normal thyroid samples (n=18) predominantly lacked CDK4 phosphorylation.

**Figure 1.**
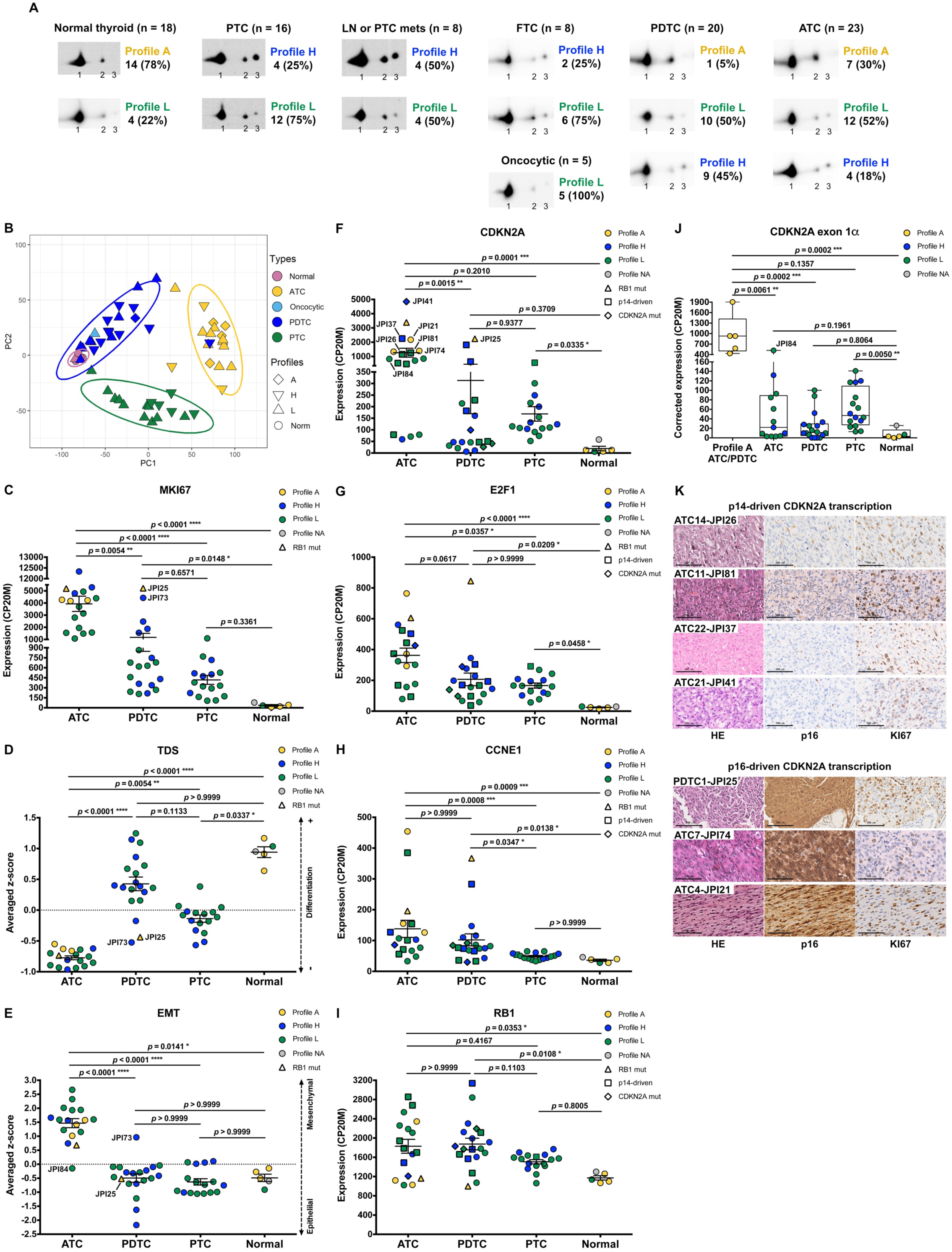
CDK4 phosphorylation is detected in most thyroid tumors and its absence is associated to higher expressions of CCNE1, E2F1 and p16*^CDKN2A^* together with lower RB1 levels. (A) Representative immunodetections of CDK4 after 2D-gel electrophoresis separation of whole protein extracts from fresh-frozen samples of normal thyroid tissues and different thyroid tumors subtypes. CDK4 modification profiles (number of cases and fraction in each subtype of tissue shown on the right side of immunoblots) were defined based on the ratio of T172-phosphorylated form of CDK4 (spot 3) over another modified form (spot 2), quantified from the immunoblots. H for high CDK4 phosphorylation: ratio ≥ 0.5; L for low CDK4 phosphorylation: 0.02 ≤ ratio < 0.5; A for absent CDK4 phosphorylation: ratio < 0.02. Unmodified native form of CDK4 is labelled as spot 1. (B) Representation of the gene expression variance between RNA-seq analyzed samples by principal component (PC) analysis (lymph node metastasis assigned as PTC). Computed 95% confidence ellipses for each sample subtype are plotted. (C-I) Expression levels of *MKI67*, thyroid differentiation score (TDS), epithelial-mesenchymal transition score (EMT), *CDKN2A*, *E2F1*, *CCNE1* (cyclin E1) and *RB1* in the samples grouped into normal tissues and the different subtypes of tumors (lymph node metastasis assigned to PTC). Expression levels calculated from the RNA-seq data and expressed as counts per 20 million reads (CP20M) or as averaged Z-scores of CP20M. Samples defined as presenting p14-driven *CDKN2A* transcription were checked in Integrative Genomics Viewer (IGV) to have exclusive expression of the exon 1β of *CDKN2A* gene. Error bars: mean +/-SEM. Statistical significance was calculated with the Kruskal-Wallis test corrected by Dunn’s multiple comparison tests. (J) Expression level of *CDKN2A* exon 1α in each group of samples, calculated from the RNA-seq data and expressed as the contribution of the exon 1α to the expression of the locus (in CP20M). Statistical significance between profile A group or normal tissues and other groups was calculated with the Kruskal-Wallis test corrected by Dunn’s multiple comparison tests. (K) Representative immunohistochemistry images of hematoxylin/eosin (HE), p16 and KI67 staining in 7 tumors with comparably high *CDKN2A* mRNA levels, as highlighted in panel F. Scale bars = 100 µm. *p<0.05, **p<0.01, ***p<0.001, ****p<0.0001; NA, not applicable.

In order to evaluate the molecular features associated with the different CDK4 profiles, we analyzed a sub-cohort of 84 cases by RNA-seq (complete gene expression data provided in Data set 1). An exploratory classification of the gene expression profiles by PCA revealed misclassified outliers (Supplementary Fig. S2). After excluding the samples with major technical issues and/or clear misclassification (as detailed in section 2.1), further analysis was done for 57 selected cases (Supplementary Table S4). As expected [37,55–57], we observed that samples were mainly separated according to their tumor sub-types (Fig. 1B). No clear segregation of samples according to the CDK4 modification profile was observed; nevertheless, most PTC samples with CDK4 profile H were closer to ATC samples (Fig. 1B). Evaluation of the proliferation with the *MKI67* gene expression (Fig. 1C), of the thyroid differentiation (TDS, Fig. 1D) and of the epithelial-mesenchymal transition scores (EMT, Fig. 1E) validated the identity of ATC cases. As expected, these were the most proliferative samples and had lost thyroid identity with acquisition of mesenchymal features. PDTC and PTC samples although being very heterogeneous, showed higher *MKI67* expressions and lower differentiation than normal tissues. Paradoxically, PDTC tended to have a higher differentiation score than PTC, as reported by others [37]. Also, worth mentioning is that the only PDTC case with CDK4 profile A (sample JPI25) had the highest *MKI67* expression and the second lowest thyroid differentiation score of PDTC group (Fig. 1C, D). Still, its EMT score (Fig. 1E) was similar to other PDTC (in contrast to sample JPI73 that is probably a misclassified ATC (Supplementary Fig. S2)).

Search for mutations (synonymous mutations excluded) in the main genes involved in thyroid cancer and in cell cycle, revealed *TP53* mutations as the major oncogenic driver for ATC, being present in 65% of cases (Supplementary Table S5). For PDTC, *NRAS* and *TP53* mutations were present in 21% and 37% of the tumors, respectively. As previously observed [32], these mutations were mutually exclusive and together affected more than half of the PDTC. *PTEN* mutations were also present in 21% of PDTC. Interestingly, these appeared to co-occur with *TP53* mutations (in 3 of 4 *PTEN* mutated cases). Mutations in other genes were involved in less than 20% of the ATC and PDTC. *BRAFV600E* was the only driver mutation identified in PTC, lymph node or PTC metastases, being present in 88% of samples. Among the five tumor samples with no phosphorylated CDK4 (profile A), we identified two samples with mutated *RB1* (a nonsense and a frameshift mutation). Three of these profile A samples had *TP53* mutations.

As in our previous studies in breast tumors [26] and pleural mesotheliomas [27], we observed the lack of CDK4 T172-phosphorylation in some thyroid tumors to be associated with high expressions of *CDKN2A* (p16), *CCNE1* (cyclin E1) and *E2F1*. Transcript levels of *CDKN2A*, *E2F1* and *CCNE1* were indeed elevated in ATC and PDTC samples with profile A of CDK4 (Fig. 1F-H). High p16 cellular levels are expected to prevent the activating phosphorylation of CDK4 by impairing its binding to cyclins D. By contrast, in normal quiescent thyroid tissue, lack of CDK4 phosphorylation was likely due to the absence of mitogenic stimulation [13,21] because it was associated to a particularly low p16/*CDKN2A* expression, an intriguing feature already reported by others [35,58–61]. Overexpression of p16/*CDKN2A* in cancer cells has been generally related to *RB1* loss of function [28,62,63]. However, three of these profile A samples (with high *CDKN2A* expression) were ATC without any detectable *RB1* mutation (samples JPI21, JPI27 and JPI74). Lack of *RB1* mutation was confirmed with targeted DNA-sequencing (Supplementary Fig. S3) in two samples (JPI21 and JPI27, although the latter showed *RB1* hemizygous loss). Two other cases of profile A ATC, which could not be RNA-sequenced, also revealed an absence of *RB1* mutations or loss. The copy number profile of the profile A PDTC (JPI25) was very complex (Supplementary Fig. S3) and we could not evaluate if the very low *RB1* variant allelic fraction of 7% (compared to the *TP53* variant allelic fraction of 25%) suffice to explain the high expression of *CDKN2A* and lack of CDK4 phosphorylation. Indeed, the RB alteration might be a subclonal event in this sample. Nevertheless, samples with profile A had a reduced *RB1* expression, comparable to the one observed in normal tissue samples (Fig. 1I).

Due to an alternative open reading frame (ORF), *CDKN2A* translates into two different proteins (p16 and p14). In contrast to breast tumors [26], high *CDKN2A* expression measured by RNA-seq analysis in thyroid tumors, did not entirely correlate with elevated p16 expression and lack of CDK4 phosphorylation. Among PDTC and ATC, eleven samples had elevated *CDKN2A* mRNA expression levels, but displayed CDK4 profiles H or L. In most of these cases, over-expressed *CDKN2A* mRNA seemed to mainly encode the p14 protein (only reads covering exon 1β, specific for p14 isoform, were observed in IGV) or would encode a truncated p16 protein (p.Q50* mutation). If we account for this by correcting *CDKN2A* transcription (as detailed in Materials and Methods), elevated expression of p16/*CDKN2A* was exclusively found in samples lacking phosphorylated CDK4 (Fig. 1J). Immunohistochemical staining corroborated the absence of p16 protein in samples with p14-driven *CDKN2A* transcription (Fig. 1K). By contrast, profile A samples with similar *CDKN2A* expression levels had strong p16 staining. Importantly, in all such cases, KI67 staining was similarly elevated in the p16-high tumor area, further demonstrating that p16 accumulation did not preclude tumor cell proliferation in these tumors.

### 3.2 Evaluation of p16/KI67 staining and a 11-gene signature as predictive biomarkers

To validate the high levels of p16 protein as a useful biomarker to identify tumors with no phosphorylated CDK4 (profile A), we performed immunohistochemistry staining in FFPE samples from 13 ATC and 17 PDTC (Supplementary Table S6 and Fig. S4). All six profile A tumors (5 ATC and 1 PDTC) were p16-positive, 3 of which with very strong staining, while 3 out of 12 profile L and 3 out of 11 profile H were also p16-positive in a significant proportion of cells. The remaining cases had no or very faint staining, and positivity appeared rather in the stroma. Staining of serial cuts for the proliferation marker KI67 revealed that profile A tumors had a higher proportion of KI67-stained cells and, more importantly, that p16-positive areas were co-stained for KI67. By contrast, no such co-expression of p16 and KI67 was observed in most profile H and L tumors (Supplementary Table S6). Only one profile L ATC case (JPI84, L* in Supplementary Table S6) exhibited areas with co-expression of both proteins, indicating that it could be a heterogeneous sample with profile A areas, which could also explain its very low ratio of CDK4 spot 3/spot 2.

We previously reported that the CDK4 modification profile of breast tumors and cell lines can be predicted using the expression values of 11 genes [26]. The tool was initially developed with Affymetrix microarrays. Our ongoing work (manuscript in preparation) indicates that the prediction tool can be run with gene expression profiles quantified with other technologies (RNA-seq, AmpliSeq) and adapted to RNA extracted from FFPE tumor samples. Therefore, we first tested whether the tool based on the genes (Supplementary Table S7) selected for the CDK4 profile prediction in breast tumors and the corresponding RNA-seq breast references could correctly predict the profiles of thyroid tumors, using raw CP20M gene expression values. The accuracy (proportion of predictions matching the observations) reached only 57.4% when applied to the prediction of the three CDK4 modification profile in thyroid tumors (Supplementary Table S8). Five samples were erroneously predicted as A profiles. Four of them were samples with exclusive contribution of exon1β (coding for p14) to the expression of the locus. In addition, profiles H or L were confounded in 19 tumor samples (7 L profiles predicted as H and 12 H profiles predicted as L). Upon use of the *CDKN2A* CP20M gene expression values corrected by the contribution of the exon1α to the expression of the locus, the overall accuracy raised to 68.5% for the prediction of the three profiles. As risk evaluation by distinguishing H from L profiles may not be practically useful in advanced thyroid cancer, we next checked whether a binary prediction of tumors as A or non-A (L or H) profiles might be more accurate. When a tumor was predicted to have A profile by a highest correlation with the A profile reference compared to the H or L profile references, the accuracy of the prediction, based on the exon1α *CDKN2A* contribution, raised to 98.2% (Supplementary Table S8). All observed A profile tumors were correctly predicted and only one sample (the PTC lymph node metastasis JPI38) was falsely predicted as A profile.

### 3.3 Higher levels of CDK4 phosphorylation are associated to higher proliferative potential and worse clinical outcomes

ATC with CDK4 profile H were significantly more proliferative than profile L cases (*p* = 0.0063), as judged by the higher expression of *MKI67* (Fig. 2A, Supplementary Fig. S5A). A similar tendency could be seen for PDTC (*p* = 0.0854). However, we noticed that only a subset of profile H PDTC samples had higher *MKI67* expression levels than profile L samples (Fig. 2A, Supplementary Fig. S5B). By contrast in PTC, several tumors displayed a H profile, but none was observed to be more proliferative than L profile ones (Fig. 2A, Supplementary Fig. S5C). Expression of *MKI67* was indeed significantly correlated with phosphorylated CDK4 levels in ATC (*p* = 0.0462) and PDTC (*p* = 0.0490) but not in PTC (*p* = 0.4329) (Supplementary Fig S5A-C). Nevertheless, PTC with profile H had significantly lower thyroid differentiation than profile L (*p* = 0.0289), a tendency that was not clearly associated to BRAF mutations (Fig. 2B). In each of the ATC and PDTC groups, samples with profile H showed significantly higher expression of *E2F1* than samples with profile L (*p* = 0.0288 and *p* = 0.001, respectively; Fig. 2C). *CCNE1* and *RB1* expressions did not differ significantly between the two profiles (Supplementary Fig. S5D, E).

**Figure 2.**
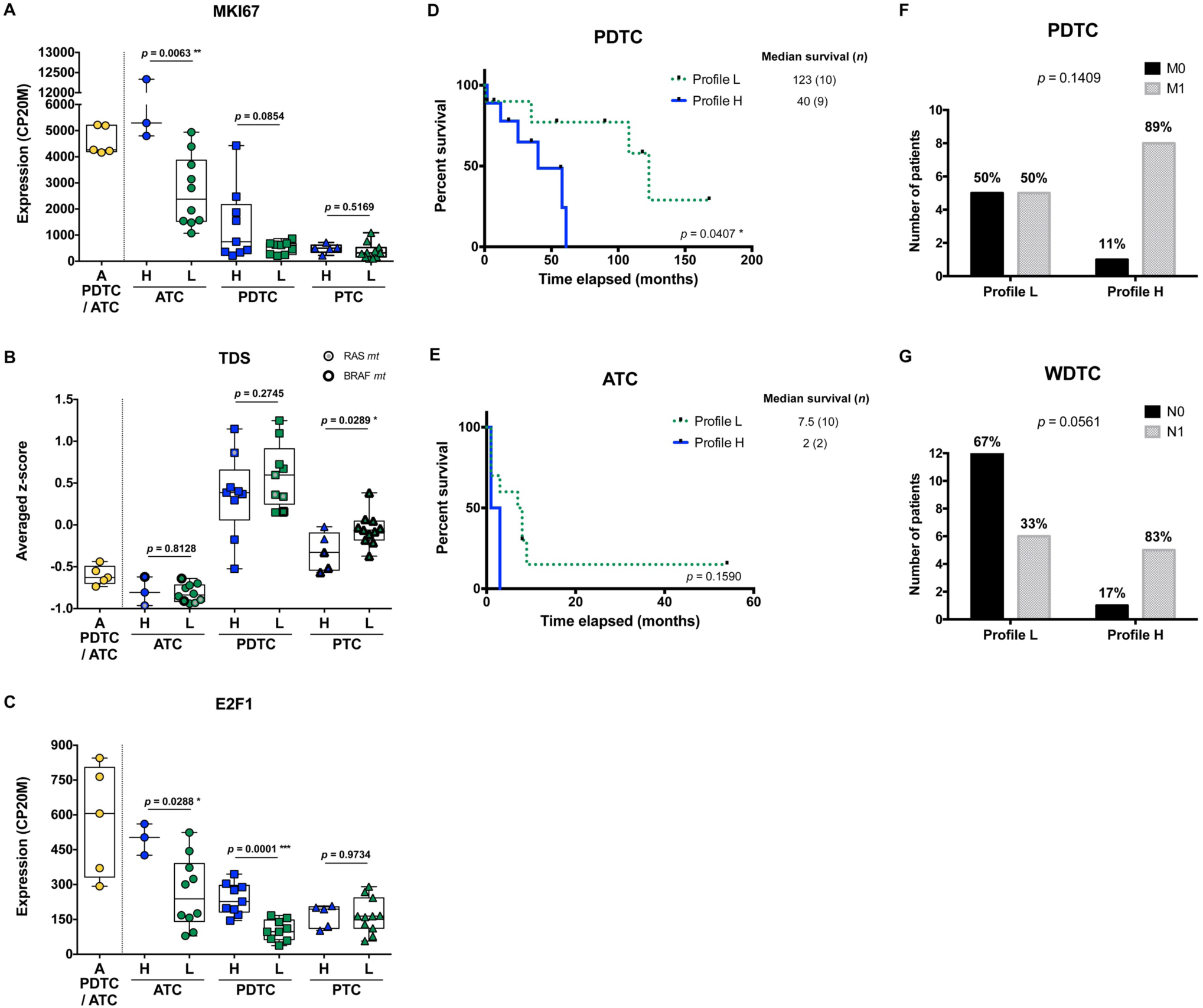
Higher levels of CDK4 phosphorylation are associated to higher proliferative potential and worse clinical outcomes. (A-C) Expression levels of *MKI67*, thyroid differentiation score (TDS) and *E2F1* in the samples grouped according to their CDK4 modification profiles (A, H or L as defined in Fig. 1A) in each subtype of tumors (lymph node metastasis assigned to PTC). Expression levels calculated from the RNA-seq data and expressed as counts per 20 million reads (CP20M) or as averaged Z-score of CP20M. Statistical significance between profile H and profile L groups in each type of tumors were calculated with unpaired t-test. (D, E) Kaplan–Meier curves comparing overall survival of ATC and PDTC patients with profile L or profile H of CDK4. Statistical significance was calculated using a log-rank test. (F, G) Proportion of PDTC or WDTC patients (patients with oncocytic FTC included), grouped according to their tumor’s CDK4 modification profile, for which distant (M) or regional nodal (N) metastases were assessed (0, not found; 1, found). Statistical significance was calculated with Fisher’s exact test. *p<0.05, **p<0.01, ***p<0.001.

PDTC patients, for which tumors exhibited a profile H, had significantly shorter overall survival than those with profile L tumors (*p* = 0.0407; Fig. 2D). In ATC, the median survival tended to be lower for profile H (2 months vs 7.5 months for profile L) (*p* = 0.1590; Fig. 2E). Similarly, PDTC and WDTC patients associated to profile H were more likely to have distant metastases (89% vs 50%; Fig. 2F) and nodal metastases (83% vs 33%; Fig. 2G), respectively. Therefore, even if it was not associated with a higher proliferation in WDTC, a higher CDK4 activation appeared associated to features of higher aggressiveness.

Unfortunately, completed clinical data could not be retrieved for several profile A tumor patients and several ATC patients were lost to follow-up. Remarkably, one of these profile A-ATC patients (JPI21) is still alive more than 8 years after diagnosis and chemotherapy.

### 3.4 Most thyroid cancer cell lines are sensitive to CDK4/6 inhibitors, which correlates with presence of phosphorylated CDK4 and can be predicted by the signature tool

Both commercial (n=17) and patient-derived cell lines [64] (n=4) from different subtypes of tumors were analyzed in order to characterize their response to two CDK4/6i – palbociclib and abemaciclib. Only three cell lines were intrinsically resistant to the drugs (Fig. 3A). For the remaining cells, including all ATC-derived lines, both inhibitors reduced S-phase entry (as measured by BrdU incorporation into DNA), in a dose-dependent way. Palbociclib’s half-maximal inhibitory concentrations of the cell proliferation (GI50) were similar to the ones reported in other types of cancer cells and had no apparent correlation with the cell line subtype or driver mutations (Fig. 3B). Nevertheless, six cell lines had a significant residual proliferation (defined by more than 20% of cells in S-phase, relatively to the control) at the maximum dose of 1µM of palbociclib (Fig. 3A). Assessment of cell viability by other methods, dependent on mitochondrial dehydrogenases activity (MTT) or on the cellular protein content (SRB), showed less pronounced effect (Supplementary Fig. S6), stressing the mainly cytostatic properties of CDK4/6i. CDK4 immunoblotting from 2D-gel electrophoresis revealed that T172 phosphorylation of CDK4 was present in all the sensitive cell lines. By contrast, the three completely resistant cells lacked the (active) phosphorylated CDK4 form (Fig. 3C). Further analysis by western blotting showed that these resistant cell lines had no RB or no phosphorylated RB expression, in accordance with the mutations previously reported [65] (a frameshift mutation in T243, a nonsense mutation in FTC-238 and two amino acids deletion in FTC-133) and confirmed by our RNA-seq data (Fig. 3D; Supplementary Table S2). The absence of functional RB in these cells probably accounted for the elevated expressions of p16 and Cyclin E1 observed in Fig. 3D. In almost all sensitive cell lines, p16 was affected either by mutation, deletion or promoter methylation of the p16-coding transcript and thus was not expressed (Fig. 3D, E). Nevertheless, *CDKN2A* transcripts were detected in most sensitive cell lines without reported *CDKN2A* gene deletion. Analogously to what we observed in profile L and H tumors with high *CDKN2A* transcript levels, *CDKN2A* expression was exclusively due to the alternative ORF p14 expression in these cell lines (Fig. 3E). Exon 1β, which is unique to p14 transcript, was expressed in more than half of the cell lines and, consistently, p14 was detected by western blotting (Fig. 3E). On the other hand, *CDKN2A* exon 1α, which is exclusively present in p16 transcript, was only expressed in 5 cell lines. In two of them (the sensitive HTC-C3 and THJ-29T cells), *CDKN2A* is mutated (H66R+D108N and G55Afs*91, respectively) and p16 protein was not detected. Cyclin E1 and CDK4 expressions were elevated in all the resistant cell lines and were more variable in sensitive ones (Fig. 3D). High residual proliferation was seen in cell lines with higher expression of Cyclin E1 (WRO, CAL-62 and IHH-4) or CDK4 (THJ-11T). At the transcript level, *CDK4* expression showed a significant negative correlation with the sensitivity of the cell lines and the expression of both *CDK4* and *CCNE1* were significantly correlated with the residual proliferation of the cell lines (Supplementary Fig. S7). The basal proliferation of the cell lines was also significantly correlated with the residual proliferation.

**Figure 3.**
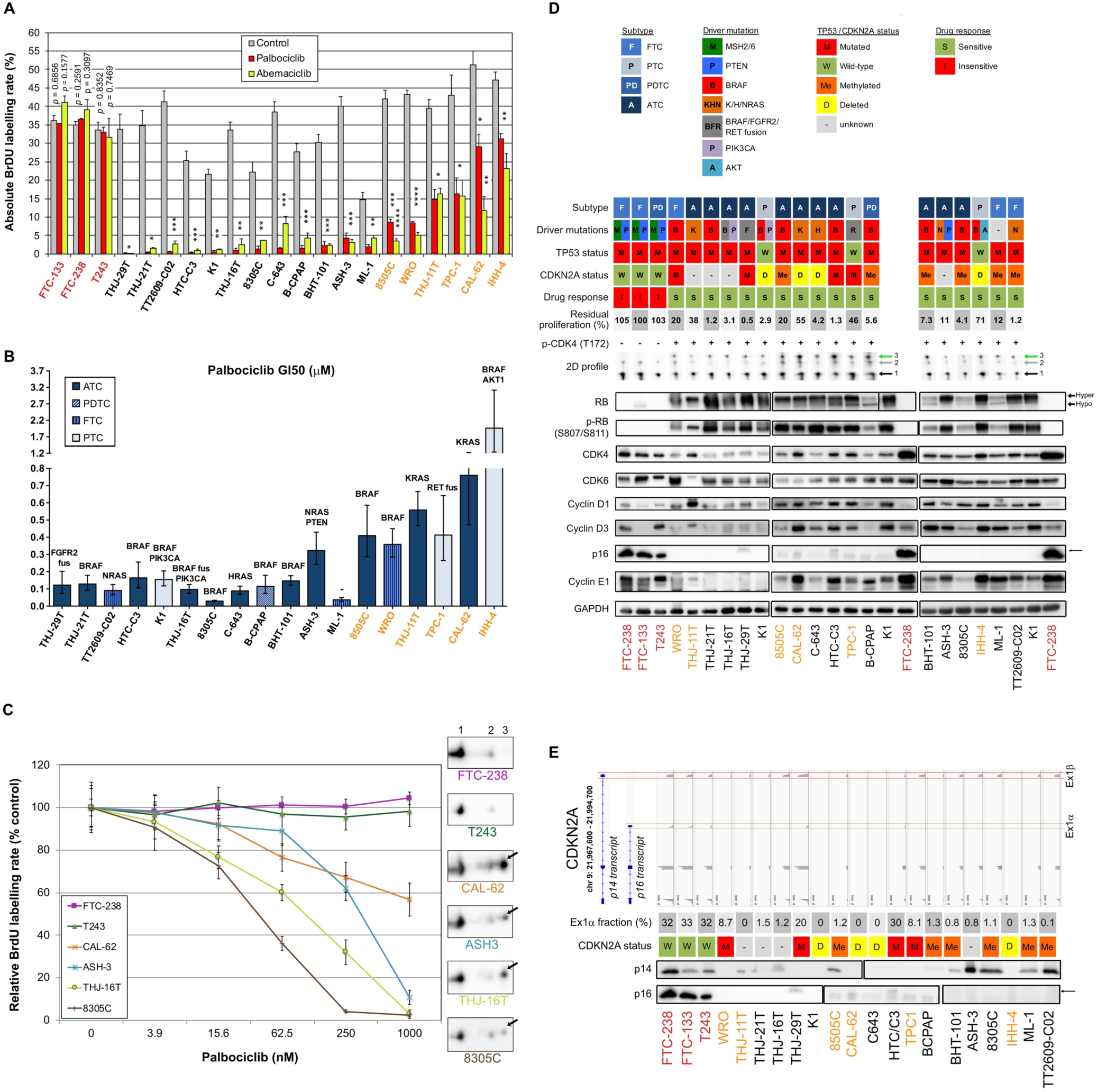
Most thyroid cancer cell lines are sensitive to CDK4/6 inhibitors, which correlates with presence of phosphorylated CDK4. (A) BrdU incorporation rate (during a 1 h pulse) following 24 h treatment with vehicle or with CDK4/6 inhibitors (1 µM). Pooled data from at least two independent experiments (error bars: mean ± SEM). Statistical significances were calculated by unpaired t-test. *p<0.05, **p<0.01, ***p<0.001, ****p<0.0001. (B) Half-maximal inhibitory concentration of the cell proliferation (GI50) determined from palbociclib’s response curves with serial dilutions of the drug. The main known driver oncogenes in each cell line are shown (gene fusions denoted as ‘fus’). Pooled data from at least two independent experiments were fitted by nonlinear regression model (error bars: best-fit value ± 95% confidence intervals). (C) Palbociclib’s response curves and relationship with T172 phosphorylation of CDK4. The illustrated cell lines are representative of the analyses performed in the 21 thyroid cancer cell lines. The 2D-gel electrophoresis and CDK4 immunodetection for each cell line is presented in the same font color as the correspondent response curve. BrdU incorporation was determined as in panel A with serial dilutions of the drug. T172-phosphorylated form (spot 3) is indicated by a black arrow. Native unmodified form of CDK4 and another modified form are labelled as spot 1 and spot 2, respectively. Pooled data from at least two independent experiments (error bars: mean ± SEM). (D) Western blot analyses of the indicated proteins extracted from cells cultured in control conditions. The vertical solid line in RB detection separate parts of the same blot detection that were re-assembled. K1 and FTC-238 were loaded on each gel to compare protein expression between the different gels. Relevant genomic features (from the analysis of the RNA-seq data and/or in accordance with Landa and colleagues [65]) and the 2D-gel electrophoresis profile of CDK4 (native form – spot 1, another modified form – spot 2, T172-phosphorylated form – green arrowed spot 3) are shown for each cell line. *CDKN2A* status refers to p16 protein and CpG methylated status at exon 1α (exclusive for p16 protein) was retrieved from the Cancer Cell Line Encyclopedia (CCLE) project. *CDKN2A* status was considered unknown for cell lines with no expression of p16 protein, with unknown methylation status and with no genomic aberration detected. Residual proliferation corresponds to the average relative BrdU labelling rate after 24 h treatment with palbociclib (1µM). Hyper– and hypo-phosphorylated forms of RB are indicated. Correct p16 band is indicated by a black arrow. (E) Illustration of the genomic organization of *CDKN2A* locus (not on scale) adapted from Integrative Genomics Viewer (IGV) visualization of each cell line’s BAM file. Coverage track of the transcripts from p14 and p16 proteins, and respective protein immunodetection by western blot (p16 detection as in panel D) are presented for each cell line. Exon 1α fraction calculated from the exon 1α coverage over the sum of exon 1α and 1β coverages. *CDKN2A* status defined as in panel D. Cell lines with residual proliferation ≥ 90% (resistant), ≥ 20% (sensitive with residual proliferation) or < 20% (sensitive) are indicated by red, orange or black font colors, respectively.

By contrast, the transcript levels of *CDK6*, *RB1* or *CCND1* (cyclin D1) were not correlated with the residual proliferation (Supplementary Fig. S7).

We next explored the impact of palbociclib treatment for 4 d (tested at 1 µM), on selected cell cycle related protein expressions (Fig. 4A). At this drug concentration, no alteration was seen in the resistant cells upon palbociclib treatment. This suggests that any effect observed in responsive cells was specific. In sensitive cells, we observed diminished RB expression/hyper-phosphorylation and decreased levels of Cyclin A2 and E2F1 (Fig. 4A). Palbociclib-treated cells exhibited elevated levels of CDK4, Cyclins D and E1. These observations are consistent with a cell cycle arrest at G1 phase. Drug-exposed cells observed at the microscope were flat, enlarged and seemed to have increased cellular complexity (not shown), suggestive of cells undergoing senescence. Palbociclib treatment induced higher senescence-associated β-galactosidase (SA-β-gal) activity in all the sensitive cell lines (Fig. 4B). However, this increase was detected in more than 30% of cells only in three cell lines (THJ-29T, HTC-C3 and BHT-101). Aside from decreased levels of PARP1 (E2F target already described as down regulated by CDK4/6 inhibition [16]), treated cells showed no appreciable evidence for increased apoptosis (no cleaved PARP1 nor cleaved caspase 3) or autophagy (minimally changed p62 and LC3B levels) (Supplementary Fig. S8). This is in agreement with previous studies, where no or minimal induction of apoptosis was seen in thyroid cancer cell lines [66–68]. Permanent arrest in the three cell lines with elevated SA-β-gal activity was further evidenced by reduced S-phase entry during prolonged palbociclib treatment (up to 10 d), which was maintained even after drug washout (Fig. 4C). However, in most of the other sensitive cell lines, palbociclib did not induce a durable cell cycle arrest. Indeed, the proportion of cells entering into S-phase raised when the treatment was prolonged, or rapidly increased once the compound was withdrawn (Fig. 4C).

**Figure 4.**
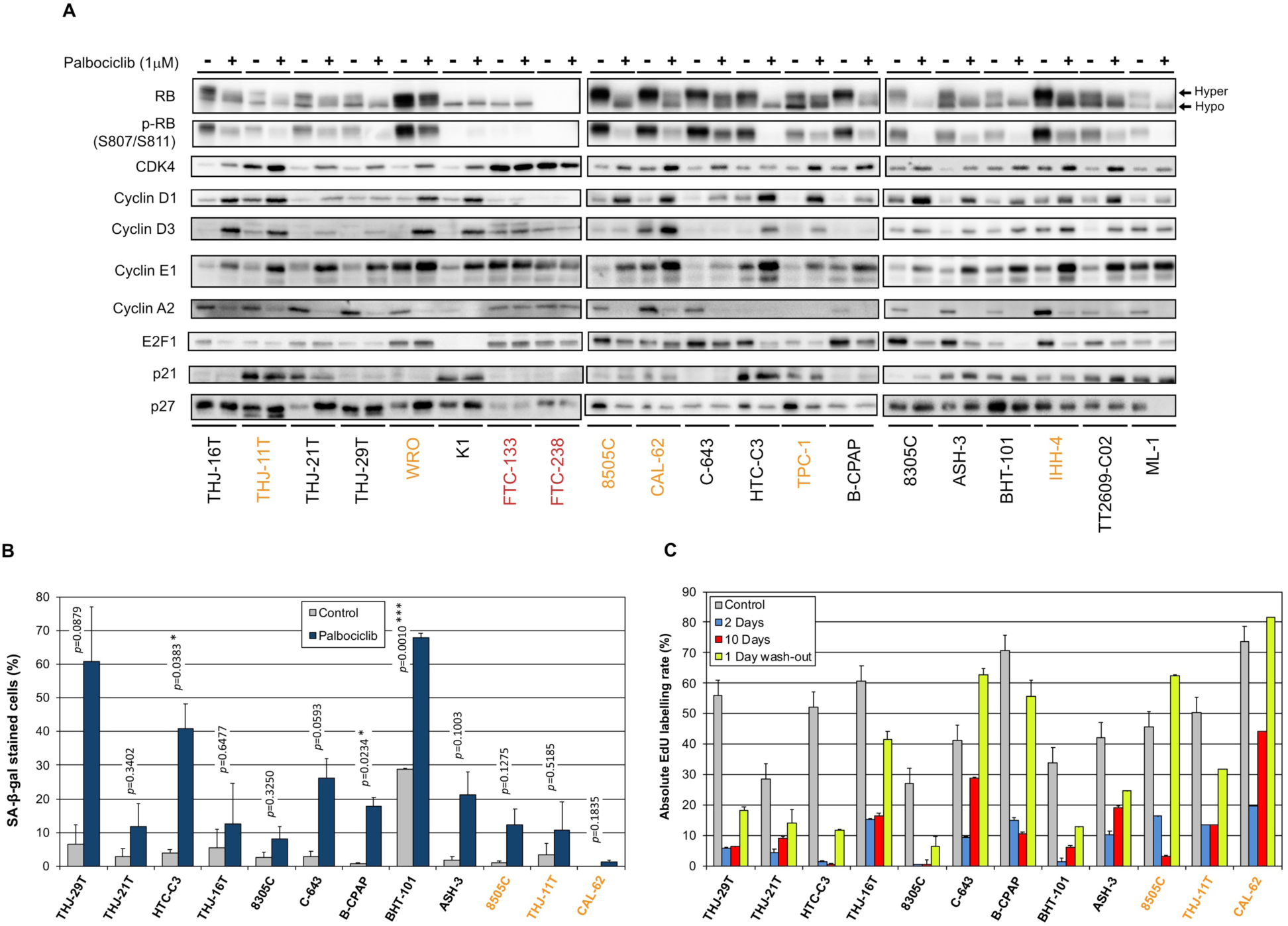
Thyroid cancer cell lines are responsive to palbociclib, but the induced cell cycle arrest is transient. (A) Indicated proteins were immunodetected after SDS-PAGE of total protein extracts from cells treated for 4 d with vehicle or with 1 µM palbociclib. Hyper– and hypo-phosphorylated forms of RB are indicated. (B) Cells stained for senescence-associated β-galactosidase (SA-β-gal) activity in ATC– and PDTC-derived cell lines were quantified following treatment for 6 d with vehicle or with 1 µM palbociclib. Pooled data from two independent experiments (error bars: mean ± SEM). Statistical significance was calculated by unpaired t-test. *p<0.05, ***p<0.001. (C) EdU incorporation rate (during a 1 h pulse) in ATC– and PDTC-derived cell lines, treated with vehicle for 11 d or treated with 1 µM palbociclib for 2 d, for 10 d or for 10 d followed by 1 d without drug. Error bars: mean ± SD (of duplicates; n = 1). Cell lines defined as resistant, with residual proliferation or sensitive (as detailed in Fig. 3) are indicated by red, orange or black font colors, respectively.

The 11-gene expression signature used to predict the CDK4 profile in tumors was further evaluated in these 21 thyroid cell lines (Supplementary Table S9), as performed for the tumors in section 3.2. Upon use of the *CDKN2A* gene expression values corrected by the contribution of the exon1α to the expression of the locus, the accuracy of the prediction of A or non-A CDK4 profiles raised to 90.5% (Supplementary Table S10). The three profile A cell lines were correctly predicted and only two cell lines were falsely predicted to have a profile A. These two cell lines (HTC-C3 and THJ-29T) have mutated *CDKN2A* with high expression of the locus. These data further validate the use of the gene expression signature to predict the presence of CDK4 phosphorylation and hence the potential sensitivity to CDK4/6 inhibition. Nevertheless, in case of high expression of *CDKN2A*, potential mutations of the locus will have to be identified to exclude this confounding factor.

### 3.5 Combination of CDK4/6 inhibitors with MEK/BRAF inhibition potentiates the growth-inhibitory effects and prevents the clonogenic ability of thyroid cancer cells

We have observed that MAPK/ERK activity was increased upon palbociclib treatment, as judged by the elevated phosphorylation levels of ERK in most sensitive cell lines, and this mirrored the effect seen for Cyclin D1 (Fig. 5A). In 12 of 18 sensitive cell lines, palbociclib also induced increased phosphorylation of p70S6 kinase and an electrophoretic mobility shift of this protein to a slower-migrating form, suggestive of hyperphosphorylation [69]. By contrast, a palbociclib effect on 4EB-P1 and ribosomal protein S6 (RPS6) was less apparent (Fig. 5A). We have thus investigated the possible synergism between CDK4/6 inhibition and BRAF and/or MEK inhibition using a combination of dabrafenib (BRAF inhibitor) and trametinib (MEK inhibitor). This dabrafenib/trametinib combination is the only targeted therapy approved by the FDA for the treatment of patients with locally advanced or metastatic ATC harboring *BRAF* V600E mutation [70]. In a subset of seven ATC-derived cell lines carrying different oncogenic drivers and having distinct responses to palbociclib (Sup Table S11), triple combined treatment potentiated drugs action, leading to complete suppression of proliferation (Fig. 5B). A strong synergistic effect was obtained, as judged by the calculated combination indexes (CI below one) (Fig. 5C). Double combination of trametinib and palbociclib was equally effective against cell lines possessing mutated oncogenes other than BRAF. The synergism was reached even in cells displaying partial response to palbociclib alone (8505C and CAL-62 lines). Identical drug cooperation was observed using either palbociclib or abemaciclib (Supplementary Fig. S9).

**Figure 5.**
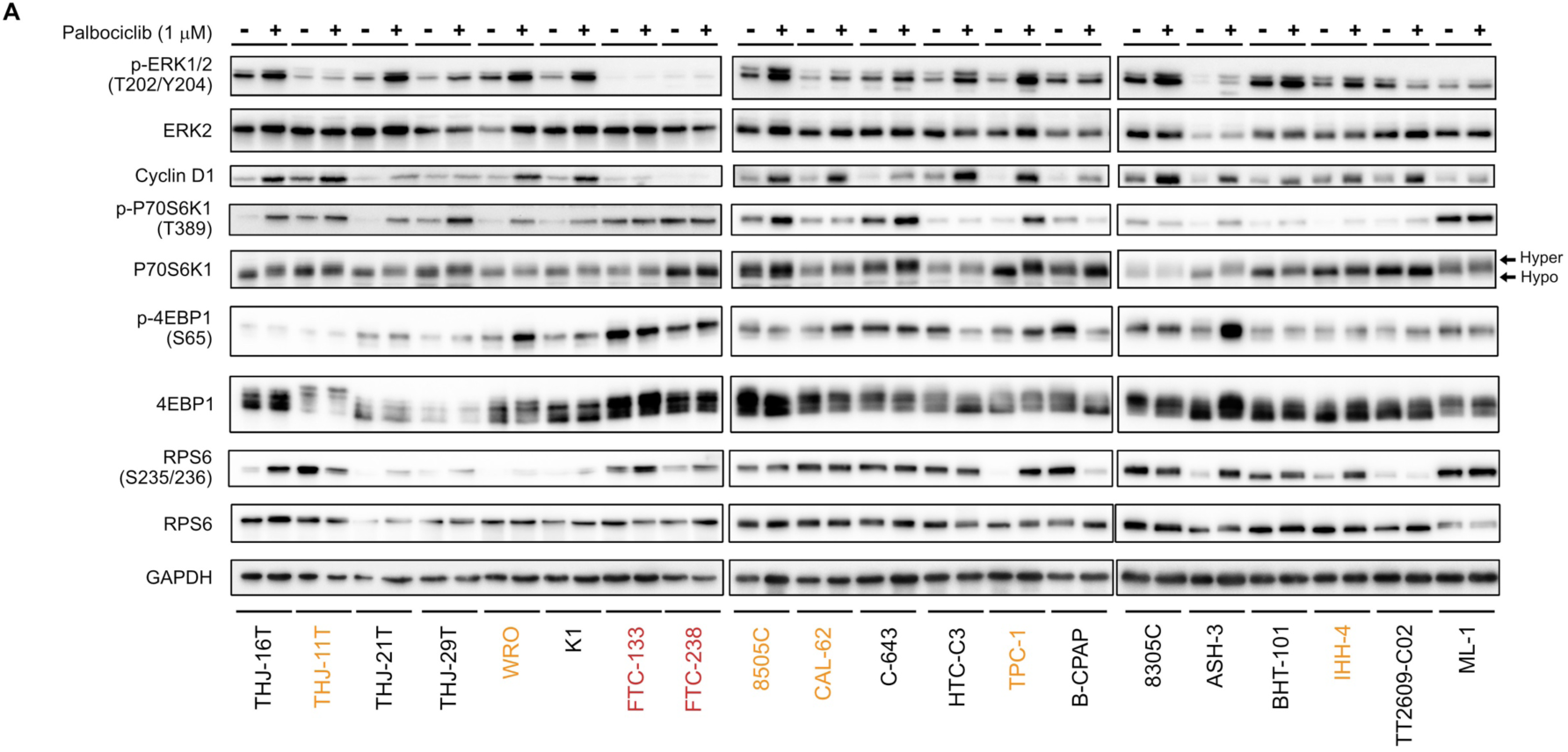
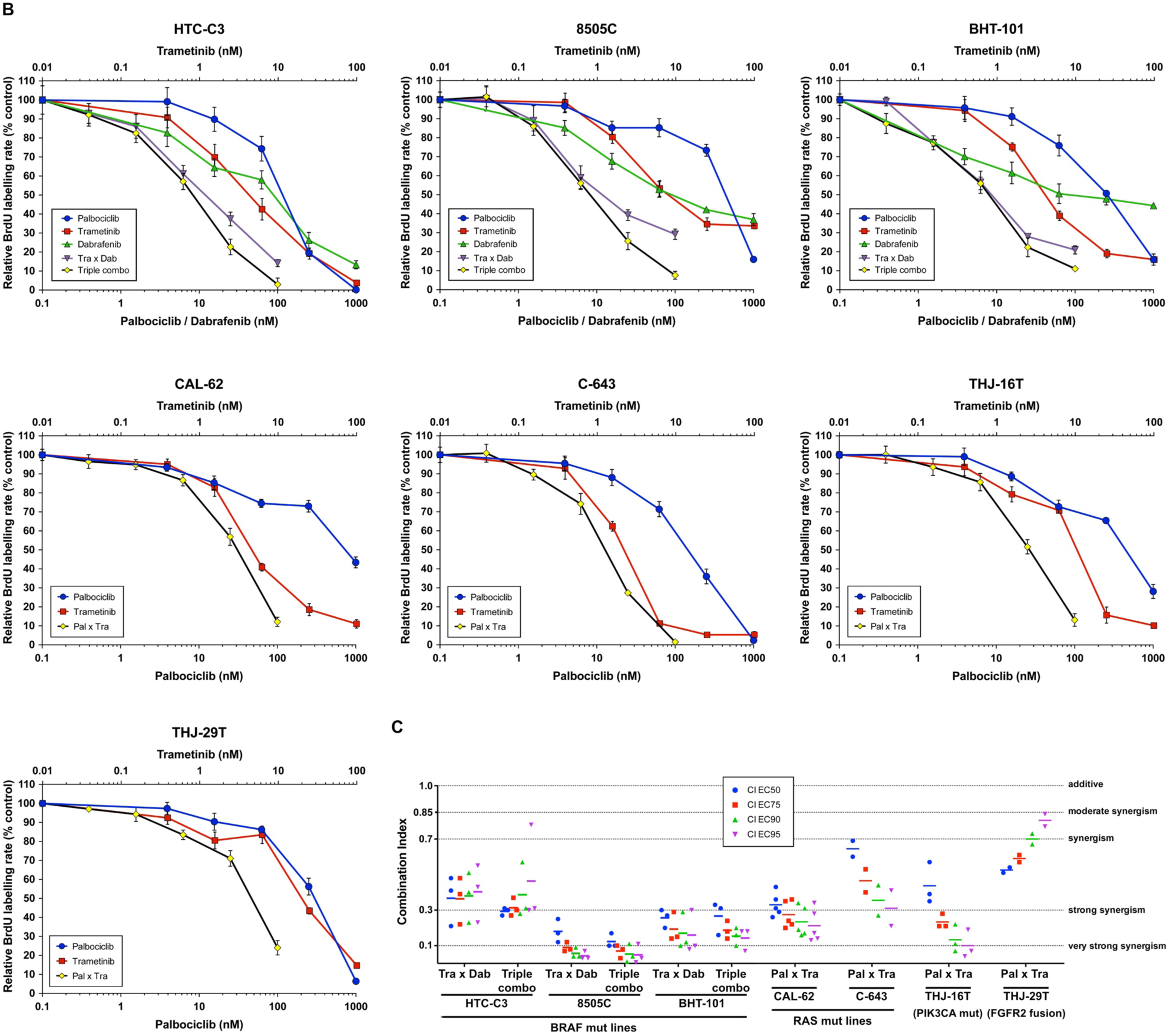
Combination of CDK4/6 inhibitors with MEK/BRAF inhibition potentiates the growth-inhibitory effects. (A) Indicated proteins were immunodetected after SDS-PAGE of total protein extracts from cells treated for 4 d with vehicle or with 1 µM palbociclib (Cyclin D1 detection as in Fig. 4A). Hyper– and hypo-phosphorylated forms of p70S6K1 are indicated. Cell lines defined as resistant, with residual proliferation or sensitive (as detailed in Fig. 3) are indicated by red, orange or black font colors, respectively. (B) Response curves (measured as BrdU incorporation during a 1 h pulse) in ATC-derived cell lines, following 24 h treatment with serial dilutions of dabrafenib (BRAF inhibitor), trametinib (MEK inhibitor) or palbociclib either alone, in combination of two (Tra x Dab; Pal x Tra) or in combination of three drugs (triple combo). Representative data of two or more independent experiments as shown in panel C (error bars: mean ± SD of triplicates). (C) Drug combination indexes (CI) calculated at different effective concentrations (EC) in the tested ATC– derived cell lines. Values obtained in each independent experiment are presented (line at mean).

To evaluate the long-term effect of the combinations, we performed clonogenic assays at two different concentration schemes, in two RAS– and in two BRAF-mutated cell lines (Fig. 6A). Tested concentrations were chosen in accordance to the concentration-response curves for each of the drugs and to known pharmacodynamics data (100-200 nM dabrafenib [71]; 15-30 nM trametinib [72,73]; 200-260 nM palbociclib [74]). Hence, higher palbociclib concentration was tested (1µM instead of 500 nM) for the most palbociclib-partially resistant line CAL-62, and trametinib concentration was unchanged between the two treatment schemes for cell lines with BRAF V600E mutation (highly effective at 5 nM). Cell lines treated for 10 d had their ability to form colonies reduced by each of the drugs alone (for cell lines with mutant RAS, this was seen only with higher concentrations). However, when the compounds were removed, there was a complete or almost complete regrowth (except for HTC-C3 line, which was more sensitive to palbociclib and trametinib than other cell lines). In combined treatments (either trametinib with dabrafenib, trametinib with palbociclib or the three drugs), clonogenic potentials of the cell lines were drastically reduced during the continuous presence of drugs (Fig. 6A). Importantly, following drugs withdrawal, only regimens combining palbociclib to BRAF and/or MEK inhibitors showed the most efficient and durable responses, completely abolishing the regrowth of most cell lines. For RAS mutant lines, this effect was observed only with higher concentration of drugs, while the triple combination was already effective at lower concentrations in BRAF mutant lines. The same effects were also seen for cell lines possessing other oncogenes (Supplementary Fig. S10). We observed that SA-β-gal activity was higher following combined treatment than with each of the drugs alone (Fig. 6B), whereas neither cleaved caspase 3 nor cleaved PARP1 were changed (data not shown). Thus, inhibition of ERK pathway synergizes with CDK4/6i, potentiating the proliferation-suppressive properties of each of the drugs and strongly impairing the long-term growth of thyroid cancer cells and the development of resistance.

**Figure 6.**
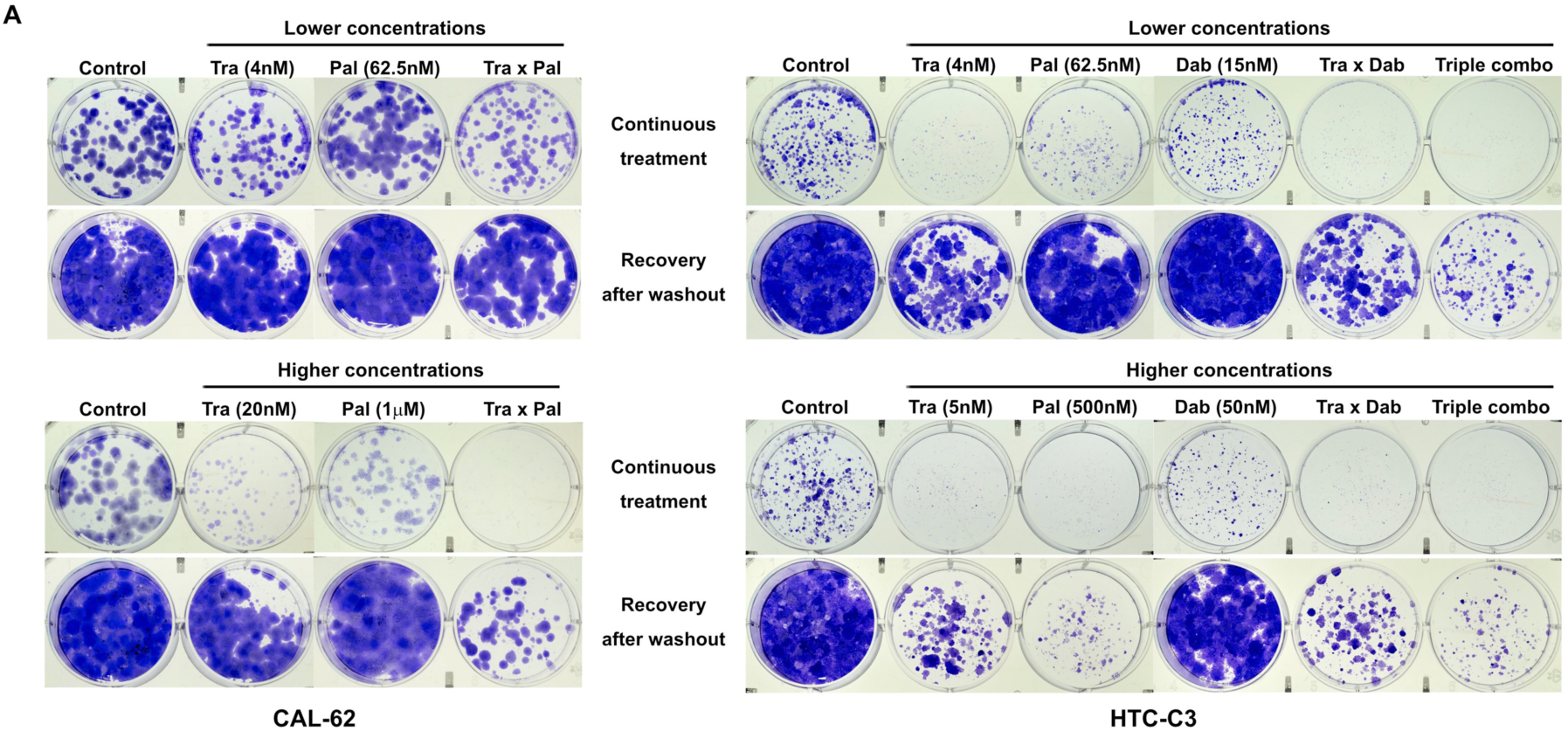
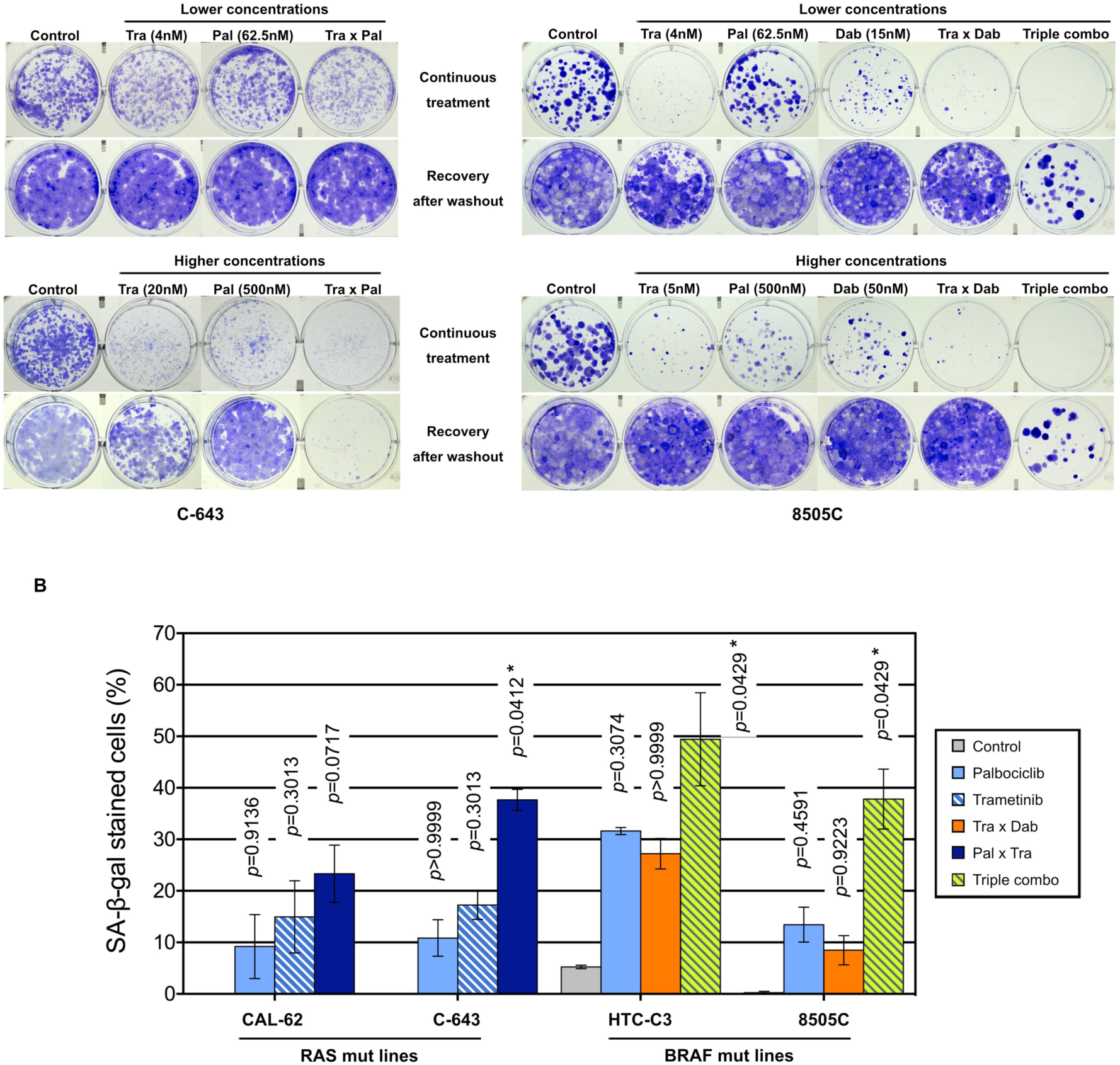
Combination of CDK4/6 inhibitors with MEK/BRAF inhibition prevents the clonogenic ability of thyroid cancer cells. (A) Clonogenic potentials of ATC-derived cell lines treated for 10 d at indicated concentrations with vehicle, dabrafenib (dab, BRAF inhibitor), trametinib (tra, MEK inhibitor) or palbociclib (pal) either alone, in combination of two or in combination of three drugs (triple combo), followed or not by a 10 d recovery period (drugs withdrawal). Representative photographic images of two independent experiments. (B) Cells stained for SA-β-gal activity in ATC-derived cell lines were quantified following treatment for 8 d (higher concentrations of drugs were used as indicated in panel A). Pooled data from two independent experiments (error bars: mean ± SEM). Statistical significance between control and each treatment was calculated with the Kruskal-Wallis test corrected by Dunn’s multiple comparison tests. *p<0.05.

### 3.6 Palbociclib treatment induces a paradoxical stabilization of activated Cyclin D– CDK4 complexes, which can be prevented by MEK/BRAF inhibitors

We next explored the molecular mechanisms behind the synergistic inhibition of cell cycle progression and cell proliferation in response to combined targeting of CDK4/6 and MEK/ERK pathways. We observed that a 3 d treatment with the combination of the drugs at low concentrations, similar to the ones used for clonogenic assays (Fig. 6A), already allowed to efficiently arrest the cell proliferation by repressing RB phosphorylation and Cyclin A2 expression (Fig. 7A). In addition, the combination with trametinib and dabrafenib prevented the upregulation of both Cyclins D and Cyclin E1, as well as the activation of ERK and mTOR pathways (as inferred by the decreased phosphorylation levels of ERK, p70S6K1, RPS6 and 4EB-P1), that were observed upon palbociclib treatment (Fig. 7A). These effects were also maintained or further enhanced when 250 nM of palbociclib was used instead of 62.5 nM (Supplementary Fig. S11).

**Figure 7.**
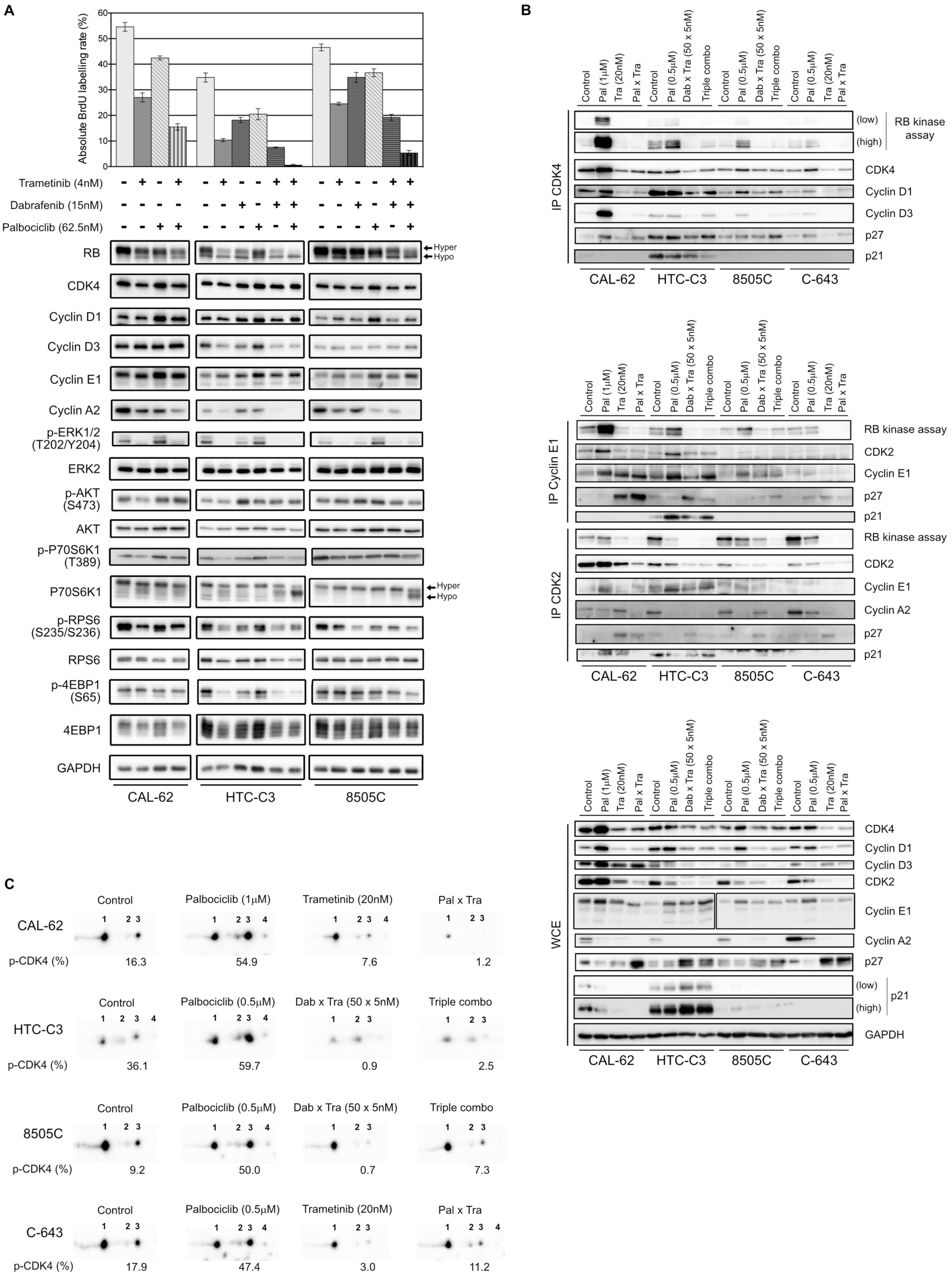
Palbociclib treatment induces a paradoxical stabilization of active Cyclin D-CDK4 complexes, which can be prevented by MEK/BRAF inhibitors. (A) BrdU incorporation rate (during a 1 h pulse) following 24 h treatment with vehicle or with indicated drugs. Error bars: mean ± SD (of triplicates; n = 1). Indicated proteins were immunodetected after SDS-PAGE of total protein extracts from cells treated for 3 d with vehicle or with indicated drugs. Hyper– and hypo– phosphorylated forms of RB and p70S6K1 are indicated. (B) Co-immunoprecipitation assays (IP) with an anti-CDK4 (upper panel), anti-Cyclin E1 or anti-CDK2 antibody (middle panel) followed by *in vitro* RB-kinase assay and immunodetection of indicated proteins, from cells treated for 6 d with vehicle, dabrafenib (dab, BRAF inhibitor), trametinib (tra, MEK inhibitor) or palbociclib (pal) either alone, in combination of two or of three drugs (triple combo). RB-kinase activity was evaluated by the immunodetection of the *in vitro* phosphorylated RB fragment. Immunodetection of the indicated proteins were also done after SDS-PAGE of whole-cell extracts (WCE) from cells treated in the same conditions (lower panel). Low: low exposure time; high: high exposure time. (C) Immunodetection of CDK4 after 2D-gel electrophoresis separation of whole protein extracts from cells treated as in panel B. Below each detection is shown the ratio of T172-phosphorylated form of CDK4 (spot 3) over the total of detected CDK4 forms (native form, another modified form, T172-phosphorylated form and double modified form labelled as spot 1, spot 2, spot 3 and spot 4, respectively), quantified from the immunoblots.

We then evaluated the effect of the combined treatments on the formation and specific kinase activities of CDK4 complexes, by performing co-immunoprecipitation assays from cells treated for 6 d (Fig. 7B, upper panel), with the high concentration scheme of drugs tested in the clonogenic assays (Fig. 6A). The *in vitro* RB-kinase activity of CDK4 complexes (assayed in the absence of palbociclib, hence reflecting the presence of activated CDK4) was increased by palbociclib mostly in CAL-62 cells (the more palbociclib-resistant cell line) and to a lesser extend in HTC-C3, 8505C and C-643 cell lines. This increase was prevented by the co-treatment with trametinib in CAL-62 cells or by the dabrafenib/trametinib combination in the BRAF-mutated cells (Fig. 7B, upper panel). In CAL-62 (and more weakly in 8505C cells), the dramatic increase of RB-kinase activity of CDK4 induced by palbociclib was associated with a similarly increased formation of cyclin D3-CDK4 complexes, which were devoid of p21 and thus did not require the p21 assembly activity. This observation exactly corresponds to our previous description of a paradoxical effect of palbociclib that specifically stabilized activated cyclin D3-CDK4/6 complexes, which could become hyperactive upon drug removal, because their activity was not restricted by binding to CIP/KIP inhibitory proteins [52]. Similar to what we observed in Fig. 4A, palbociclib-treated cells exhibited increased levels of either CDK4, Cyclin D3 and/or Cyclin D1, and reduced levels of p27 (Fig. 7B, lower panel). Conversely, trametinib and the dabrafenib/trametinib combination reduced the protein concentrations of CDK4, Cyclin D1 and Cyclin D3, while increasing p27 accumulation (Fig. 7B, lower panel). The p27 modulation is in agreement with the already described reduction of p27 levels upon expression of RAS, BRAF or RET/PTC mutants in thyroid cancer cells, which would be reversed by MEK inhibition [75]. These effects, which were variably observed in the different cell lines, might also concur (respectively) to increase the formation of activated CDK4 complexes in response to palbociclib, and to explain the inhibition of this formation by BRAF/MEK inhibitors (Fig. 7B, upper panel). To further characterize the effect of the drugs on CDK4 activity, whole protein extracts from treated cell lines were resolved by 2D-gel electrophoresis (Fig. 7C). Treatment with palbociclib increased the presence of all the CDK4 forms, and even more the proportion of the activated T172-phosphorylated form. Again, combination with MEK/BRAF inhibitors was able to counteract these effects, consistently with our previous observation that CDK4 phosphorylation depends on MEK activity [24].

In the same experiments, we similarly evaluated the effects of the combined treatments on the formation and RB-kinase activity of CDK2 complexes that were (co-)immunoprecipitated using either CDK2 or Cyclin E1 antibodies (Fig. 7B, middle panel). Intriguingly, the RB– kinase activity associated with Cyclin E1 complexes also increased in response to palbociclib treatment in CAL-62, HTC-C3 and 8505C cells, mirroring the situation observed in CDK4 complexes. Again, this paradoxical effect of palbociclib was prevented by BRAF/MEK inhibitors (Fig. 7B, middle panel). These effects were (respectively) associated with an increased abundance of Cyclin E1 and Cyclin E1-CDK2 complex in palbociclib-treated cells, and a reduction of the binding of CDK2 to Cyclin E1 together with an increased association of p27, in cells treated with MEK or MEK/BRAF inhibitors.

At variance with the situation observed in Cyclin E1 co-immunoprecipitates, the total RB–kinase activity associated with CDK2 complexes (immunoprecipitated using a CDK2 antibody, and thus comprising both Cyclin E1 and Cyclin A2 complexes) was not increased by palbociclib in CAL-62 cells and was reduced in the three other cell lines with the most complete inhibition in HTC-C3 (the most palbociclib-sensitive cells) (Fig. 7B, middle panel). In all the cell lines, accumulation of Cyclin A2 and its association with CDK2 were reduced in palbociclib-treated cells, compensating for the increased presence and activity of Cyclin E1-CDK2 complexes. Intriguingly, treatments with trametinib and with dabrafenib/trametinib used alone (respectively in CAL-62 and in 8505C), reduced Cyclin A2 expression (Fig. 7B, lower panel) but not its presence in CDK2 complexes (Fig. 7B, middle panel). Nevertheless, the most complete inhibitions of both CDK2 activity and CDK2-Cyclin A2 association were observed in cells treated with the combination of palbociclib and MEK/BRAF inhibitors (Fig.7B, middle panel).

To summarize, the upregulation of Cyclins D with stabilization of activated (T172-phosphorylated) CDK4 complexes and the upregulation of Cyclin E1 with increase of its kinase activity are most probably the mechanisms that bypass CDK4/6 inhibition in thyroid cancer cells. Nonetheless, this can be prevented through combined MAPK/ERK inhibition, which reduces the accumulation of Cyclins D and Cyclin E1, the phosphorylation of CDK4, and increases p27 to further inhibit CDK2 activity.

## 4. Discussion

CDK4/6 inhibitors have been proposed as an important tumor therapy for which there should be the fewest bypass mechanisms [76]. It is generally considered to be reserved to RB-proficient tumors, because intrinsic resistance to CDK4/6i is mostly associated to RB loss. However, aberrant activation of E2F factors, Cyclin E or CDK2 can also circumvent CDK4/6 inhibition and may be involved in either intrinsic or adaptive resistance. One of the novelties of our work comes from our ability to detect in tumors the presence of the main target of CDK4/6i, i.e. T172-phosphorylated active CDK4. This phosphorylation, which is required for the opening of the catalytic site of CDK4 [77,78], actually represents the rate-limiting step of CDK4 activation because, contrary to other CDKs, like CDK2 and CDK1 [79,80], CDK4 activity is not limited by inhibitory phosphorylations [13,81,82]; and because the T172 phosphorylation is finely regulated, while depending on all the preceding steps, including binding to a cyclin D [11]. Here, we interrogated, for the first time, the presence of T172-phosphorylated CDK4 in the different sub-types of thyroid cancer. Our results extend our demonstration in breast tumors and for pleural mesotheliomas: CDK4 phosphorylation was absent or very weak in normal quiescent thyroid tissue, but detectable in most thyroid tumors and derived cell lines. Nevertheless, its absence in tumors (despite active proliferation) was also observed, which correlated with insensitivity to CDK4/6i in cell lines and with known resistance markers to these drugs. In our transcriptome analyses, in one of the biggest cohorts of ATC and PDTC samples studied by RNA-seq so far, we showed that the absence of phosphorylated CDK4 (profile A) is mainly found in ATC (30%, n=7) and in one exceptional PDTC case (5%). This is associated with lower *RB1* expression and higher *CCNE1* and *E2F1* expressions. Still, *RB1* loss or mutation was not observed in several profile A cases and aberrations of *E2F1* or *CCNE1* were not found. In one ATC (JPI21) with no detectable *RB1* alteration, *CCNE1* was highly expressed, possibly as a result of *E2F1* amplification (Supplementary Fig. S3). Therefore, although relatively minor, the prevalence of ATC that should be intrinsically resistant to CDK4/6i might well exceed the low mutation rate of *RB1* gene reported in ATC [5], indicating that the presence or absence of phosphorylated CDK4 could be the most relevant biomarker to predict CDK4/6i sensitivity.

The biochemical detection of T172-phosphorylated CDK4 was dependent upon the availability of fresh-frozen tissue samples. Moreover, detection of CDK4 phosphorylation by IHC in FFPE samples may be precluded by the low abundance of the phosphorylated CDK4 form [54] and the likely loss of phosphorylation before and/or during formalin fixation. Surrogate biomarkers including signatures constructed from gene expression analysis like RNA-seq, may instead be used to predict the CDK4 phosphorylation status [26]. Moreover, like most profile A cases in breast cancers and mesotheliomas, the ATC and PDTC cases lacking CDK4 phosphorylation were associated with over-expression of p16 mRNA and protein. Nevertheless, high *CDKN2A* mRNA levels were also found in some profiles H and L thyroid tumors (associated with CDK4 phosphorylation). In depth analyses of these cases revealed that the *CDKN2A* expression was restricted to exons encoding the alternative protein p14 (instead of p16) or was encoding a mutated (truncated) p16 protein. An immunohistochemistry assessment showed that most profile A tumors exhibited strong p16 expression. Still the elevation of p16 staining should be carefully appreciated. Intriguingly, the expression of p16 is very low in normal thyroid tissues but often moderately elevated in different thyroid tumors, including PTC and FTC [58,60,61,83–85]. Possibly, this reflects a p16-dependent senescence mechanism limiting oncogene-driven proliferation [86–88] and thus, it would be important to characterize the proliferative state of these p16-positive cells [28]. In contrast, in profile A ATC and PDTC cases, the expression and immunostaining of p16 were much higher and were associated with a proliferation marker such as KI67. Therefore, appropriately scored p16 IHC (especially regarding intensity and homogeneity of expression) associated with positive KI67 detection might identify most of the profile A cases. Worth mentioning is that in one case, p16 elevated expression seemed to be a subclonal event – the protein was highly expressed but not present in all tumor regions. Our results show that the CDK4 modification profile of thyroid tumors and cell lines can also be predicted using the expression values of the same 11 genes used to predict the CDK4 modification profiles of breast tumors [26], reaching accuracies of 98.2% in tumors and 90.5% in cell lines, with a binary A/nonA classification. Such a binary classification of thyroid tumors, which distinguishes tumors with intrinsic resistance to CDK4/6i from tumors expressing the drug target, would meet the clinical need. Once adapted to the use of RNA extracted from FFPE tissue (manuscript in preparation for the breast scenario), our prediction tool may directly help to guide the treatment of advanced thyroid cancer. It will complement the IHC evaluation of the p16 and KI67 expression by identifying the potential false positives defined above. On the other hand, the IHC evaluation of the p16 and KI67 expression will identify tumors with exclusive p14 contribution to the expression of the locus and tumors with high *CDKN2A* expression due to frameshift or stop mutation, which are the main sources of false positives in the prediction test.

Predicting the intrinsic insensitivity of some ATC to CDK4/6i might be important because, like RB-deficient tumors in other cancers such as small cell lung cancers (SCLC), they might respond particularly well to genotoxic chemotherapy [89,90]. A larger number of cases and longer follow-up should indeed determine whether the remission of an ATC patient (JPI21), who is still alive more than 8 years after diagnosis and chemotherapy, is really exceptional or holds hope for similar cases. Among two other patients with profile A and/or high p16 detection who also received chemotherapy, one was still alive at last follow-up (PDTC JPI25), and the other is still followed currently (ATC JPI84), more than 8 months after diagnosis (Supplementary Table S3). Interestingly, two exceptional cases of advanced ATC with complete pathological response after radio/chemotherapy have been molecularly characterized and one of them was associated to a *RB1* frameshift mutation [91]. More clinical cases are needed to conclude whether the mechanisms underlying p16 over-expression and lack of active CDK4 might promote a better response to chemotherapy. In addition, transient administration of CDK4/6i in tumors predicted to be insensitive could be used to temporarily arrest proliferation of normal cells and prevent chemotherapy-induced side-effects (such as myelosuppression), hence allowing increase of the dose of genotoxic drugs [92]. This has been validated for SCLC [93], which led to the recent approval of trilaciclib for this indication by the FDA.

On the other hand, p16 deficiency (either by mutation, promoter methylation or deletion) has been associated with more aggressive cases of thyroid cancer [55,59,94–96]. This includes a lower thyroid differentiation in ATC and significant association with increased mortality in patients with metastatic PTC, PDTC and ATC [55]. Analogously, despite the limited number of cases, we observed in our study that tumors with higher CDK4 activation (CDK4 profiles H versus profiles L) tend to have lower differentiation in PTC, higher proliferative potential in PDTC and ATC, higher incidence of metastases in PTC and PDTC and a shorter overall survival for PDTC and ATC patients. Nevertheless, in PTC and in a subset of PDTC, the H profile of CDK4 was not associated with a higher proliferative activity, in a similar manner as reported for the over-expression of Cyclin D1 in PTC [97,98]. Likely, in more differentiated tumors, inhibitory mechanisms involving p53 and higher expression of p21 (*CDKN1A*) or p27 (*CDKN1B*), which also can restrict the activity of CDK4 complexes without impeding T172 phosphorylation [13,22,36,53], and also impair CDK2 activity, might still limit the cell cycle progression.

Overall, these observations support the potentially favorable clinical impact of inhibiting CDK4 activation in some thyroid tumors. In cases presenting T172-phosphorylated CDK4 (excluding the rare possibility of p16 loss co-occurring with RB deficiency), tumors are expected to depend on the CDK4 activity and CDK4/6i might represent a relevant therapy when required. We indeed observed that palbociclib and abemaciclib strongly inhibit cell cycle progression and proliferation in most thyroid cancer cell lines, including all ATC and PDTC-derived ones. Our large collection of cell lines from different tumor histotypes further completes previous works on the evaluation of CDK4/6i [66–68,99]. The inhibitory concentrations calculated in the present study were in agreement with the ones analyzed in the collection of ATC cell lines from Wong *et al*., [66], although we did not observe any tendency for differential sensitivity according to mutational status. Nevertheless, we confirmed that sensitivity to palbociclib was negatively correlated with *CDK4* mRNA levels (but not with the levels of *CCND1* nor *CDK6*) and we also evidenced a similar negative correlation with *CCNE1* expression.

Despite initial response, the cell cycle blockade by palbociclib was bypassed over time in several thyroid tumor cell lines, as also reported by Wong *et al*. [66] and in other cancers, including pancreatic ductal carcinomas [100]. As in these other cancer models, this adaptation to CDK4/6 inhibition was associated to the induction of Cyclins –D1, –D3 and –E1 [66,100]. Particularly, the upregulation of Cyclin E1 resulted in much augmented interaction of Cyclin E1 with CDK2 and associated RB-kinase activity (Fig. 7B). In addition, we identified here a novel mechanism that involves a strong increase of CDK4 T172 phosphorylation and the stabilization of activated CDK4 complexes in response to palbociclib (Fig. 7B,C). Therefore, the direct inhibition of CDK4/6 could be compensated by various adaptive responses involving both a dramatic augmentation of activated CDK4 complexes and the distal activation of Cyclin E1-CDK2. Irrespective of the mechanism, the bypass or adaptation to CDK4/6 pharmacological blockade imposes the use of combinatorial therapy [101]. In breast cancer, the combination of CDK4/6i with endocrine therapy became standard procedure and is certainly essential to restrict adaptive events, via reduction of Cyclin D1 levels and Cyclin E1 activity [102]. A PI3K/mTOR dual inhibitor was previously used to potentiate the palbociclib action in an ATC xenograft model [66] but due to lethality, doses had to be reduced. In the present study we combined CDK4/6i with anti-MEK/BRAF agents. Indeed, we observed here that the upregulation of cyclins D and T172-phosphorylated CDK4 during palbociclib treatment, for the most part, paralleled an increased activity of MAPK/ERK pathway [101]. This ERK pathway activation was critical because all the adverse responses associated with the adaptation to CDK4/6 inhibition (including upregulation of cyclins D and E and accumulation of activated CDK4 and cyclin E1 complexes) were prevented by the combination of palbociclib with dabrafenib and trametinib in BRAF-mutated cell lines and with trametinib in the BRAF-wild type cell lines. This allowed a sustained, and largely irreversible, control of cell proliferation in response to the combined treatments, with lower concentrations of palbociclib and of the BRAF and/or MEK inhibitors, compared to single agent therapy. The combination therapy of dabrafenib with trametinib was beneficial in a relevant proportion of locally advanced or metastatic ATC with *BRAF* V600E mutation [70,103], becoming the first ever regimen approved for ATC patients by the FDA. While recommended [104] and showing clear benefits over standard care also in a European real-world setting [105], this targeted therapy is not yet approved by European Medical Association (EMA). Treatment with dabrafenib plus trametinib or in combination with immunotherapy (pembrolizumab) is presently considered a useful strategy for pursuing ATC patient stabilization or slowing disease progression, as well as in the neo-adjuvant setting, with complete surgical resection achieved in some cases [9,105–108]. Indeed, the clinical benefit of these options may still be impacted by the frequent high-grade adverse events and by the emergence of acquired resistance. In *BRAF* V600E-mutated patient-derived melanoma xenografts, addition of palbociclib to dabrafenib and trametinib combination was shown to drastically increase tumor regression, in both naïve and dabrafenib/trametinib-resistant tumors [109]. Interestingly, p16 loss appears to be positively selected following treatment with anti-BRAF agents [110]. Therefore, there could be a strong rationale for combining anti-BRAF therapy with CDK4/6 inhibition, especially in neo-adjuvant settings, to improve the probability of effective tumor surgical resection.

According to NGS analysis, *BRAF* V600E mutation are found in only 13 to 44% of ATC cases [5]. Here, we provided evidence that the combination of trametinib with CDK4/6i could potentially represent a therapeutic option in the relevant proportion of wild type BRAF-ATC. Anti-CDK4/6 drugs have been shown to impact different cell types in the tumor microenvironment and enhance tumor immunogenicity [111–115]. Combination with trametinib was also shown to be effective in different cancer types and to be able to boost immune anti-tumoral activity [101,116,117]. RAS mutations are known to confer greater sensitivity to CDK4/6i and CDK4 was found to be critical for oncogenic NRAS and KRAS signaling [118–120]. Thus, the use of this drug combination most likely will be beneficial for the treatment of both PDTC and RAI-refractory differentiated thyroid cancers. Currently, only three multi-targeted tyrosine kinase inhibitors (sorafenib, lenvatinib and cabozantinib) are approved with this indication and are restricted to patients presenting symptomatic, multi-metastatic, rapidly progressive disease [8,121]. Unfortunately, the impact of these drugs on patient survival is rather limited, with resistance and disease progression occurring after a few months [8,121,122]. Moreover, the patients are exposed to numerous serious side effects, often requiring dose reduction or discontinuation of the treatment [122]. Thus, the association trametinib-CDK4/6i could actually represent a therapeutic option also as second/third line therapy in such patients.

To conclude, our study supports the potentially favorable clinical impact of CDK4/6i for the treatment of aggressive dedifferentiated thyroid tumors including in combinations with anti-BRAF/MEK and, likely, anti-ERK therapies. Most ATC and PDTC patients could at least initially respond to CDK4/6i, which may arrest tumor growth, facilitating surgery in the neo-adjuvant setting or improving the efficacy of other treatment option. Future clinical trials are warranted to determine dose limiting toxicities, maximum tolerated dose and efficacy. Eventually, the prompt identification of tumors lacking CDK4 phosphorylation could represent a useful tool for recognizing the intrinsically CDK4/6i insensitive patients as potentially better candidates to immediate chemotherapy. This can be achieved by the complementary use of the 11 genes expression-based tool with p16 and KI67 IHC evaluation.

## Data accessibility

The RNA-sequencing data from patient-derived cell lines and from thyroid tumor patients are being deposited at the European Genome-phenome Archive (EGA), which is hosted by the EBI and the CRG.

The RNA-sequencing data from commercial cell lines are being deposited in the Gene Expression Omnibus.

## Author contributions

PPR and ER conceived the project; JMP, KC and PPR designed and performed experiments; JMP, ER, KC, JED, GC and PPR analyzed and discussed the data; ER, MT and FL performed bioinformatics analyses; MD-P and EL performed the histopathological analyses; LAM and JAC provided patient-derived cell lines; MD-P, GD, LC, GA, LW, EL, CT, LAM, JAC, CD, CM and BMC provided samples and related data; JMP, ER, GC and PPR wrote the original draft. All the authors contributed to manuscript editing and approved the final version.

## Supporting information

Data Set 1

## Acknowledgments

We thank Sabine Paternot for support, discussion and critical reading of the manuscript; Vincent Vercruysse for technical assistance; The Pathology Department from Institut Jules Bordet, particularly Alex Spinette and Dr. Nicolas de Saint Aubain for the assistance; Margarida M. Moura and Carolina Pires from Unidade de Investigação em Patobiologia Molecular from IPOLFG for tumor sample selection and preparation; Teresa Pereira, Marta Mesquita and Rafael Cabrera, from the Pathology Department from IPOLFG, for immunohistochemical and histopathological analysis; Dr. Valeriano Leite, from IPOLFG, for support and revision on patients’ clinical data; Tumor banks from Centre de Ressources Biologiques (CRB) des Hospices Civils de Lyon (HCL) and Groupement de Coopération Sanitaire-Centre Régional de Référence en Cancérologie (C2RC) de Lille for the contribution and technical support on the samples; Annick Brandenburger, Laure Twyffels and Véronique Kruys for assistance on microscopic images acquisition; Manuel Saiselet and Adrien Tourneur for continued interest and support on samples collection/analysis; We also thank the Brussels Interuniversity Genomics High Throughput Core (www.brightcore.be) and Dr Anne Lefort for RNA handling and sequencing.

Pr. Jacques E. Dumont closely followed and supported this work. The authors dedicate this study to his memory.

## Supporting Information

Table S1. Description of cell lines.

Table S2. Description of antibodies.

Table S3. Tissue samples characteristics and associated clinical data.

Table S4. Total number of thyroid normal tissues and tumor subtypes samples (and number of cases of each CDK4 modification profile) analyzed by RNA-seq.

Table S5. Mutational screening of tumor samples analyzed by RNA-seq.

Table S6. Immunohistochemical analysis of p16 and KI67 in FFPE samples from ATC and PDTC tumors.

Table S7. Expression of the 11 genes signature in thyroid tumors.

Table S8. Prediction of the CDK4 modification profile in thyroid tumors.

Table S9. Expression of the 11 genes signature in thyroid cancer cell lines.

Table S10. Prediction of the CDK4 modification profile in thyroid cancer cell lines.

Table S11. Summary of relevant molecular features and responses to palbociclib treatment in the ATC cell lines tested for the combination of anti-CDK4/6, anti-BRAF and anti-MEK therapies.

Supplementary Figure S1. Immunodetections of CDK4 after 2D-gel electrophoresis separation of whole protein extracts from the fresh-frozen samples cohort.

Supplementary Figure S2. Representation of the gene expression variance between all 84 RNA-seq analyzed samples by principal component analysis.

Supplementary Figure S3. Genome-wide copy number profiles of profile A tumors.

Supplementary Figure S4. Immunohistochemical stainings for hematoxylin/eosin (HE), KI67 and p16 performed in serial sectioning of FFPE samples from ATC and PDTC tumors.

Supplementary Figure S5. CDK4 phosphorylation levels in thyroid tumors are associated to different molecular features.

Supplementary Figure S6. Comparison between BrdU incorporation, MTT and SRB assays.

Supplementary Figure S7. Correlation in thyroid cancer cell lines between palbociclib response and expression of putative markers of response.

Supplementary Figure S8.Treatment with palbociclib does not induce cell apoptosis/autophagy.

Supplementary Figure S9. Sensitivity of thyroid cancer cell lines to the combination of abemaciclib with MEK/BRAF inhibitors.

Supplementary Figure S10. Long-term effect of the combined CDK4/6 and MEK/BRAF inhibition in additional thyroid cancer cell lines.

Supplementary Figure S11. Impact of the CDK4/6, MEK and BRAF inhibitors on cell cycle– and signaling pathways-related protein levels.

Data Set 1. Complete list of gene expressions obtained from RNA-seq analysis of thyroid tissue samples (A) and thyroid cell lines (B).

