## Supplementary Figure 4 for "CDK4 phosphorylation status and rational use for combining CDK4/6 and BRAF/MEK inhibition in advanced thyroid carcinomas"

### ATC4 - JPI21 - Profile A

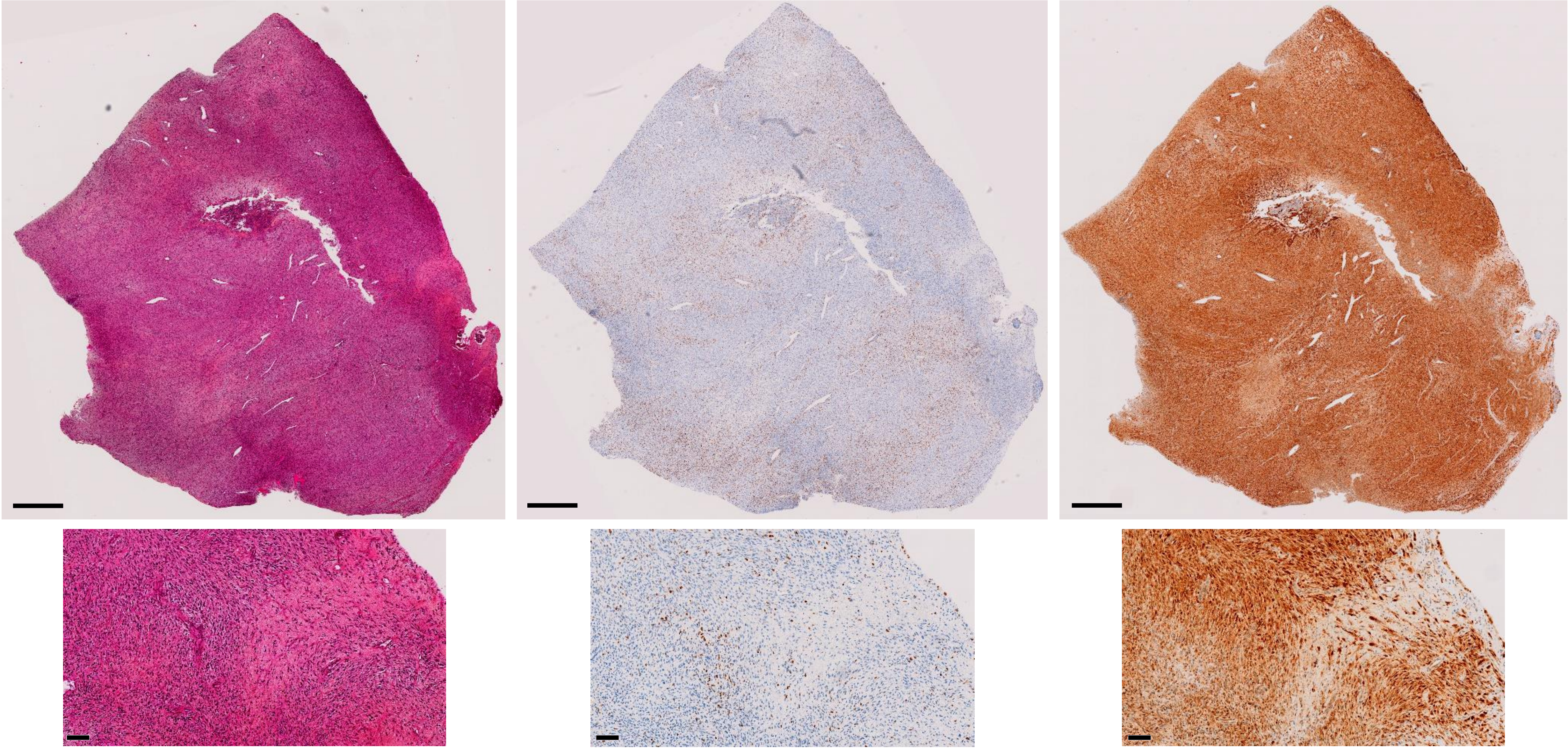

HE

KI67

p16

Scale bar = 1mm

Scale bar = 100µm

### ATC7 - JPI74 - Profile A

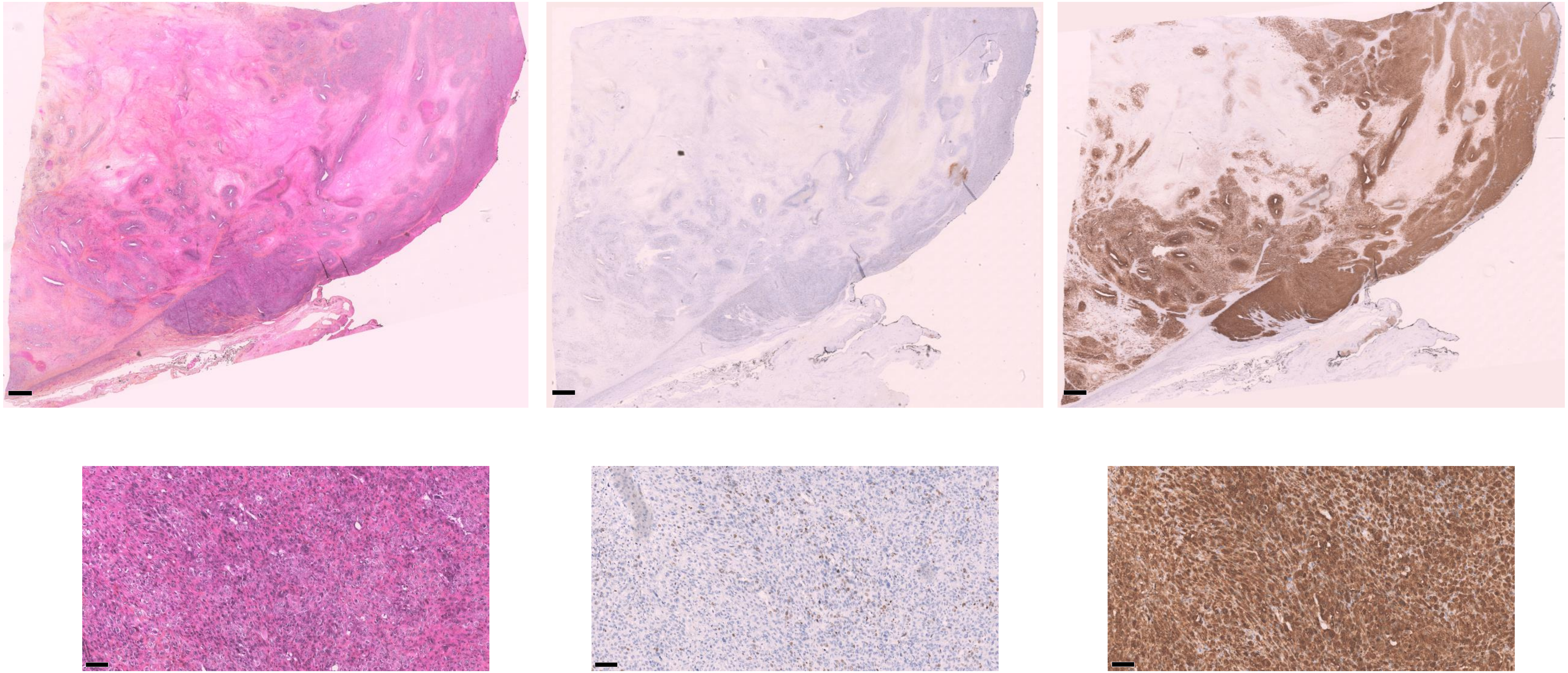

HE

KI67

p16

Scale bar = 1mm

Scale bar = 100µm

### ATC1 - Profile A

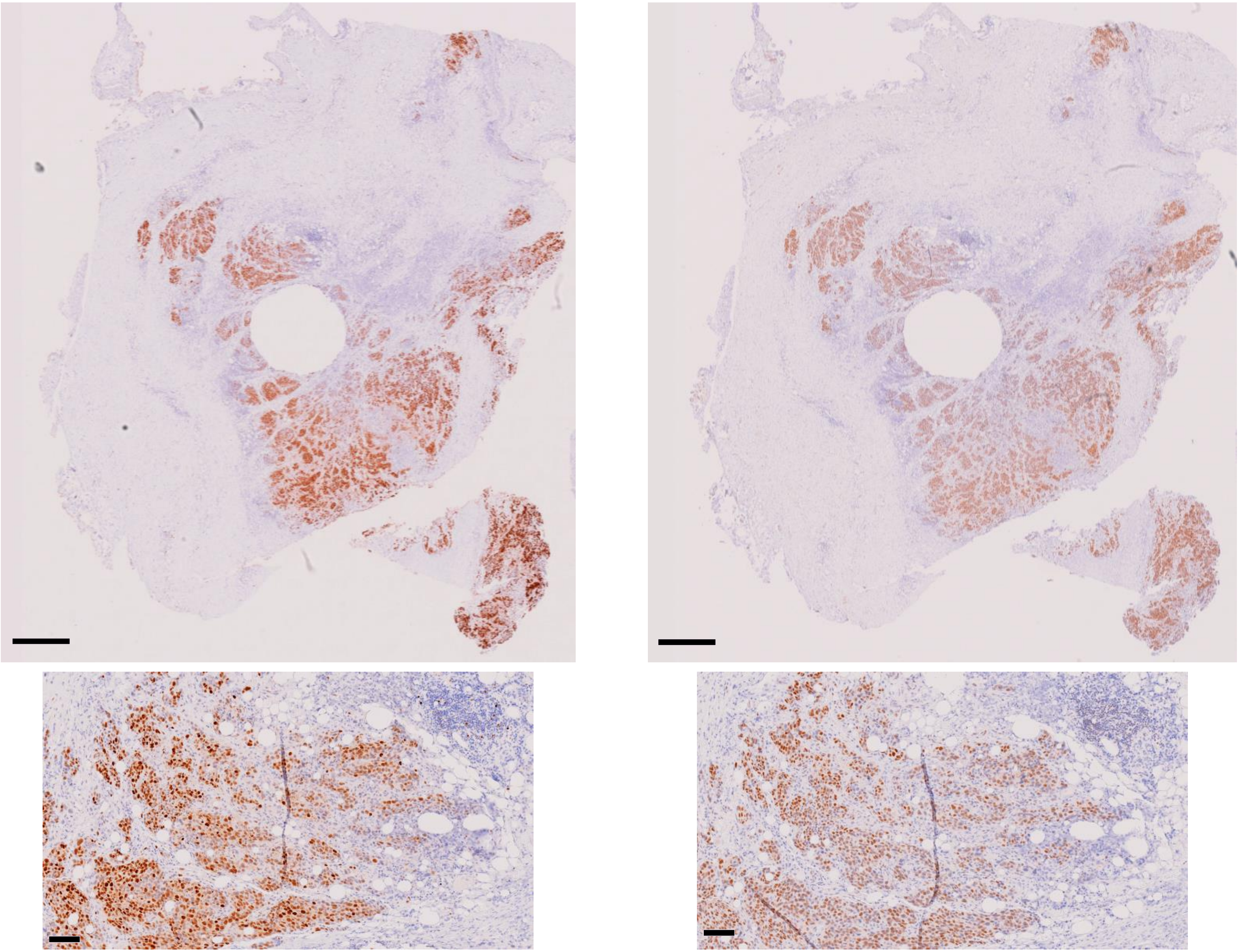

KI67

p16

Scale bar = 1mm

Scale bar = 100µm

### ATC2 - Profile A

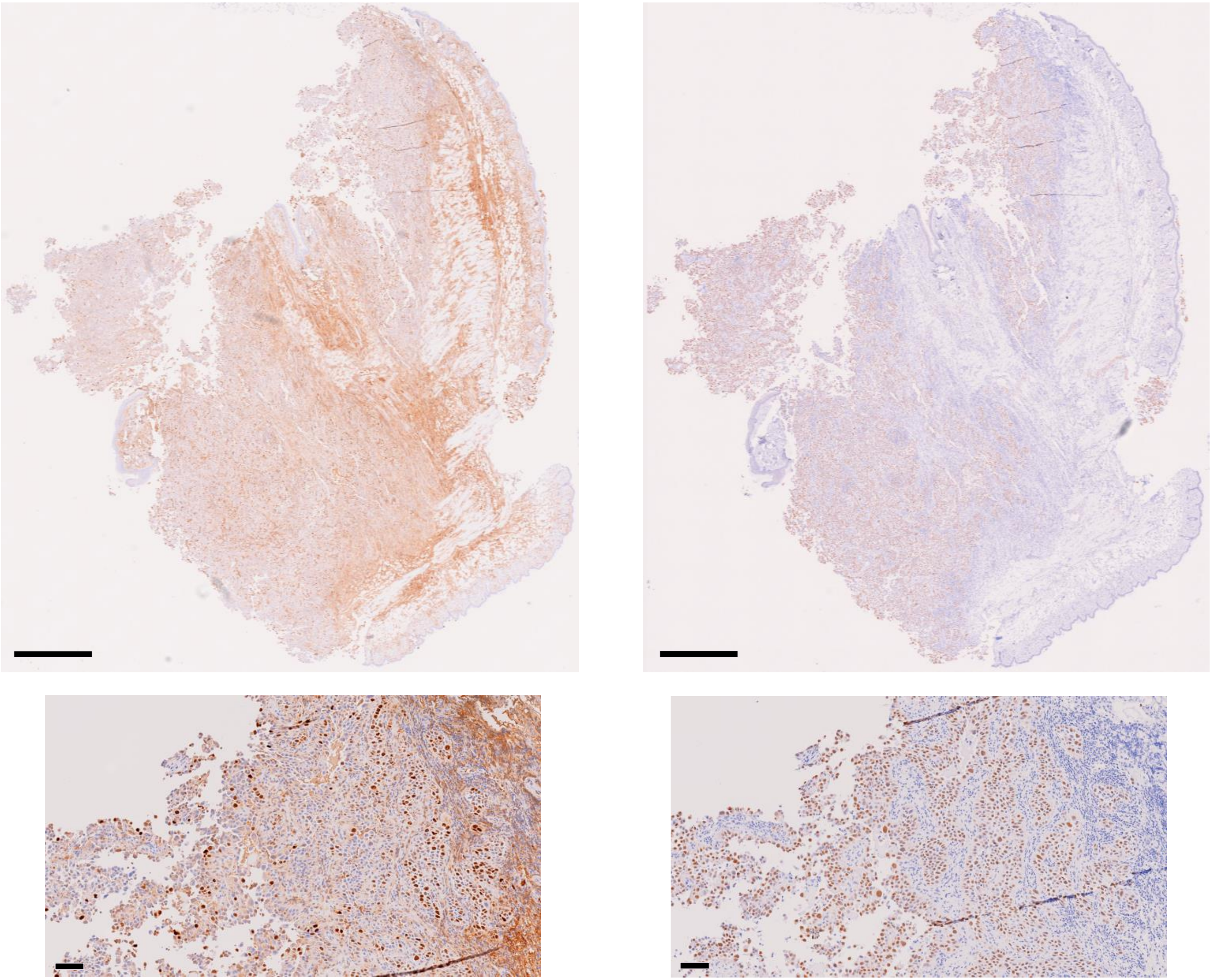


Scale bar = 1mm

Scale bar = 100µm

### ATC3 - Profile A

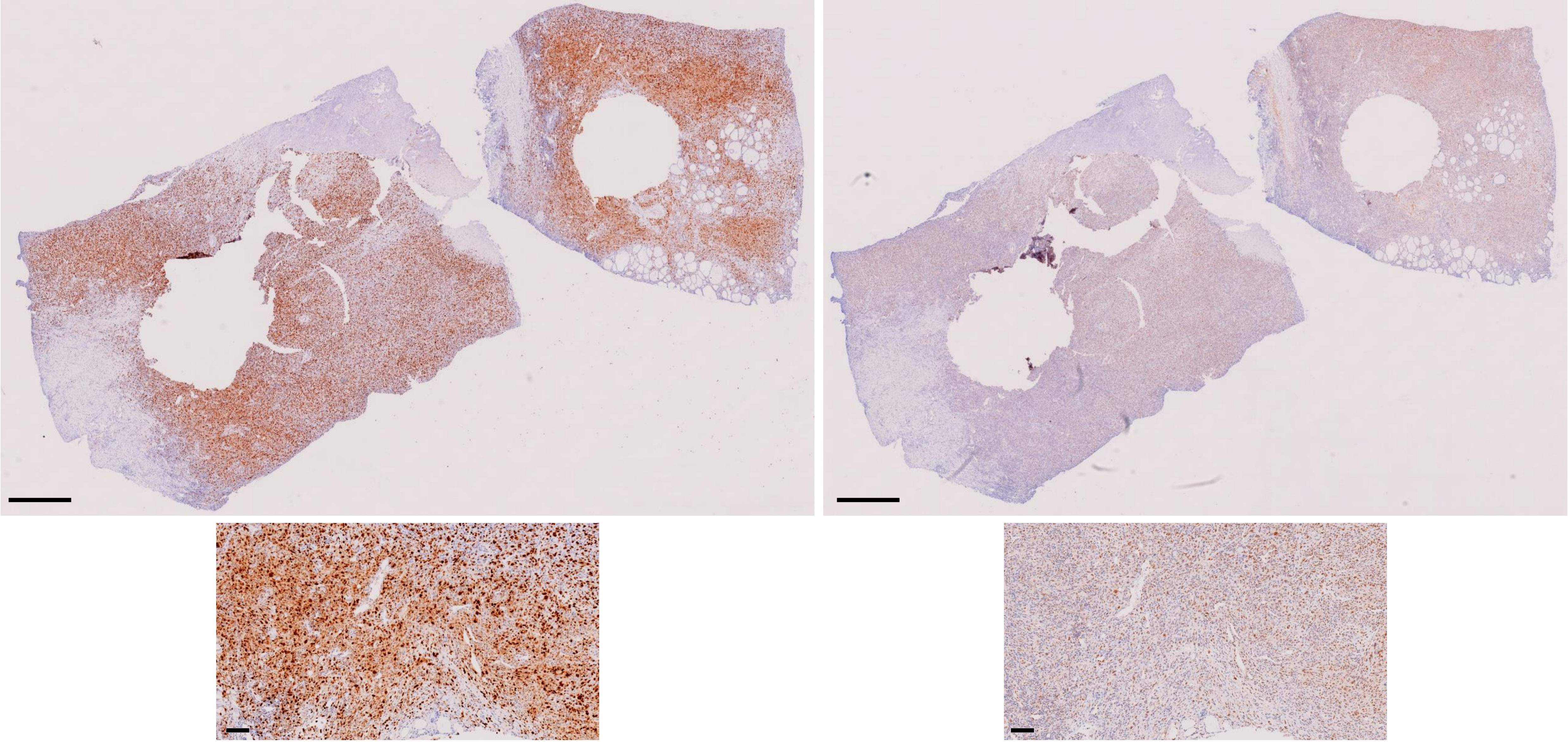

KI67

p16

Scale bar = 1mm

Scale bar = 100µm

### ATC12 - JPI84 - Profile A

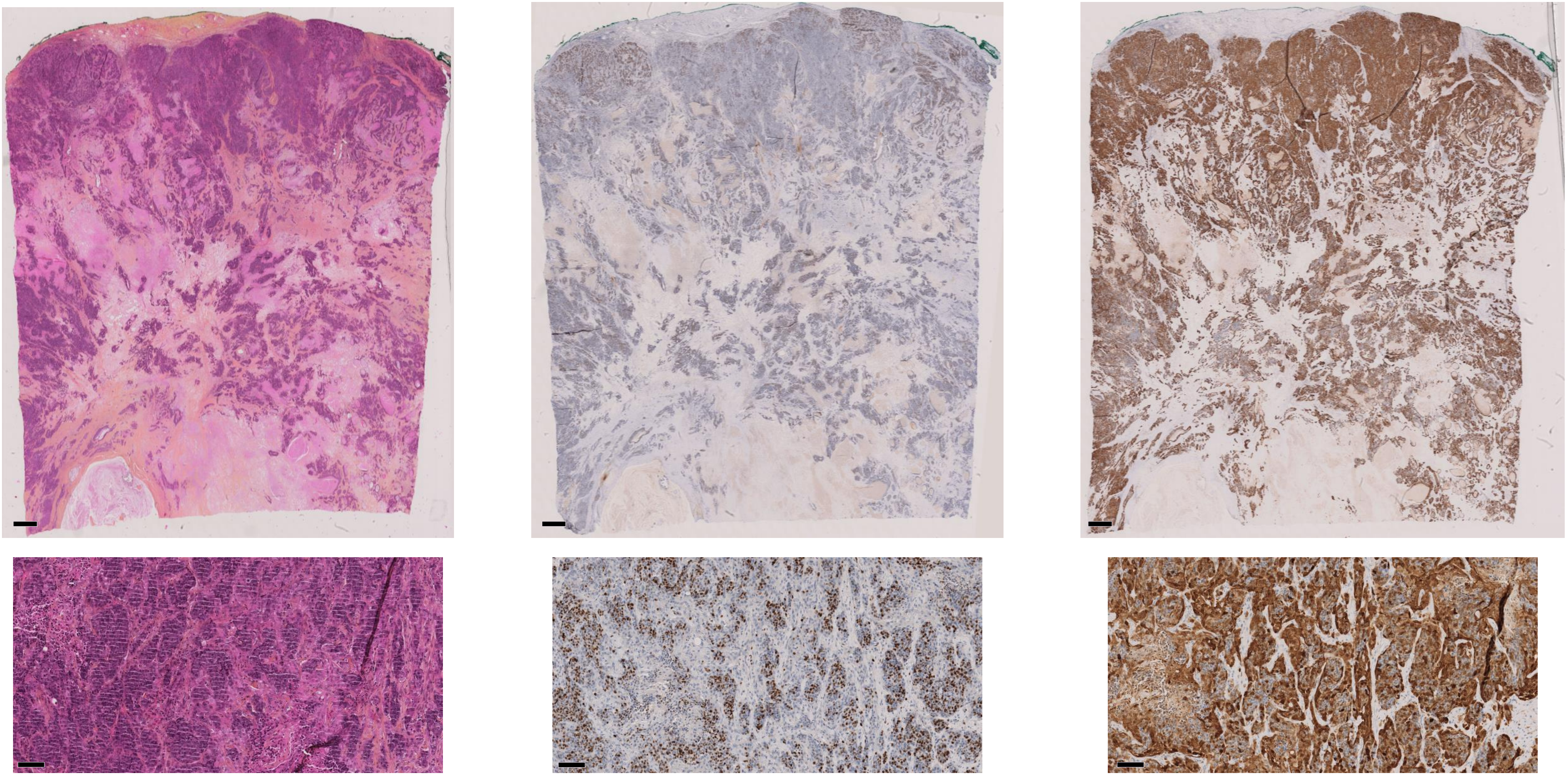

**HE** **KI67** **p16**

Scale bar = 1mm

Scale bar = 100µm

### ATC22 - JPI37 - Profile H

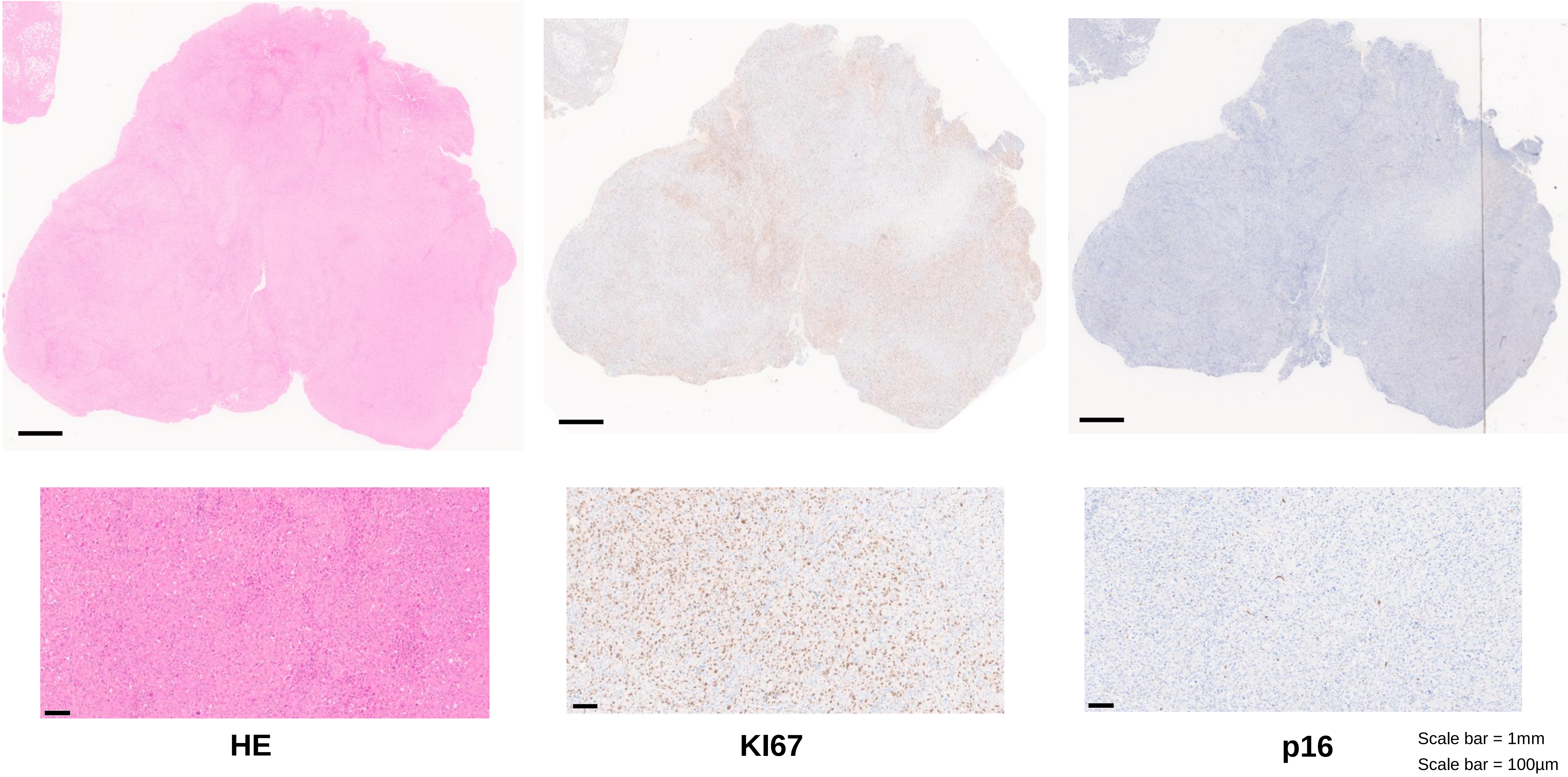

### ATC21 - JPI41 - Profile H

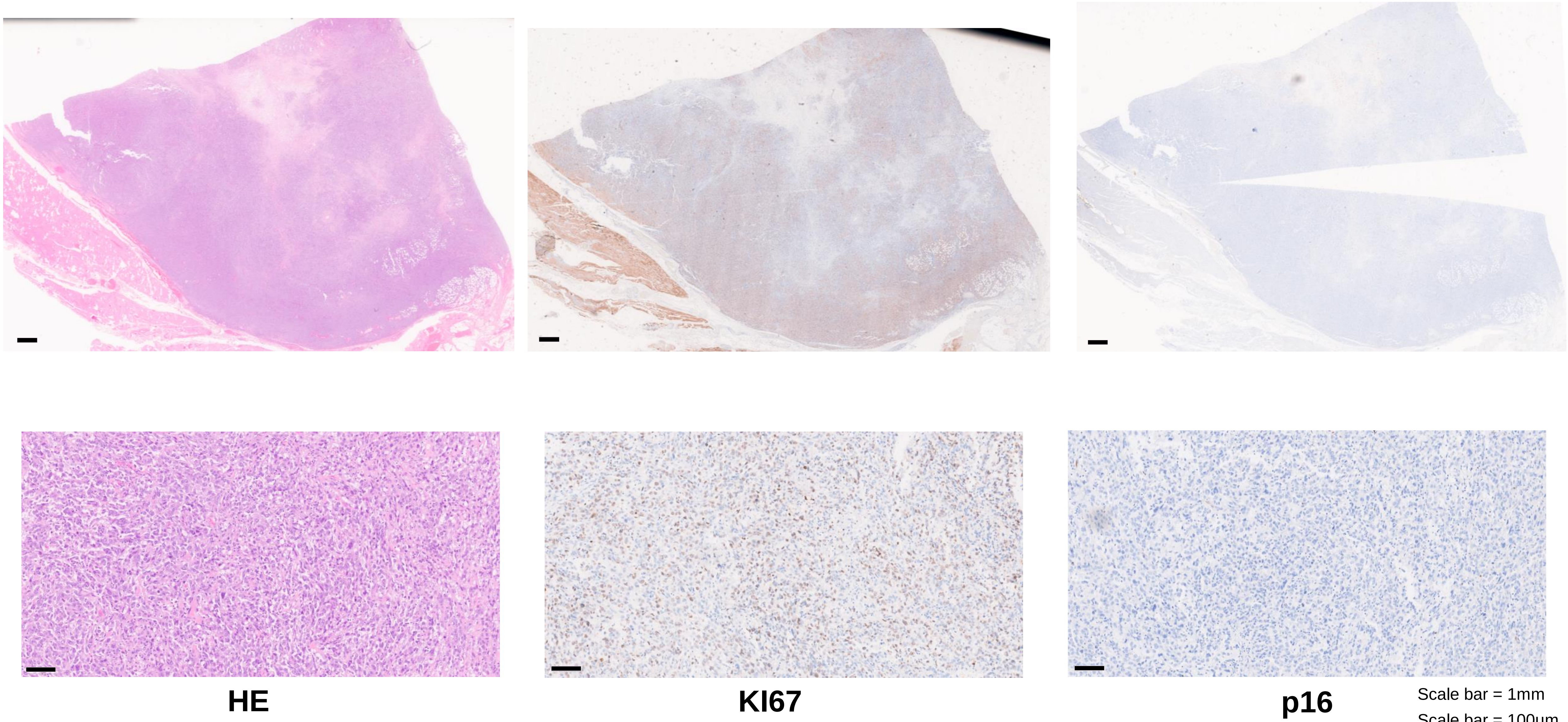


Scale bar = 1mm
Scale bar = 100µm

### ATC18 - JPI66 - Profile L

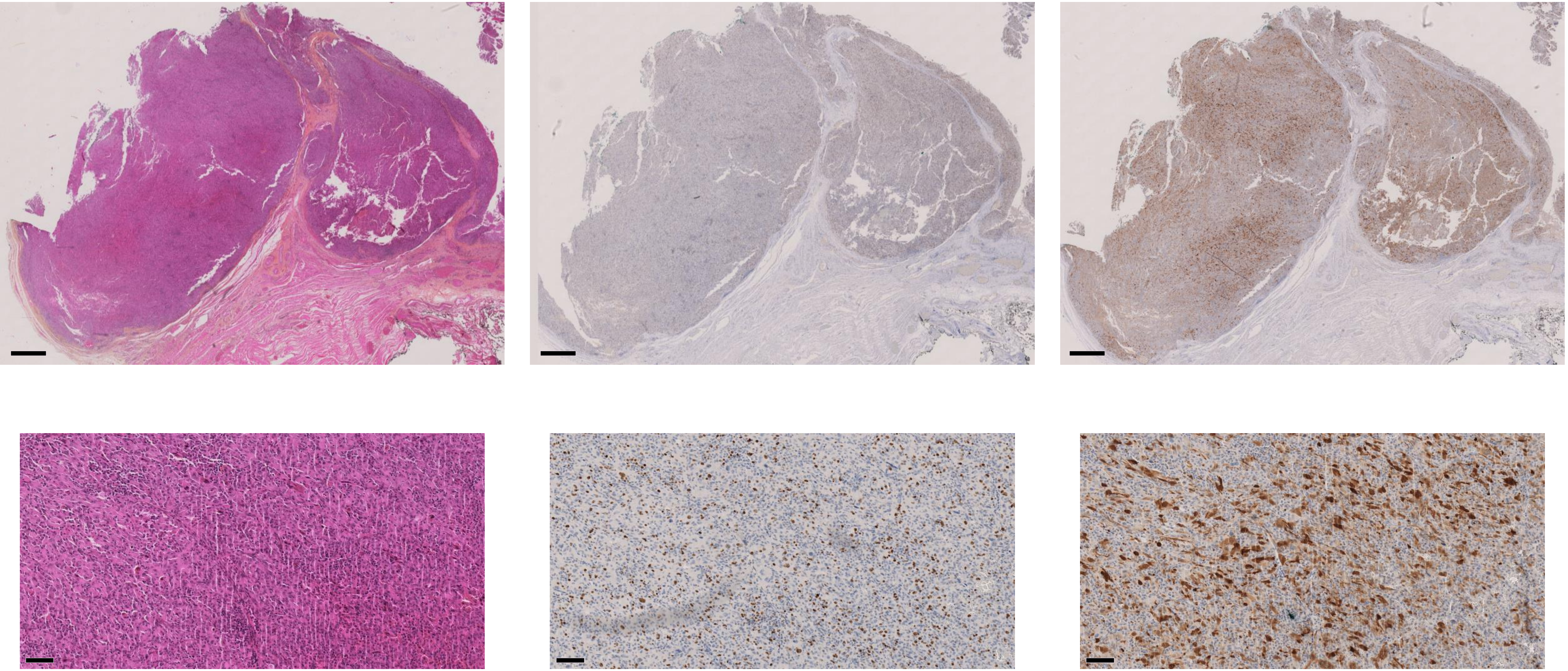

HE

KI67

p16

Scale bar = 1mm

Scale bar = 100µm

### ATC13 - JPI72 - Profile L

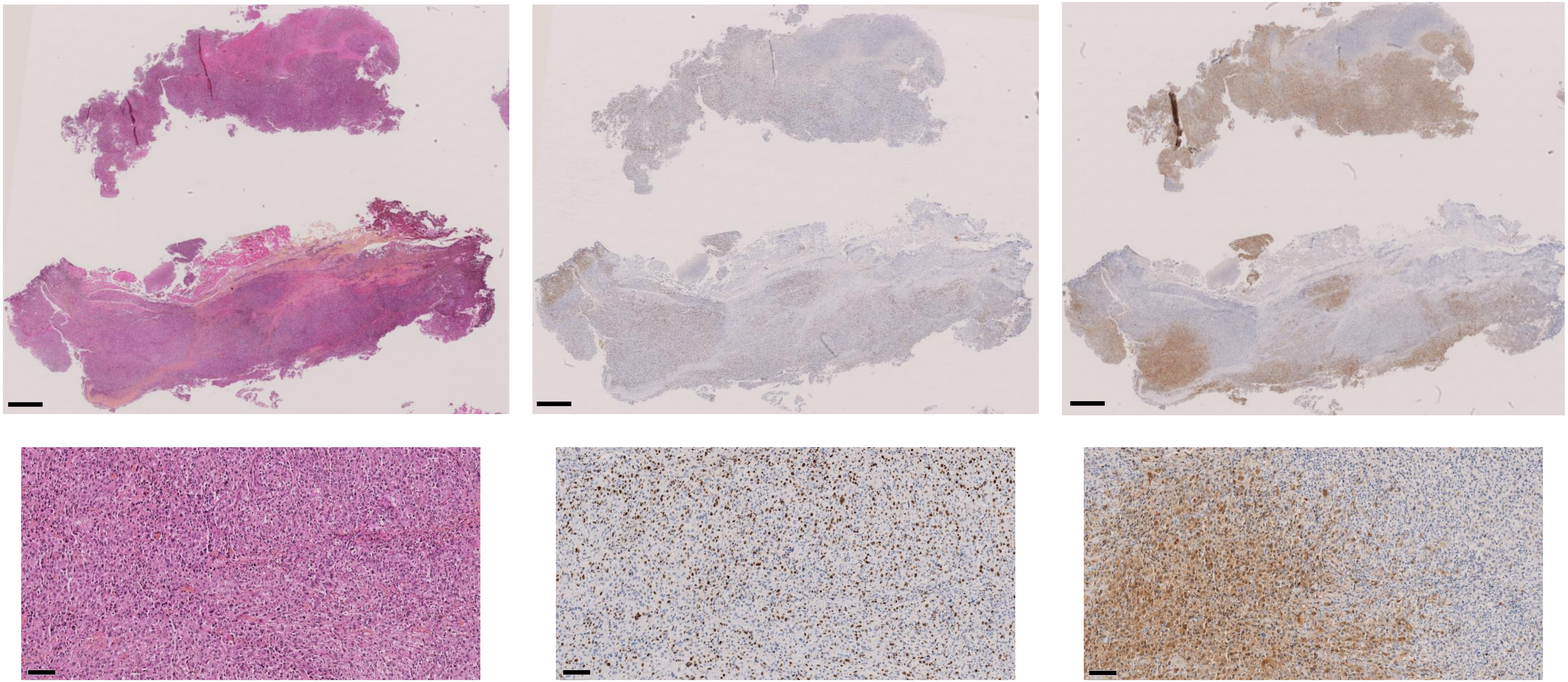

**HE** **KI67** **p16**

Scale bar = 1mm
Scale bar = 100µm

### ATC14 - JPI26 - Profile L

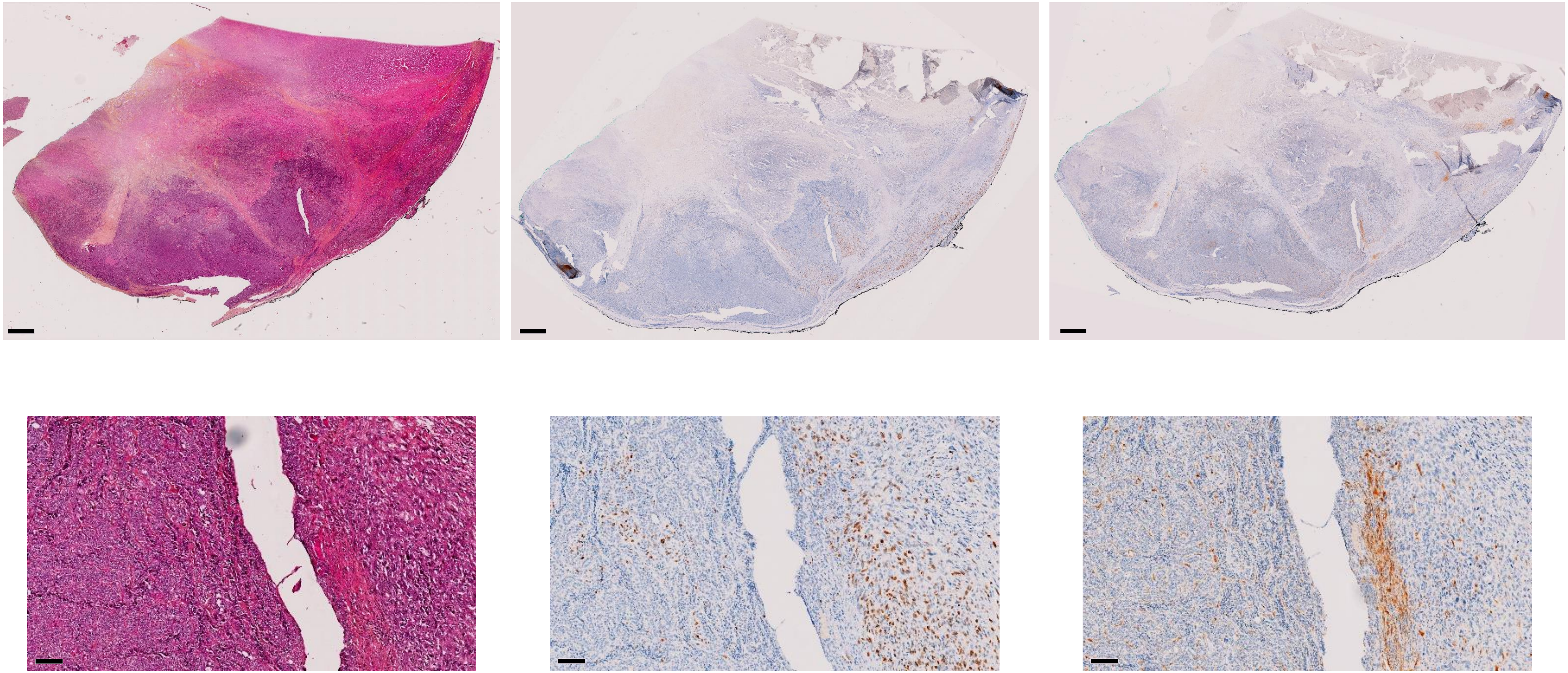

HE

KI67

p16

Scale bar = 1mm

Scale bar = 100µm

### ATC11 - JPI81 - Profile L

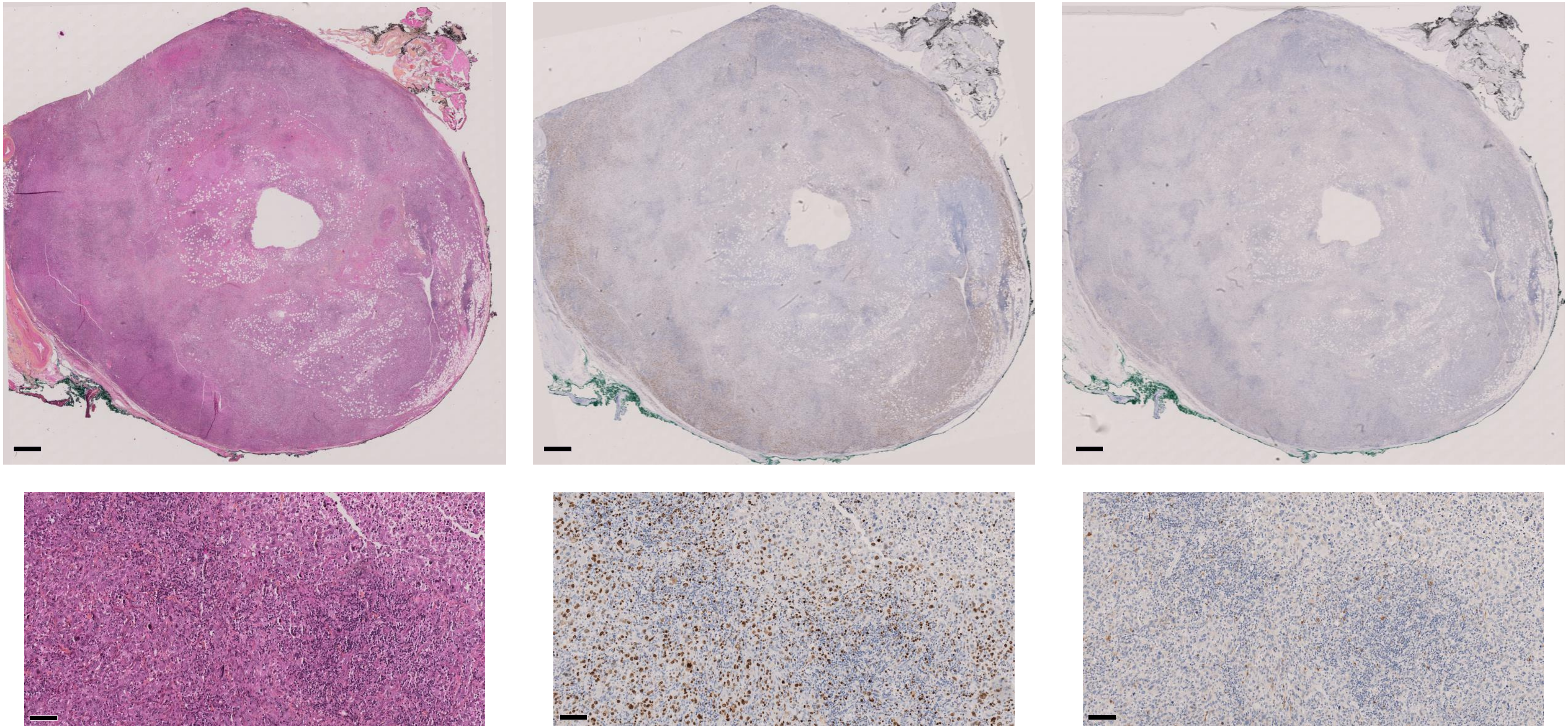

HE KI67 p16

Scale bar = 1mm
Scale bar = 100µm

### ATC19 - JPI76 - Profile L

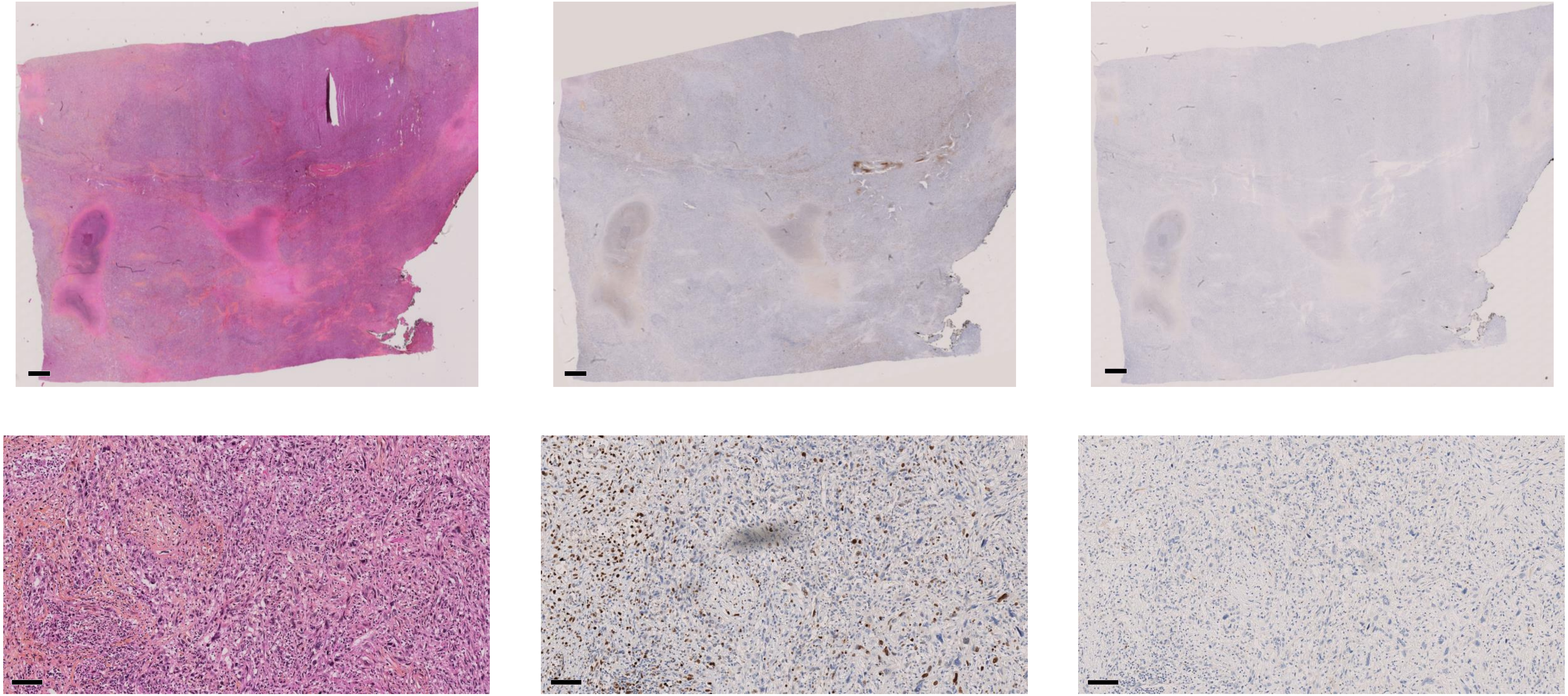

**HE** **KI67** **p16**

Scale bar = 1mm

Scale bar = 100µm

### PDTC1 - JPI25 - Profile A

#### Vascular/lymphatic invasion of the tumor (TTF-1$^{+}$ and PAX8$^{+}$) with lymphangitis carcinomatosa aspect

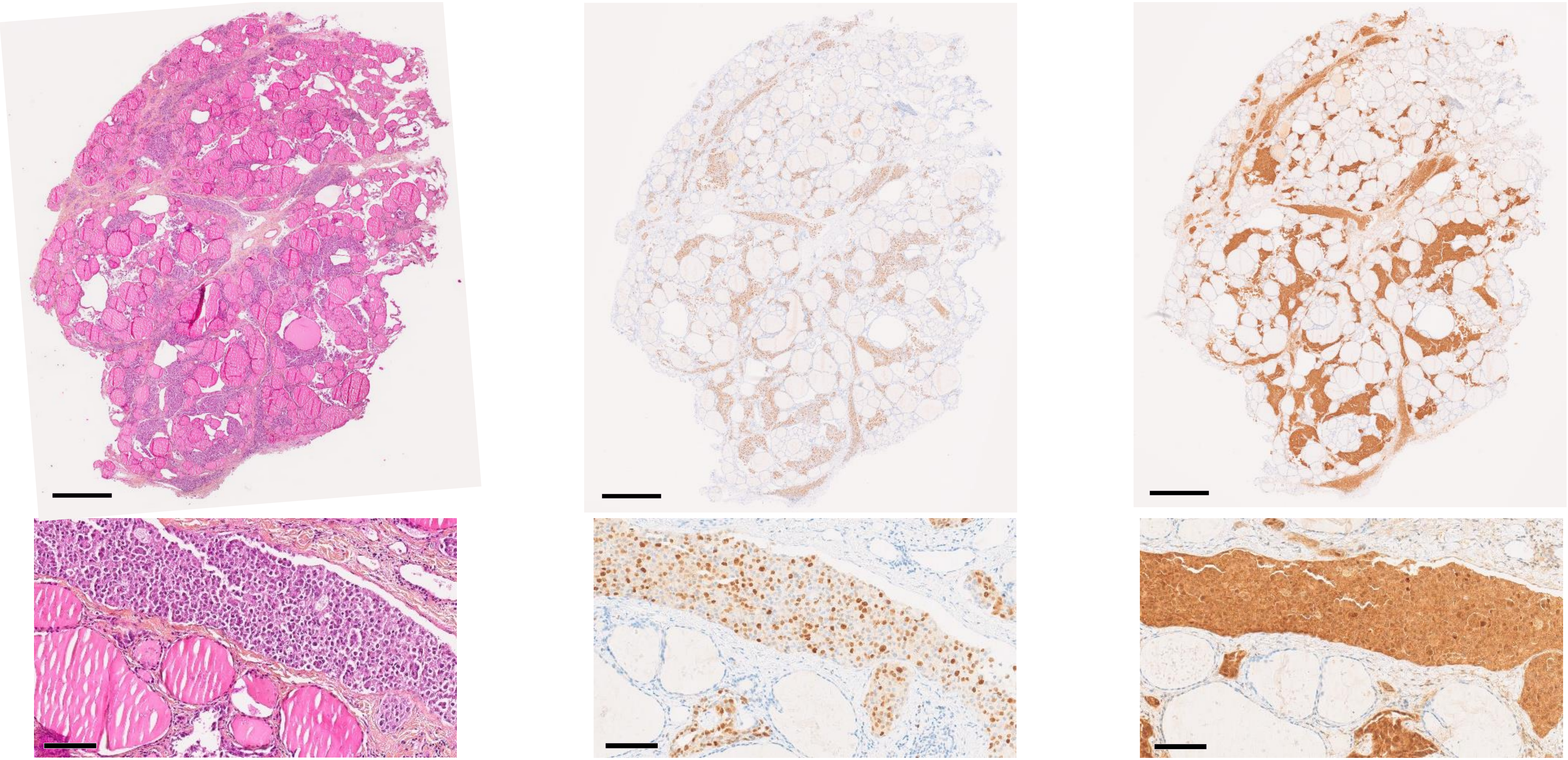

HE KI67 p16

Scale bar = 1mm

Scale bar = 100µm

### PDTC13 - JPI67 - Profile H

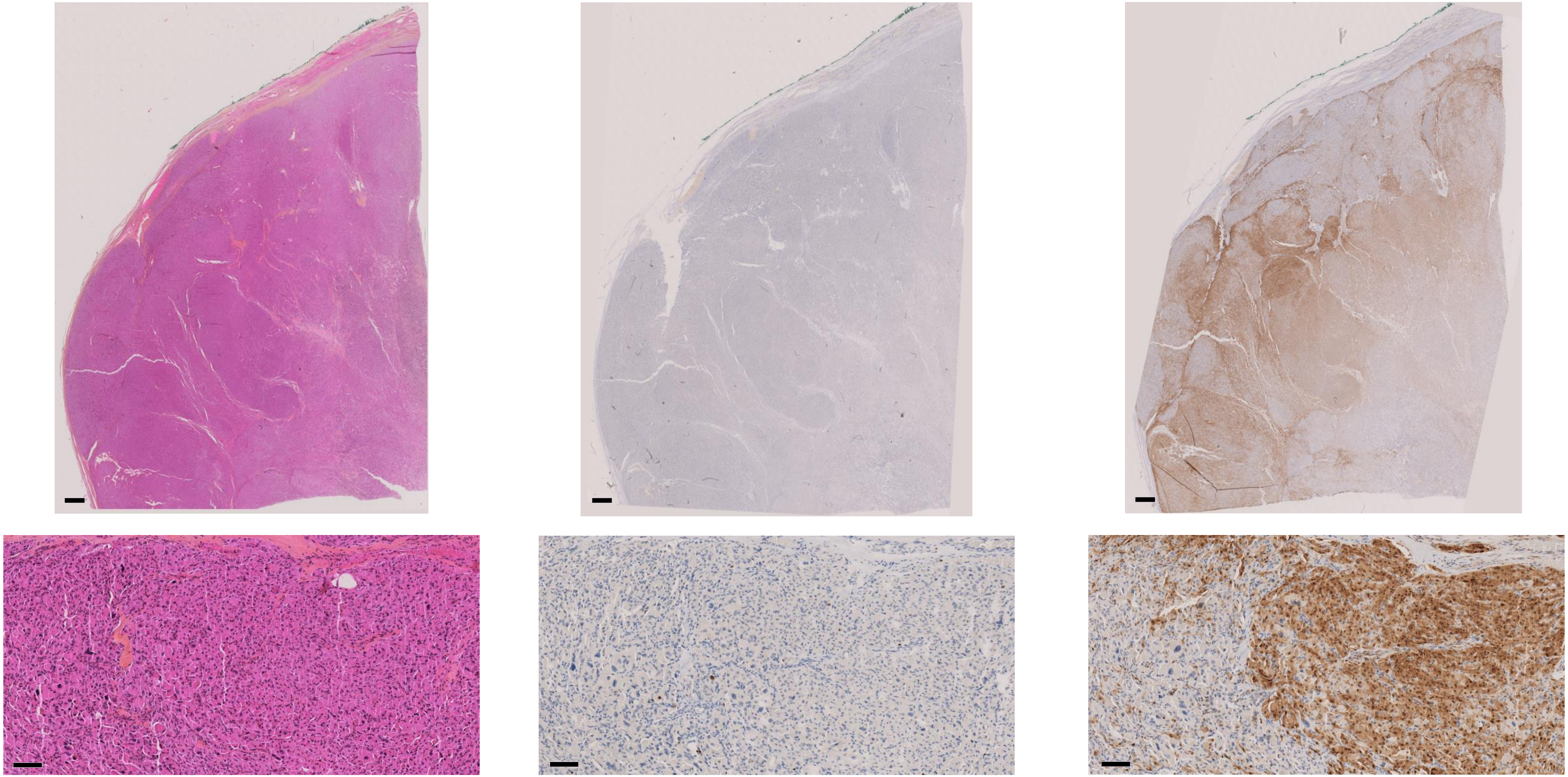

HE KI67 p16

Scale bar = 1mm
Scale bar = 100µm

### PDTC20 - JPI29 - Profile H

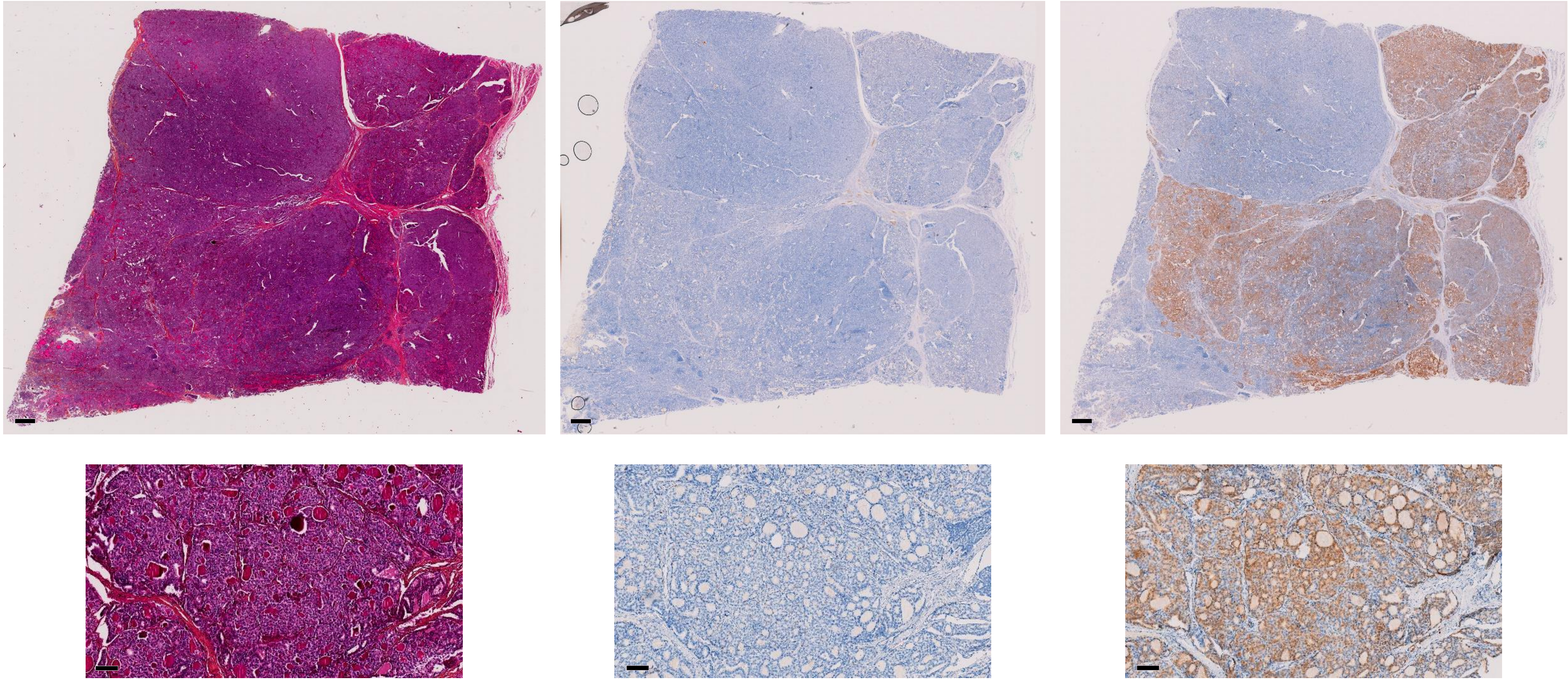

HE KI67 p16

Scale bar = 1mm
Scale bar = 100µm

### PDTC14 - JPI75 - Profile H

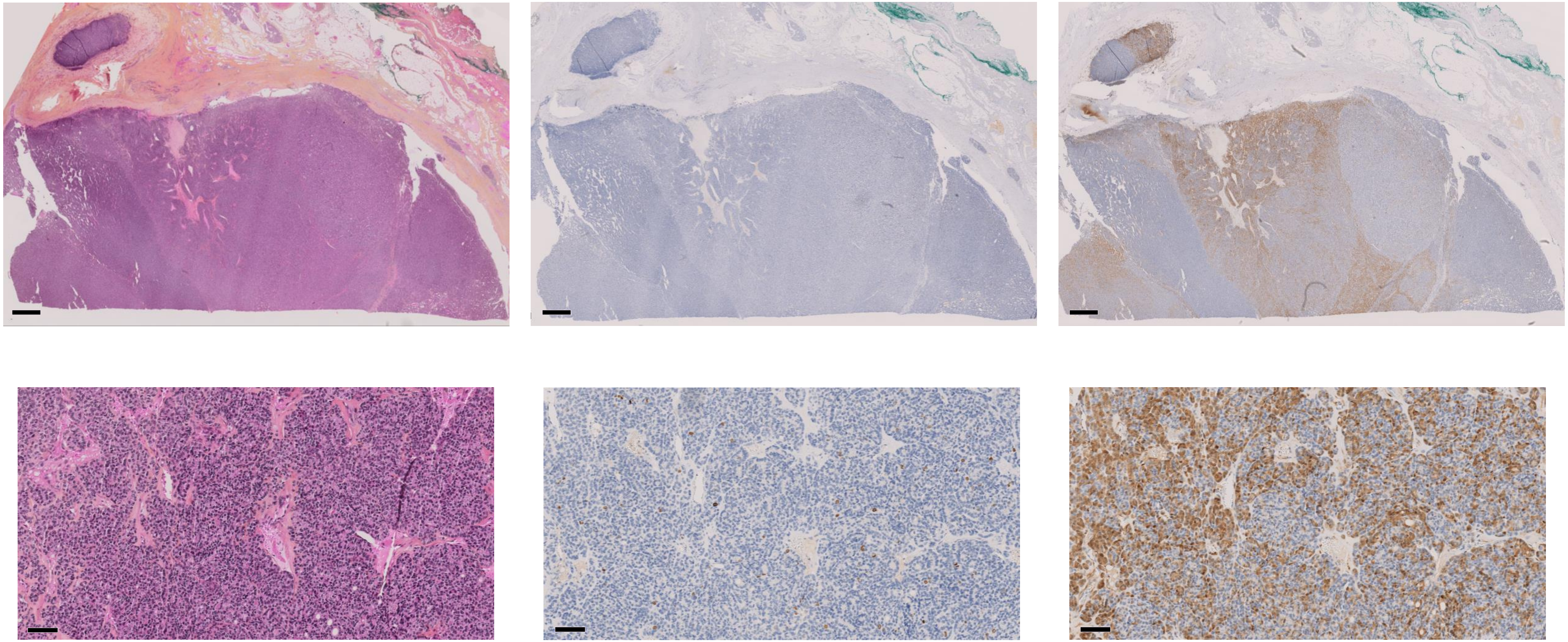

**HE** **KI67** **p16**

Scale bar = 1mm

Scale bar = 100µm

### PDTC18 - JPI34 - Profile H

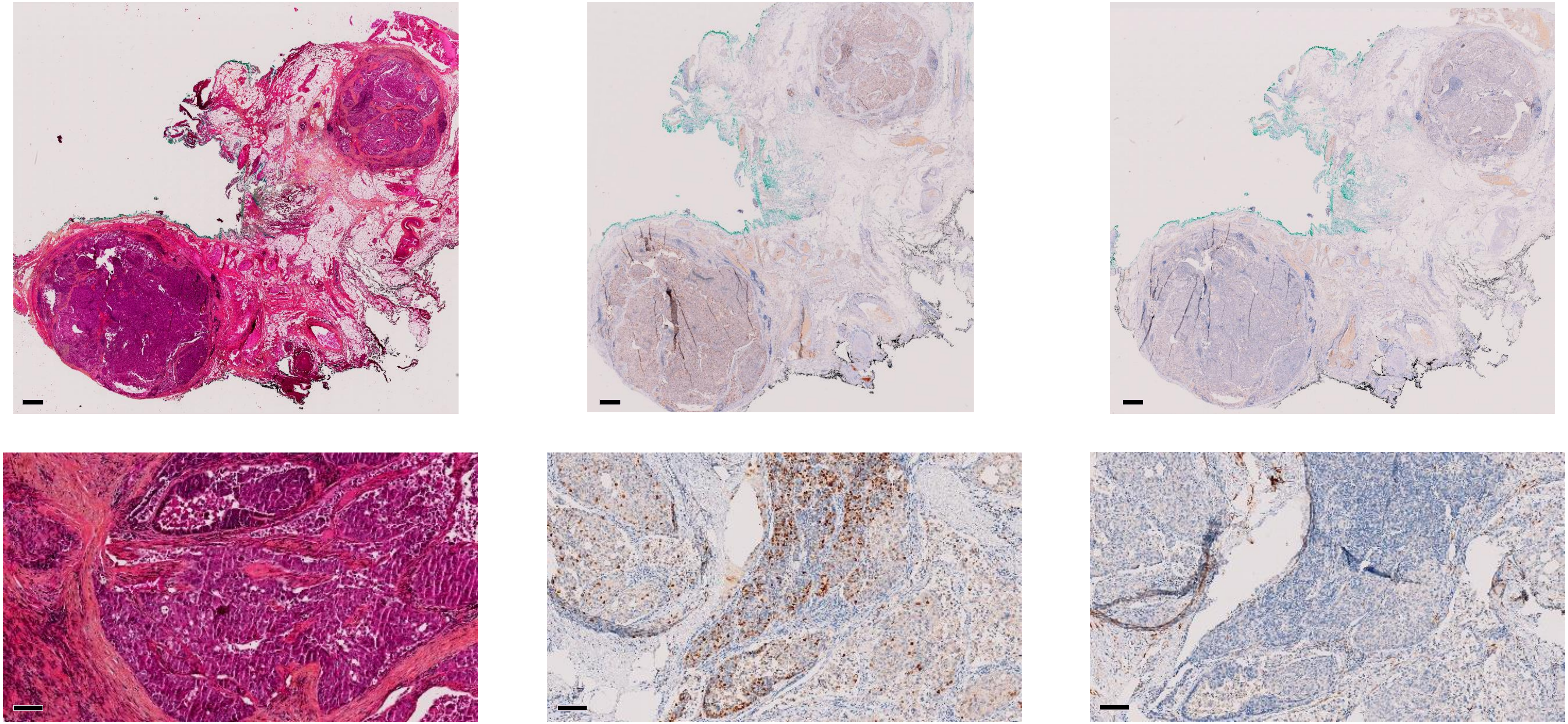

HE KI67 p16

Scale bar = 1mm
Scale bar = 100µm

### PDTC12 - JPI19 - Profile H

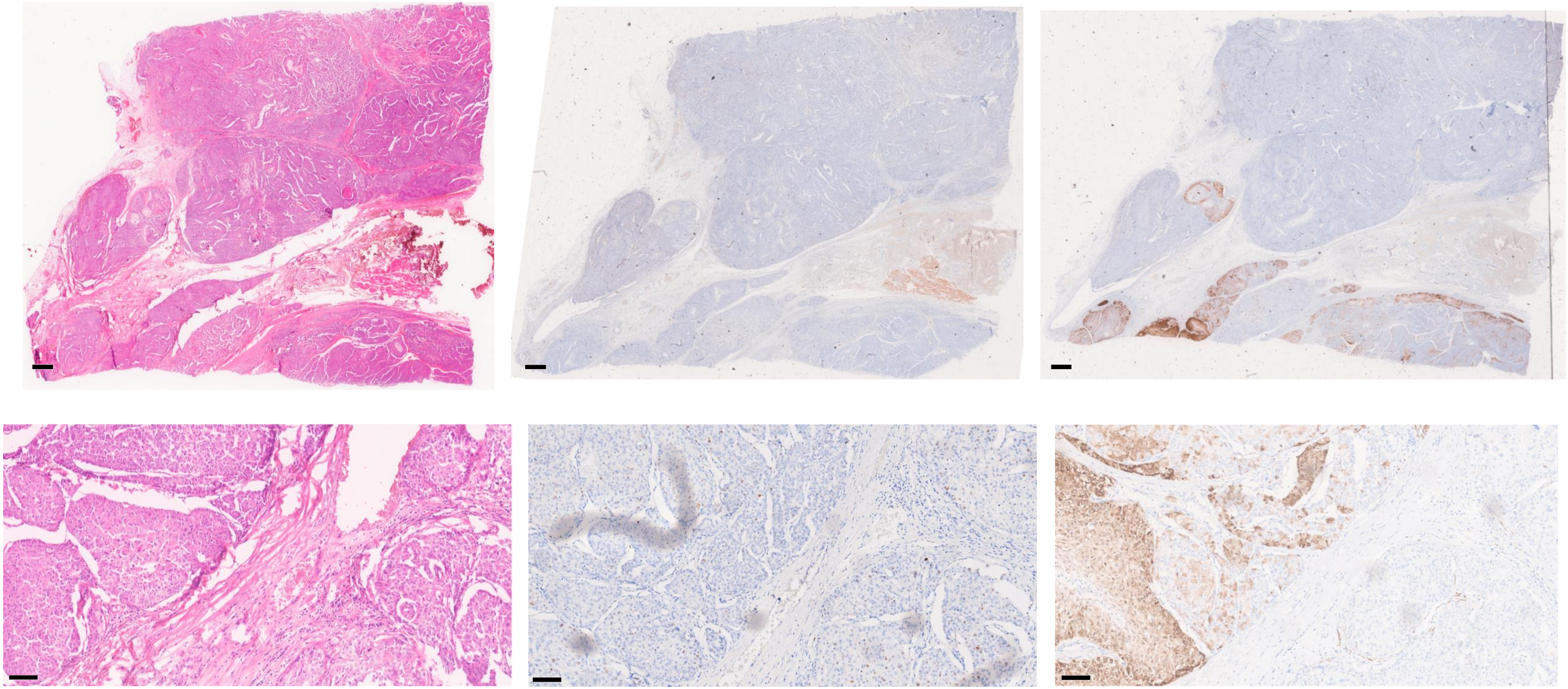

**HE** **KI67** **p16**

Scale bar = 1mm

Scale bar = 100µm

### PDTC16 - JPI73 - Profile H

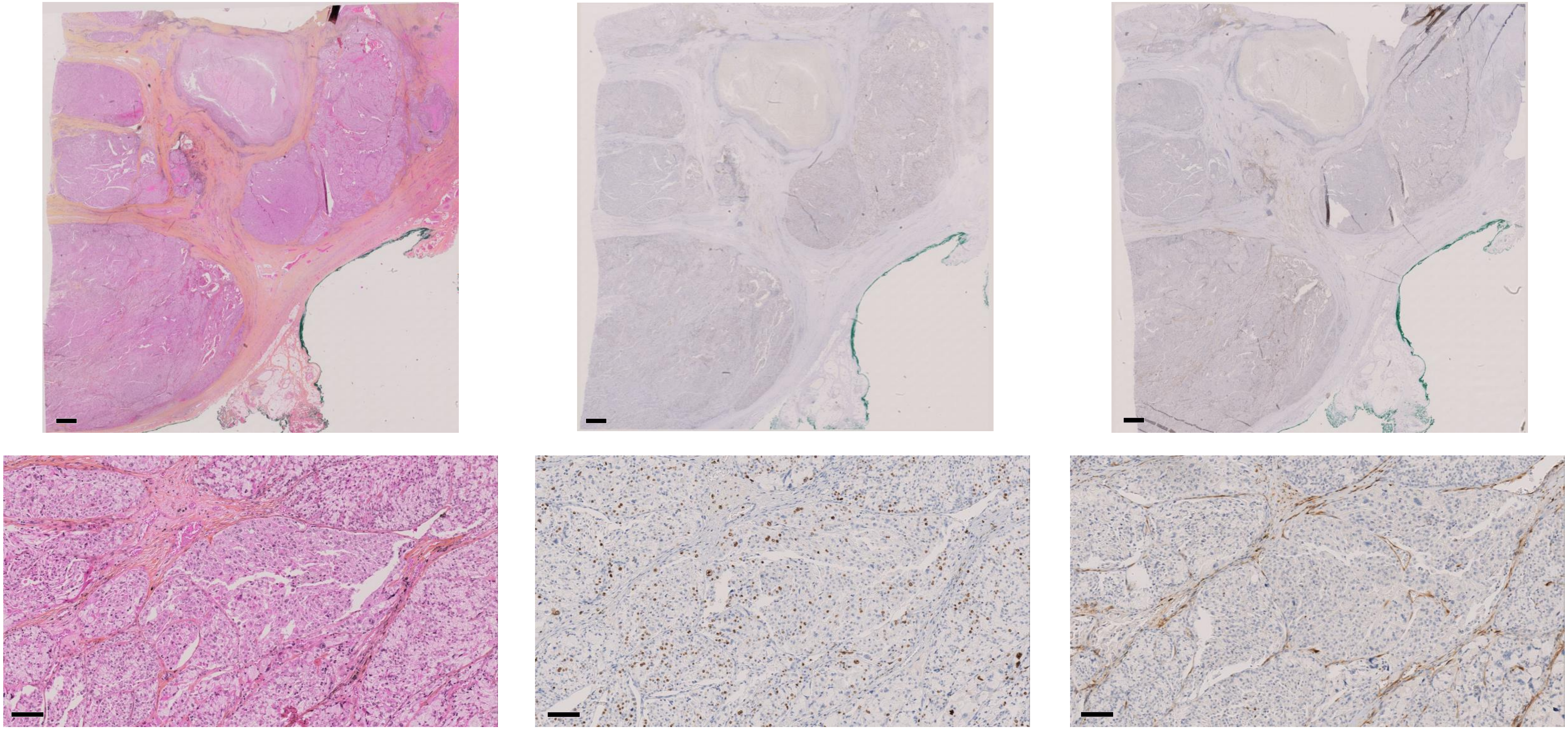

**HE** **KI67** **p16**

Scale bar = 1mm

Scale bar = 100µm

### PDTC19 - JPI40 - Profile H

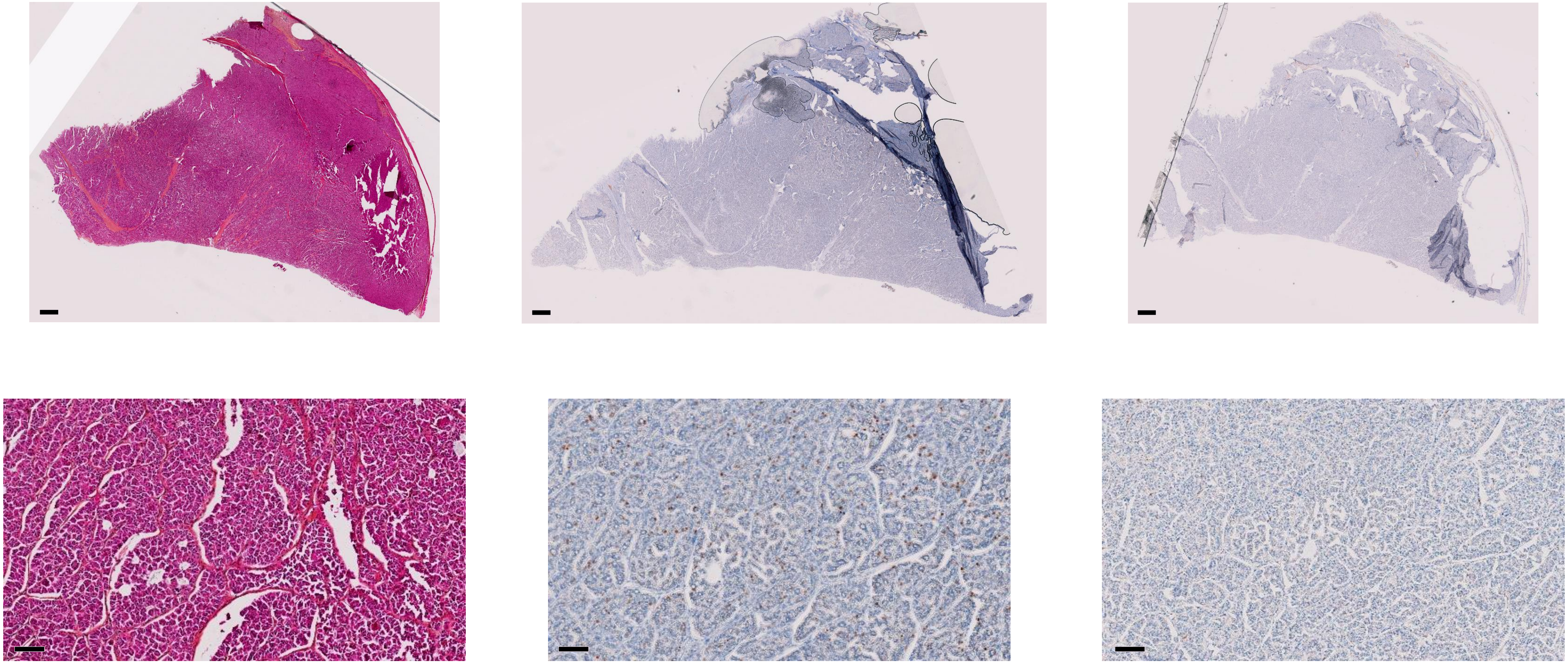

HE KI67 p16

Scale bar = 1mm

Scale bar = 100µm

### PDTC15 - JPI68 - Profile H

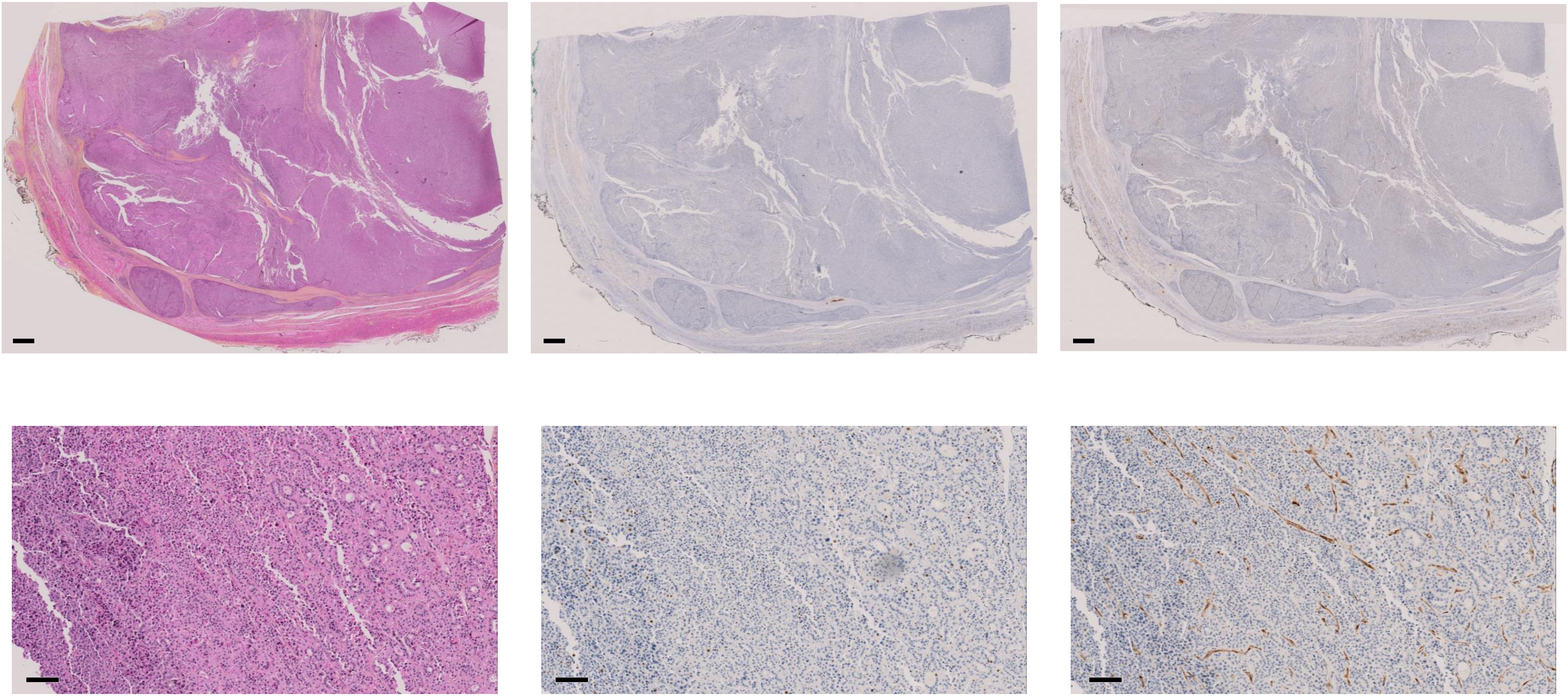

**HE** **KI67** **p16**

Scale bar = 1mm

Scale bar = 100µm

### PDTC3 - JPI83 - Profile L

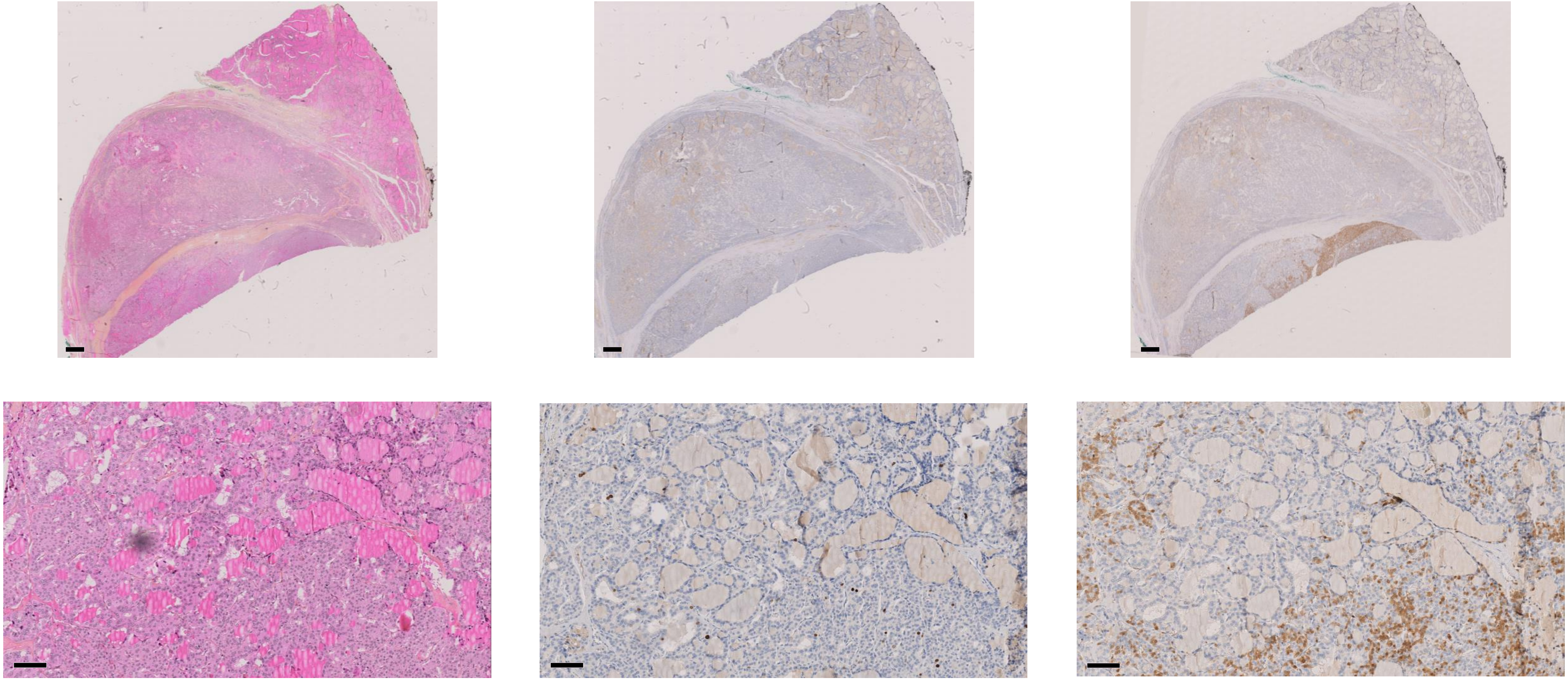

HE KI67 p16

Scale bar = 1mm

Scale bar = 100µm

### PDTC8 - JPI33 - Profile L

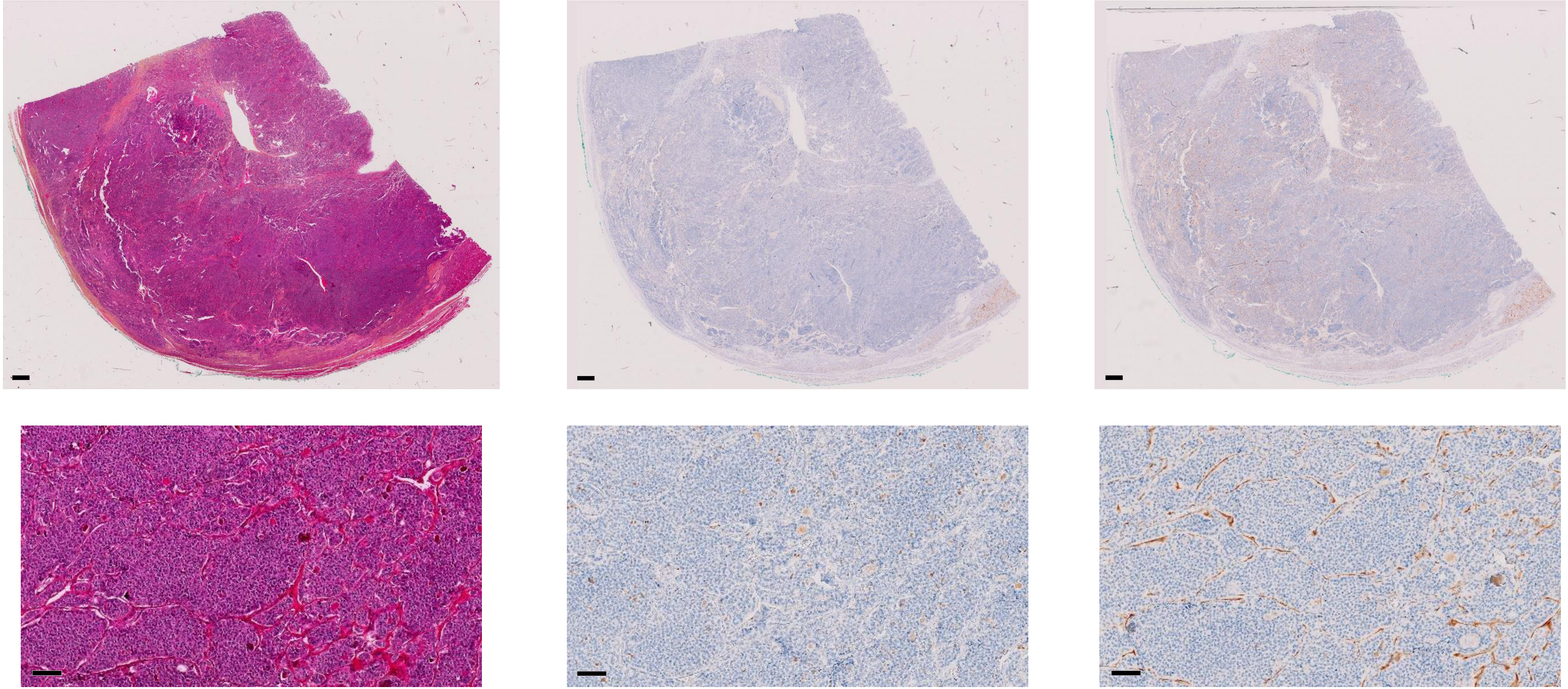

**HE** **KI67** **p16**

Scale bar = 1mm

Scale bar = 100µm

### PDTC7 - JPI31 - Profile L

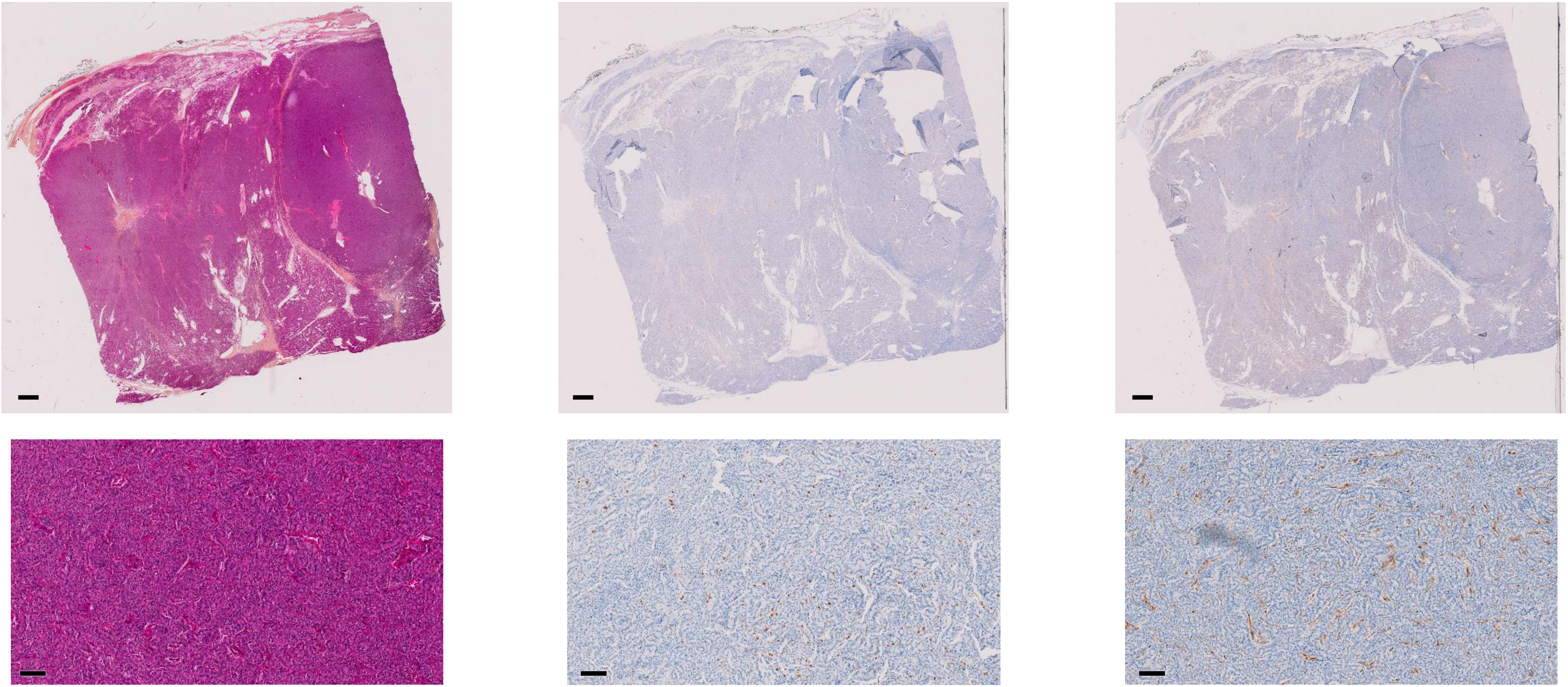

**HE** **KI67** **p16**

Scale bar = 1mm
Scale bar = 100µm

### PDTC6 - JPI82 - Profile L

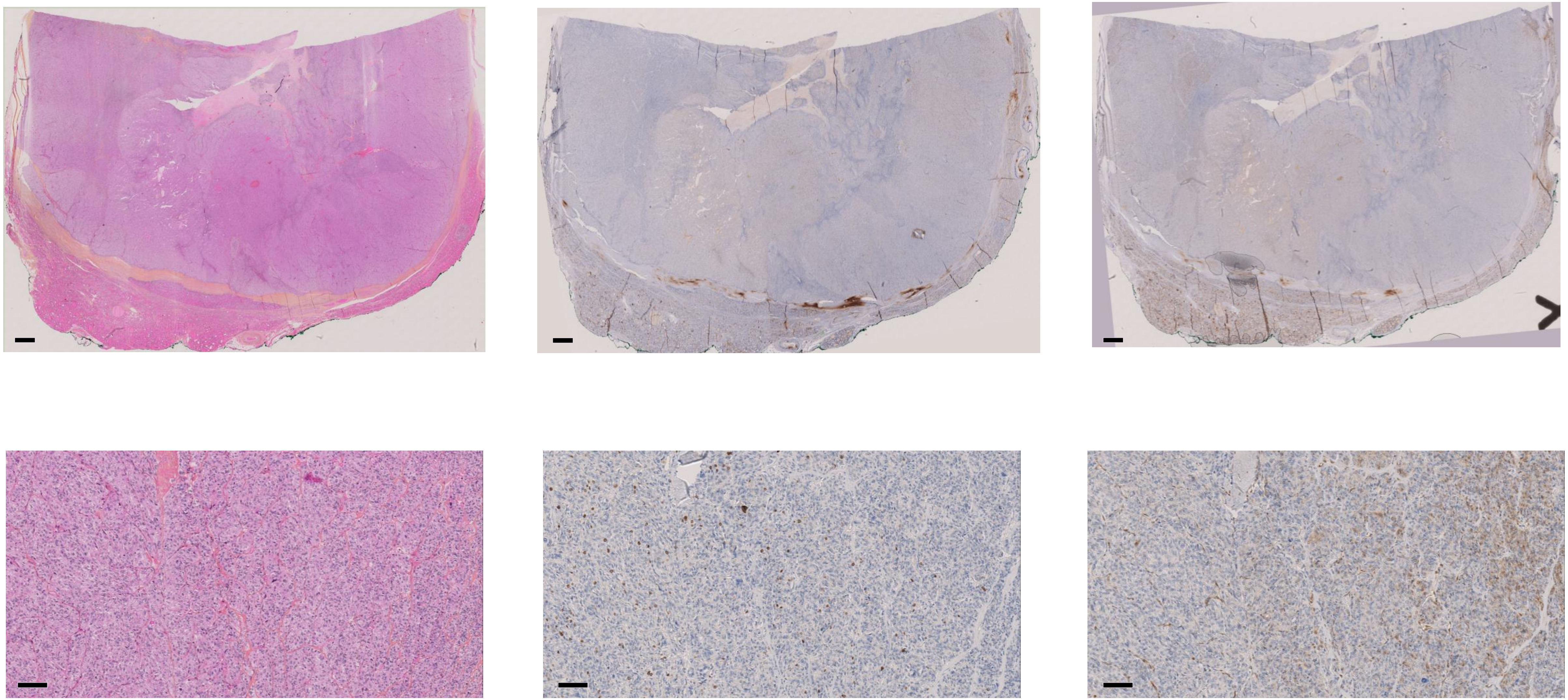

**HE** **KI67** **p16**

Scale bar = 1mm

Scale bar = 100µm

### PDTC11 - JPI79 - Profile L

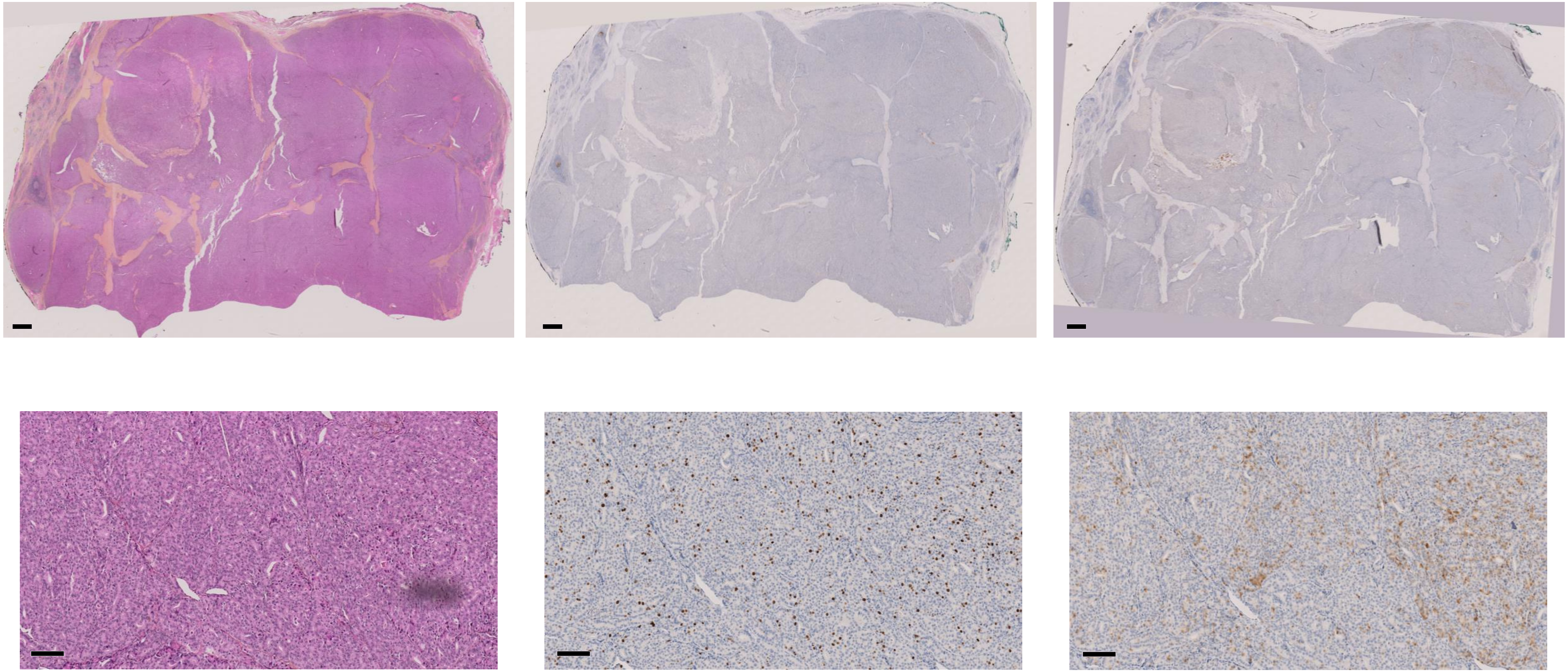

HE

KI67

p16

Scale bar = 1mm

Scale bar = 100µm

### PDTC2 - JPI70 - Profile L

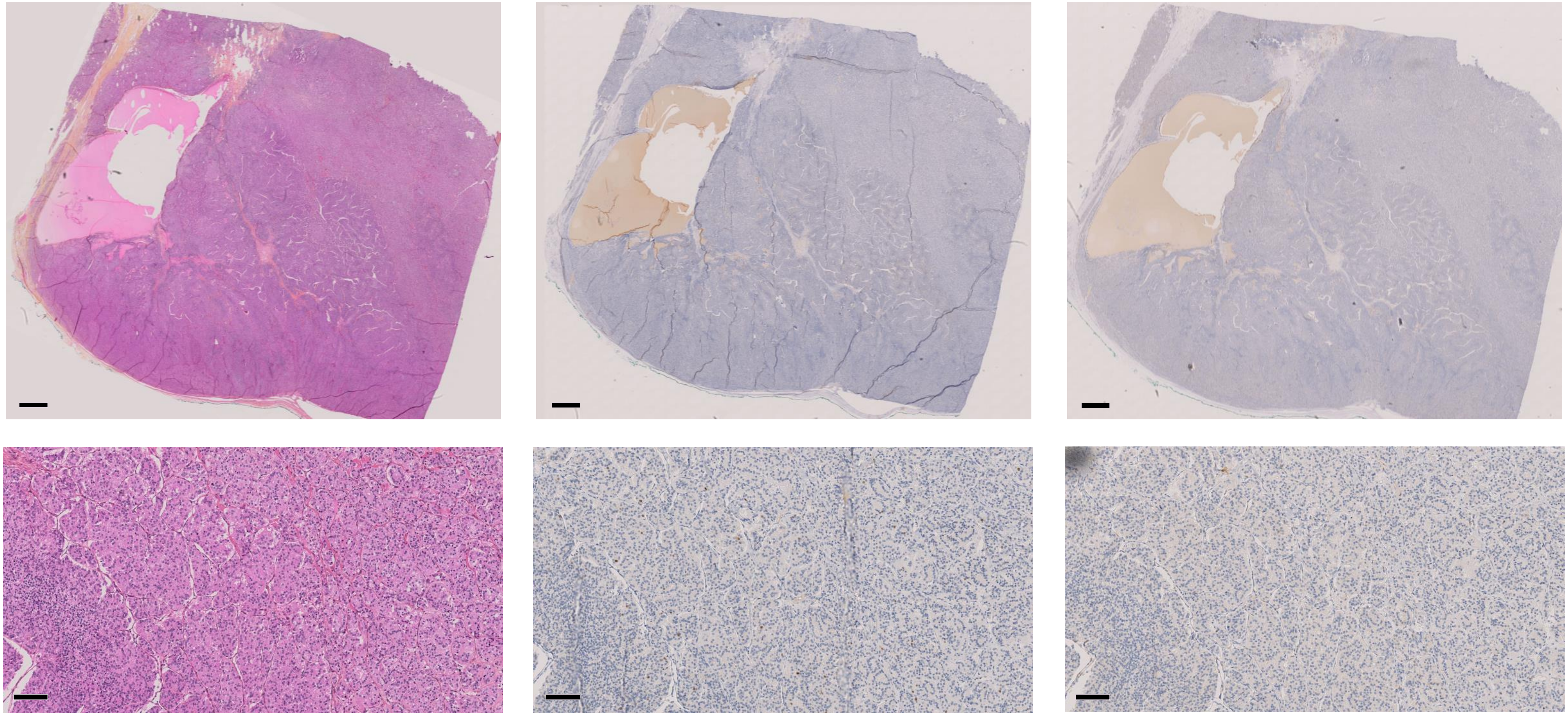

HE KI67 p16

Scale bar = 1mm
Scale bar = 100µm

### PDTC21 - JPI22 - spot 1 only

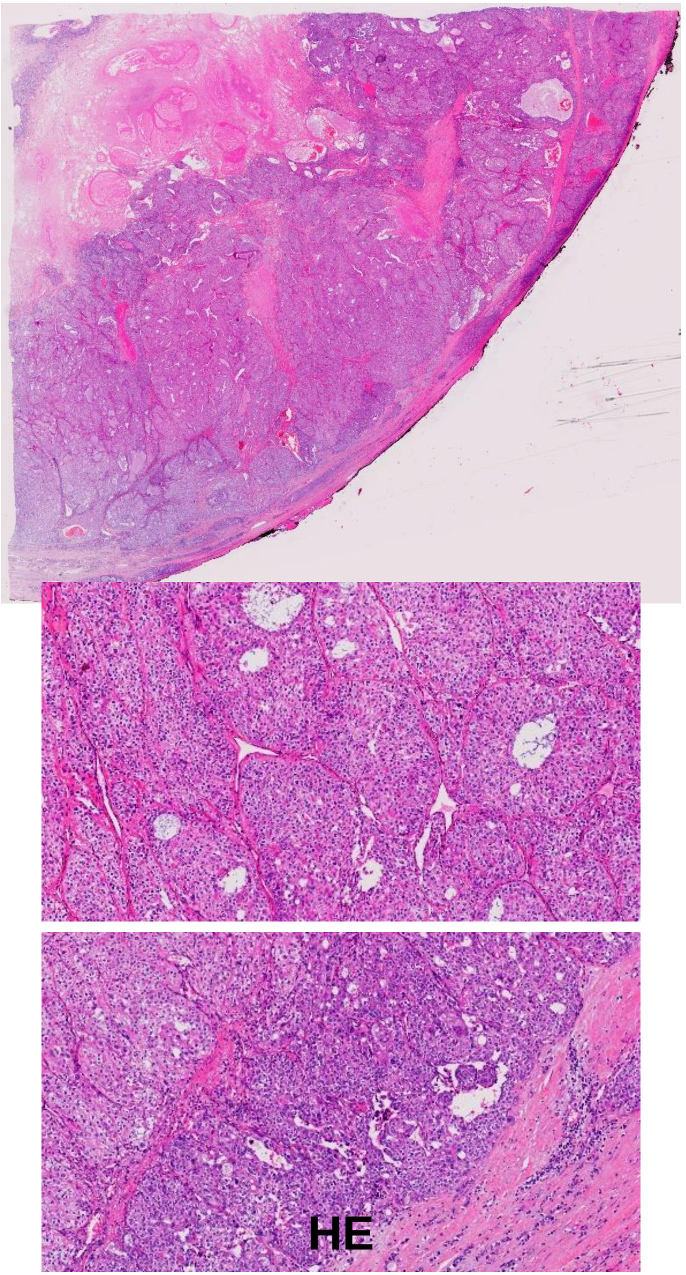


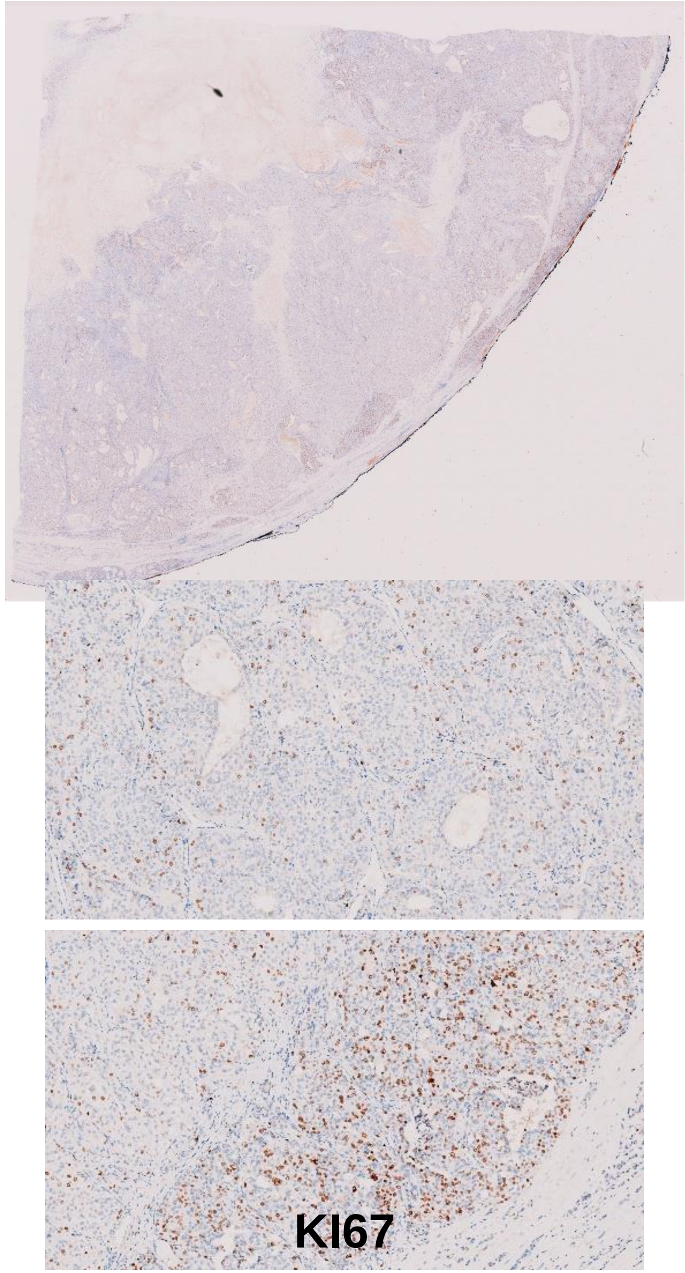


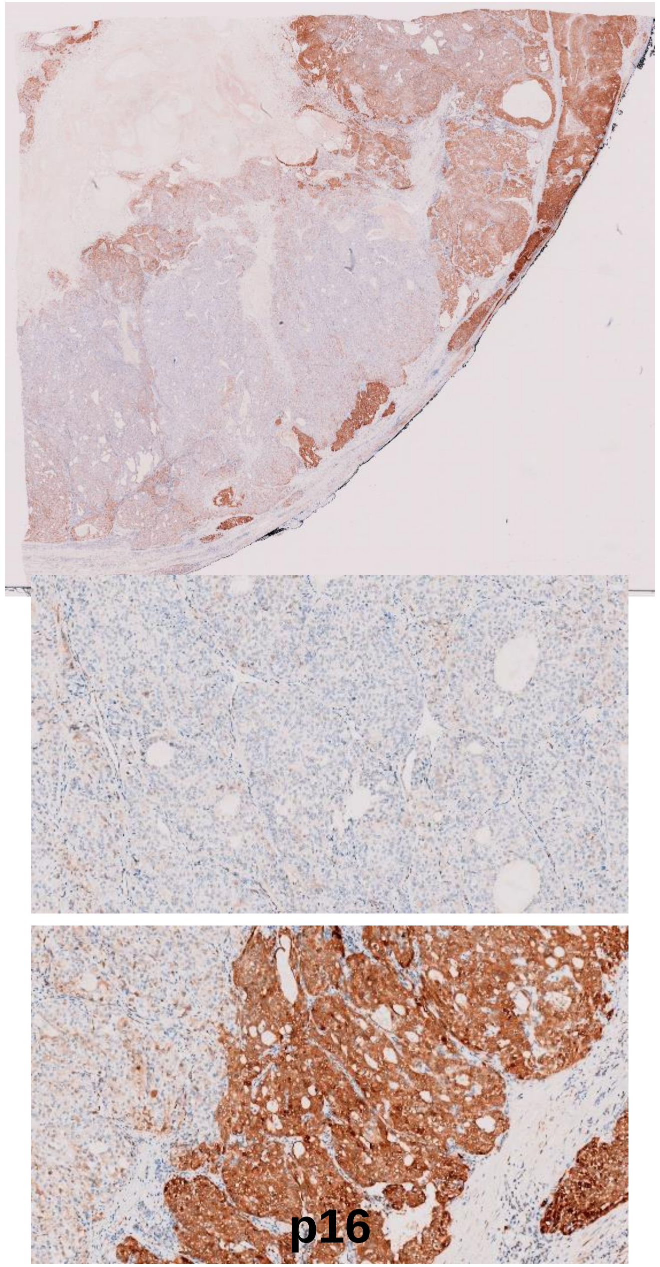

### PDTC22 - JPI17 - Profile H

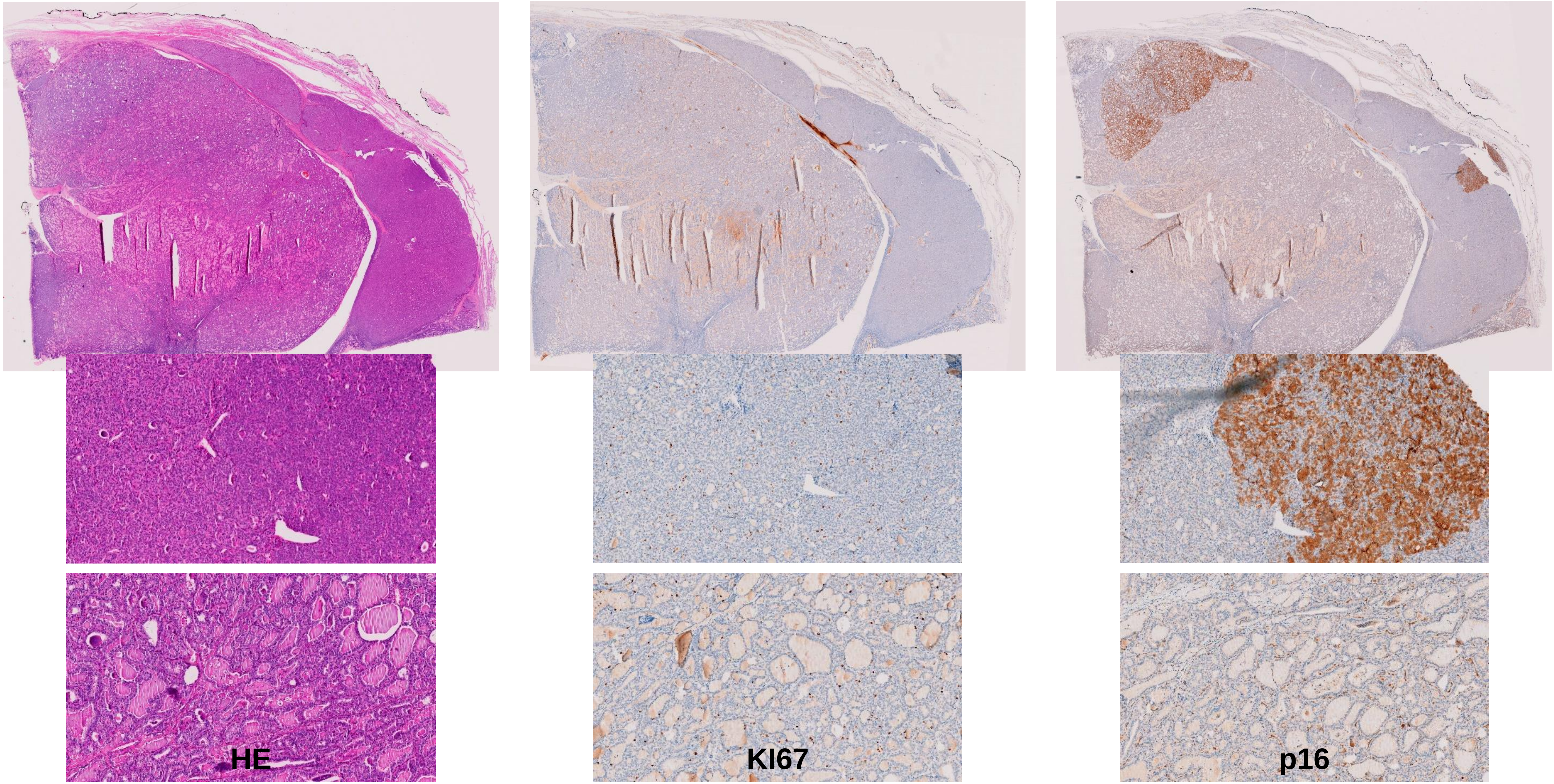
