## Supplementary Figures 1-3, 5-11 for "CDK4 phosphorylation status and rational use for combining CDK4/6 and BRAF/MEK inhibition in advanced thyroid carcinomas"

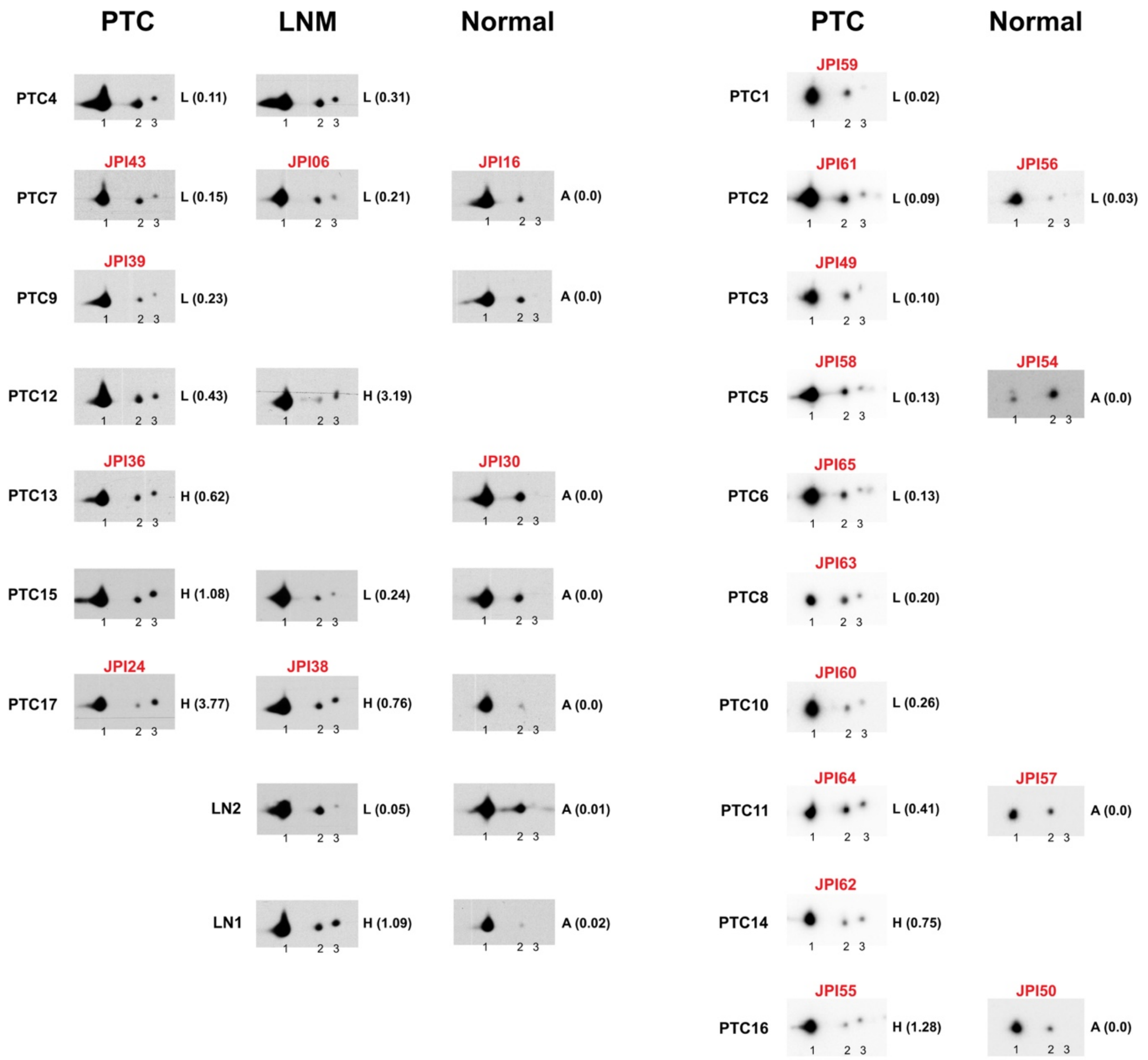


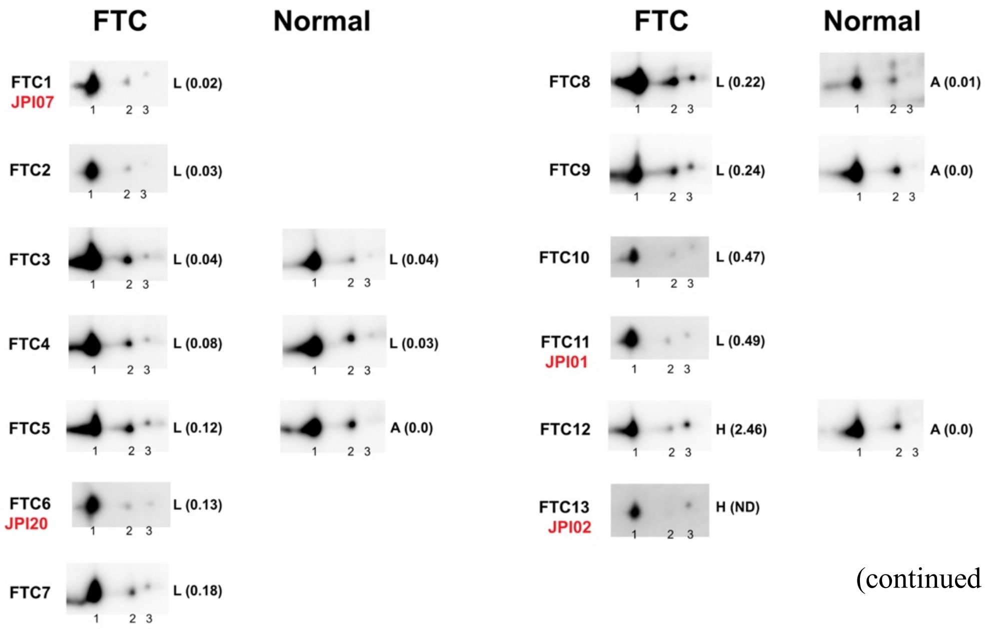


(continued on next page)

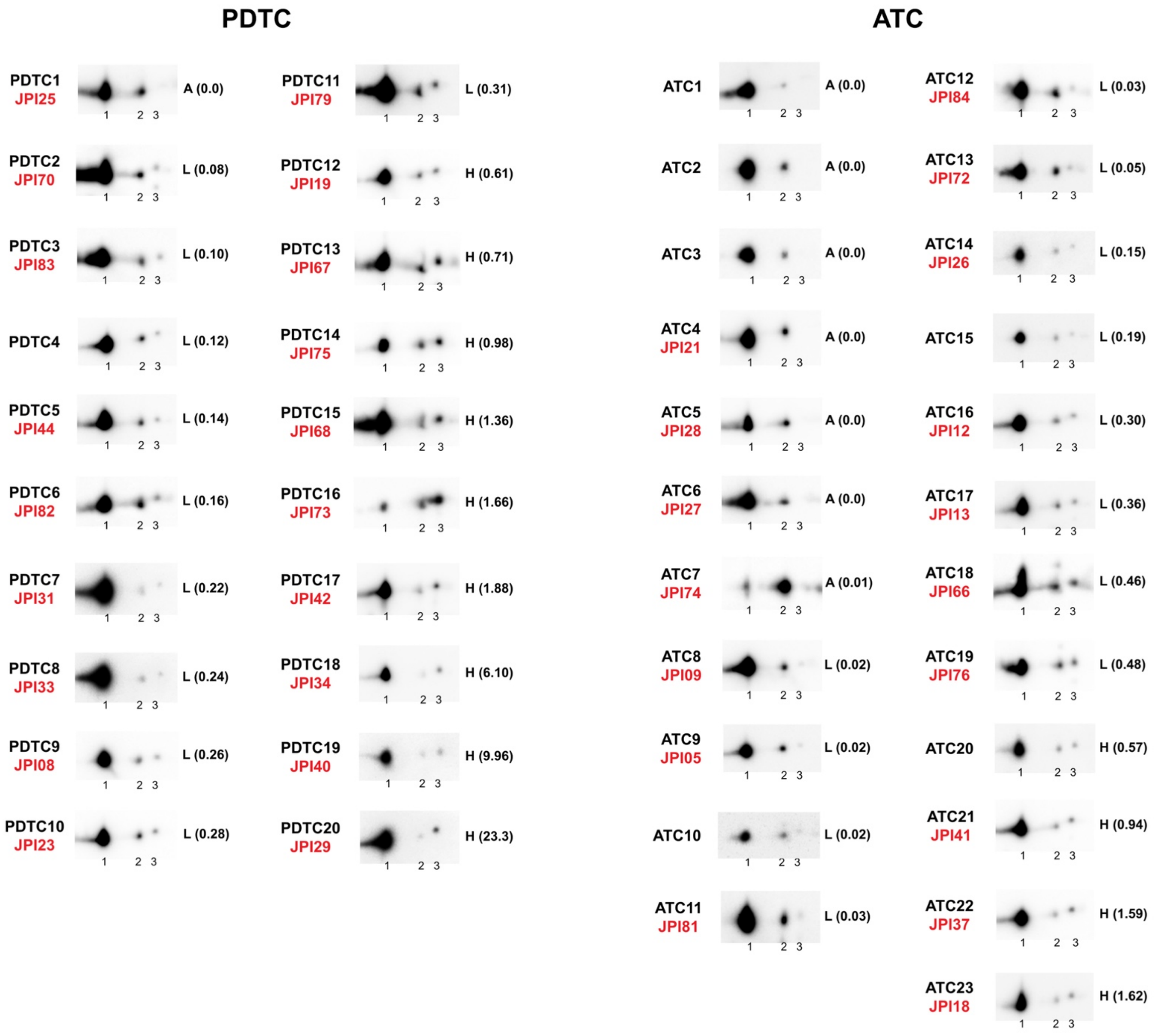


**Supplementary Figure S1. Immunodetections of CDK4 after 2D-gel electrophoresis separation of whole protein extracts from the fresh-frozen samples cohort.** Corresponding LNM and/or normal thyroid tissue from some PTC and FTC cases are presented at the right of the tumor sample. Oncocytic FTC tumors are included in FTC panel. RNA-seq IDs are shown in red. CDK4 modification profiles were defined based on the ratio of T172-phosphorylated form of CDK4 (spot 3) over another modified form (spot 2), quantified from the immunoblots. H for high CDK4 phosphorylation: ratio $\geq 0.5$; L for low CDK4 phosphorylation: $0.02 \leq$ ratio $< 0.5$; A for absent CDK4 phosphorylation: ratio $< 0.02$. Unmodified native form of CDK4 is labelled as spot 1.

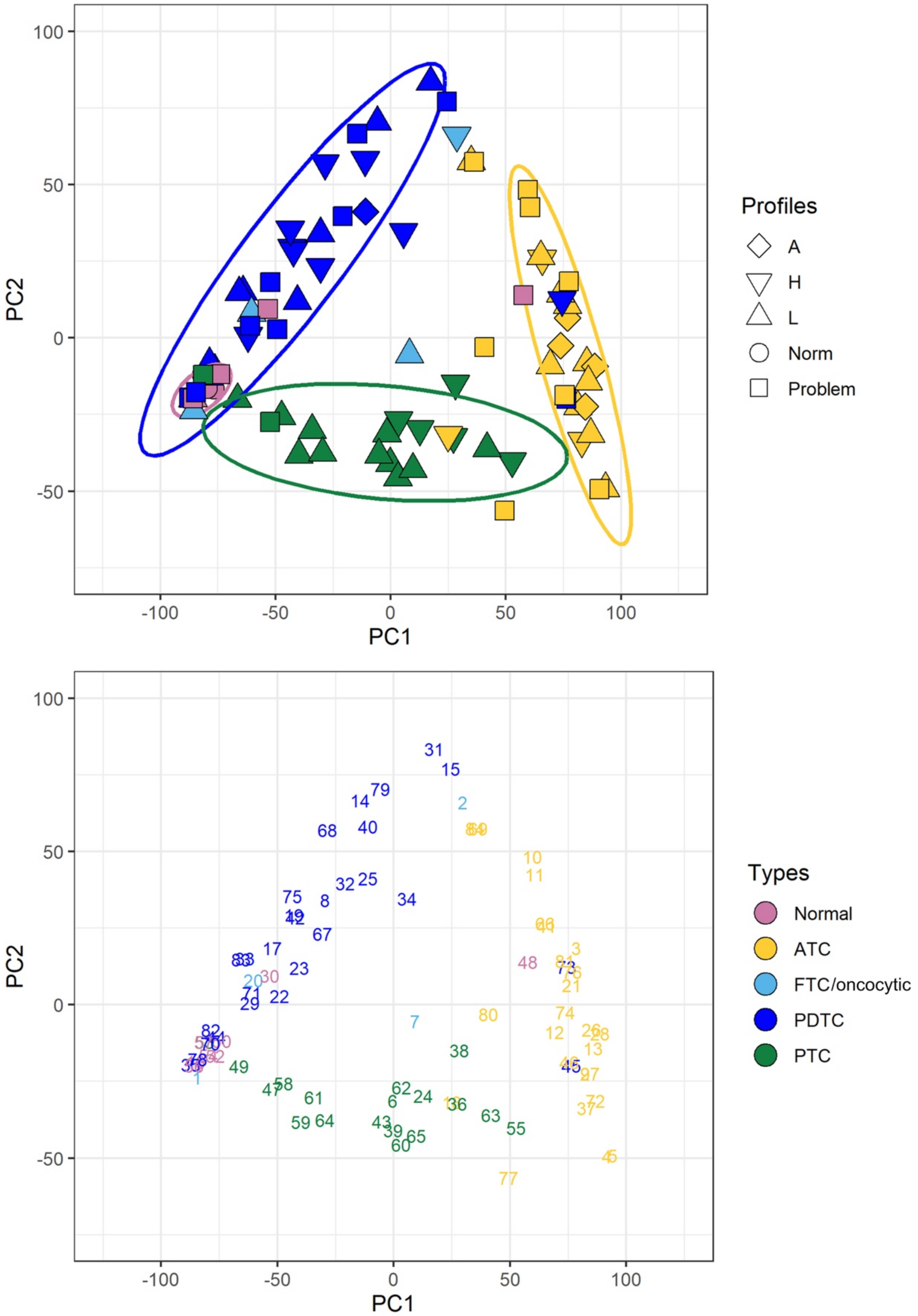


**Supplementary Figure S2. Representation of the gene expression variance between all 84 RNA-seq analyzed samples by principal component analysis.** Lymph node metastasis assigned as PTC. CDK4 modification profiles defined as described in Supplementary Fig. S1. Computed 95% confidence ellipses for each sample subtype (except for FTC and oncocytic FTC) are plotted. Samples discarded from further analysis are labelled as problem. The same representation is presented on the lower panel, showing each sample RNA-seq ID.

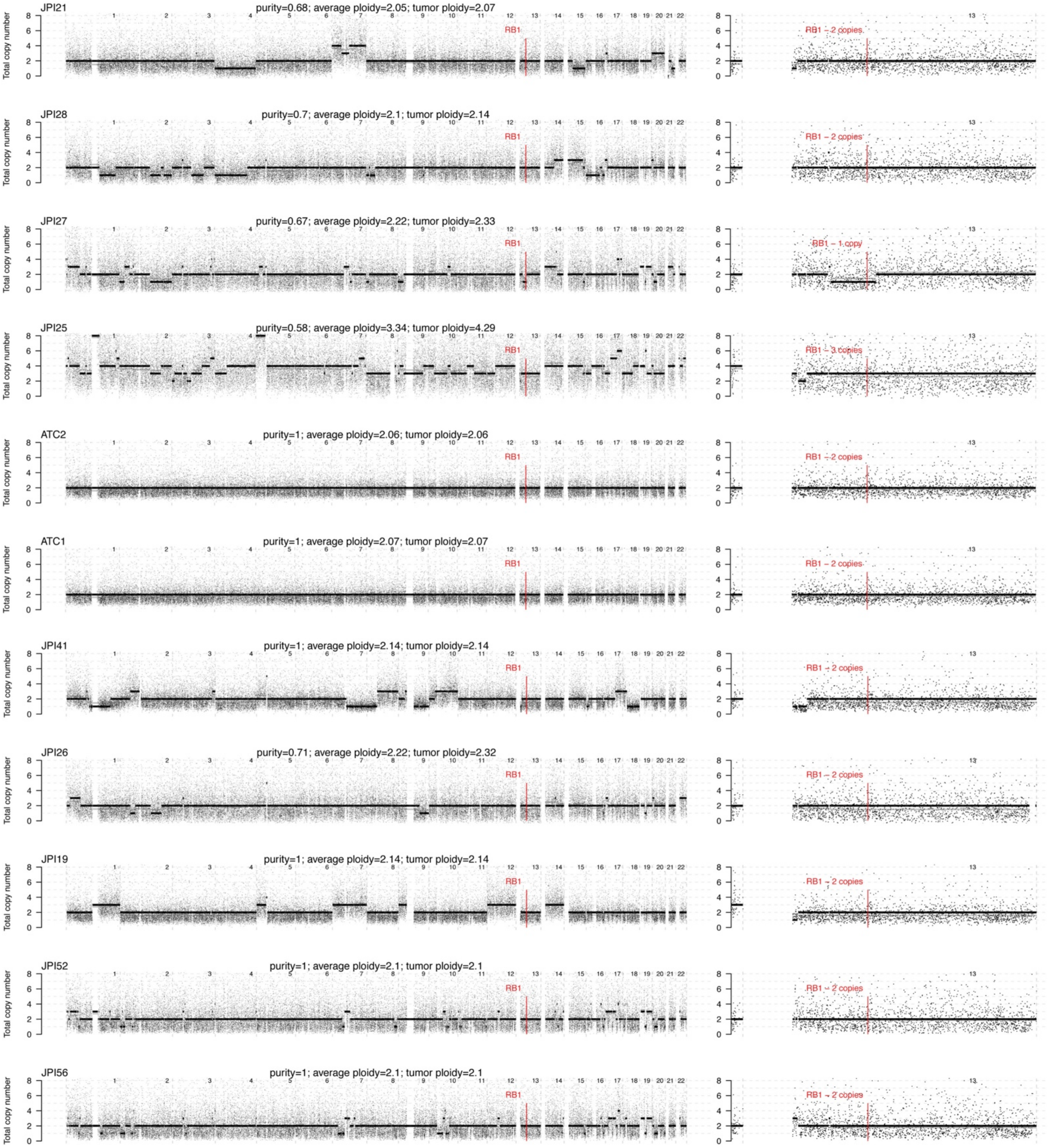


**Supplementary Figure S3. Genome-wide copy number profiles of profile A tumors.** DNA-targeted sequencing was performed for four profile A tumors with high p16 expression (JPI21, JPI28, JPI27 and JPI25) and two profile A tumors not RNA-sequenced (ATC2 and ATC1). For comparison, three profile H or L tumors (JPI41, JPI26 and JPI19) and two normal thyroid tissues (JPI52 and JPI56) are also presented. (On the left) Genome-wide total copy number profile across chromosomes is shown. The raw underlying data is shown for each 30kb bins, the raw segment average (in gray) and the fitted rounded integer value (in black); (On the right) Zoom in chromosome 13 showing the number of copies of *RB1*.

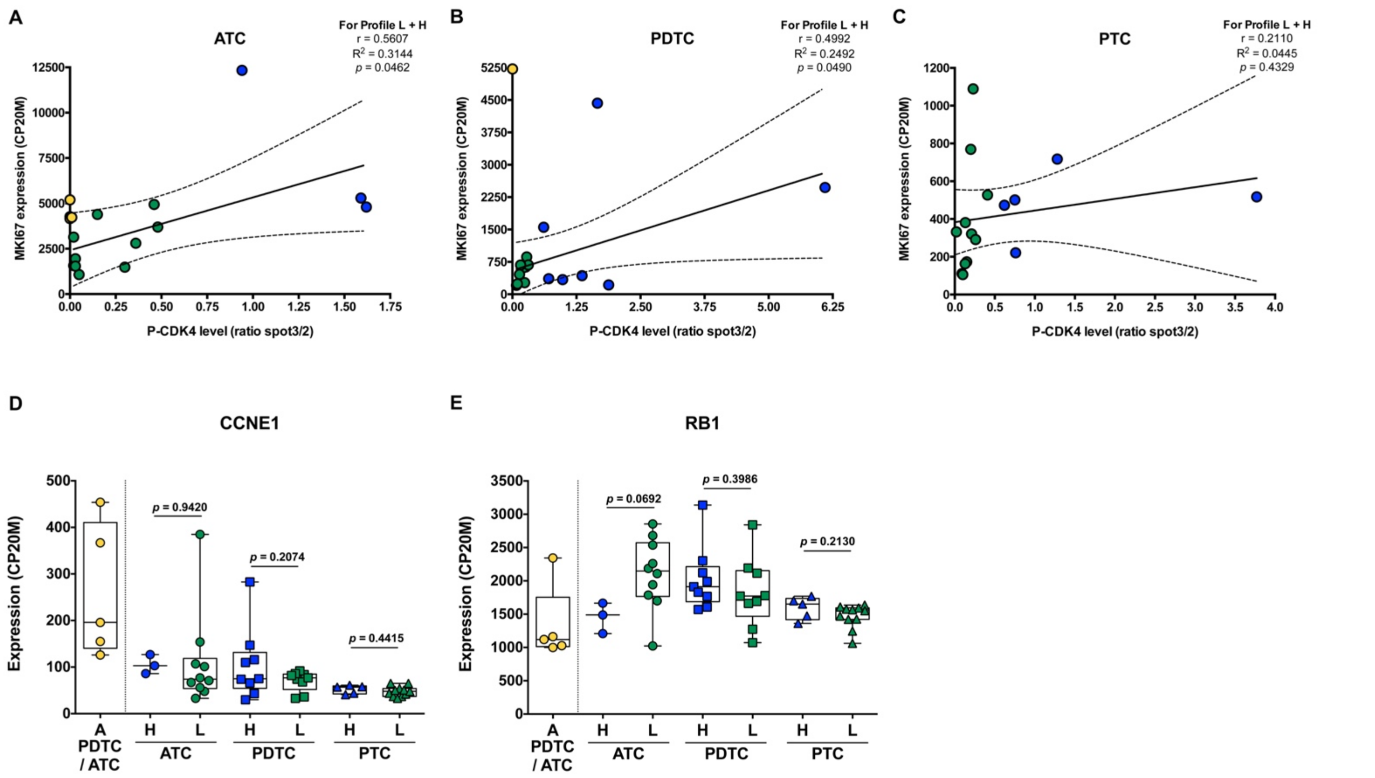


**Supplementary Figure S5. CDK4 phosphorylation levels in thyroid tumors are associated to different molecular features.**

(A-C) Correlation between CDK4 phosphorylation level (ratio of T172-phosphorylated form (spot 3) over spot 2, quantified from the tissue samples immunoblots) and *MKI67* expression (calculated from the RNA-seq data and expressed as counts per 20 million reads) in ATC, PDTC and PTC (lymph node metastasis assigned to PTC). Samples with CDK4 modification profile H (ratio ≥ 0.5), profile L (0.02 ≤ ratio < 0.5) and profile A (ratio < 0.02) are colored in blue, green and yellow, respectively. Data for samples with profile H and L were modelled by linear regression (best-fit values as solid line and 95% confidence bands as dashed lines). Pearson correlation coefficient (r), coefficient of determination ($R^2$) and correspondent statistical significance are shown.

(D, E) Expression levels of *CCNE1* (cyclin E1) and *RB1* in the samples grouped according to their CDK4 modification profiles (A, H or L as defined in Supplementary Fig. S1) in each subtype of tumors (lymph node metastasis assigned to PTC). Expression levels calculated from the RNA-seq data and expressed as counts per 20 million reads (CP20M). Statistical significance between profile H and profile L groups in each type of tumors were calculated with unpaired t-test.

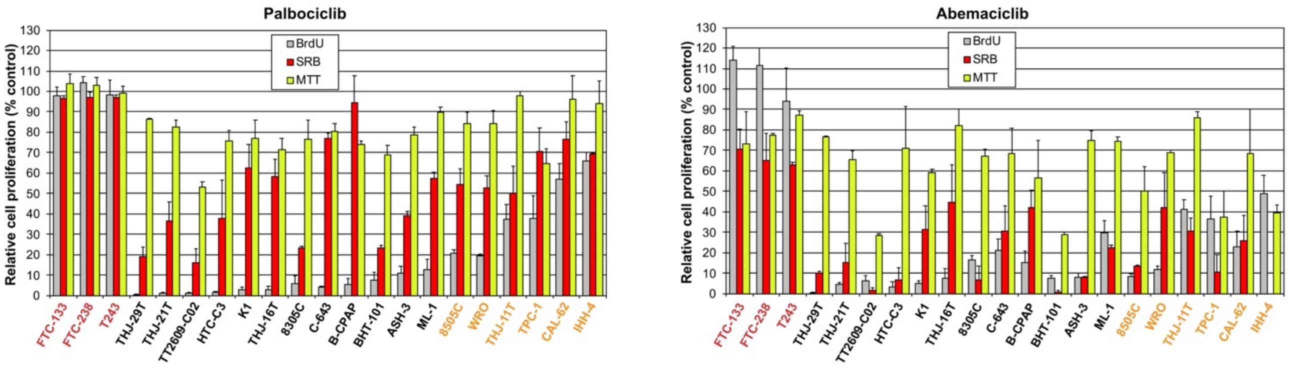


**Supplementary Figure S6. Comparison between BrdU incorporation, MTT and SRB assays.** Cell proliferation effect of CDK4/6 inhibitors on thyroid cancer cell lines was evaluated following 24 h, 48 h and 6 d treatment with 1 µM of drug, respectively. Pooled data from at least two independent experiments (error bars: mean ± SEM). Cell lines defined as resistant, sensitive with residual proliferation or sensitive (as detailed in Fig. 2) are indicated by red, orange or black font colors, respectively.

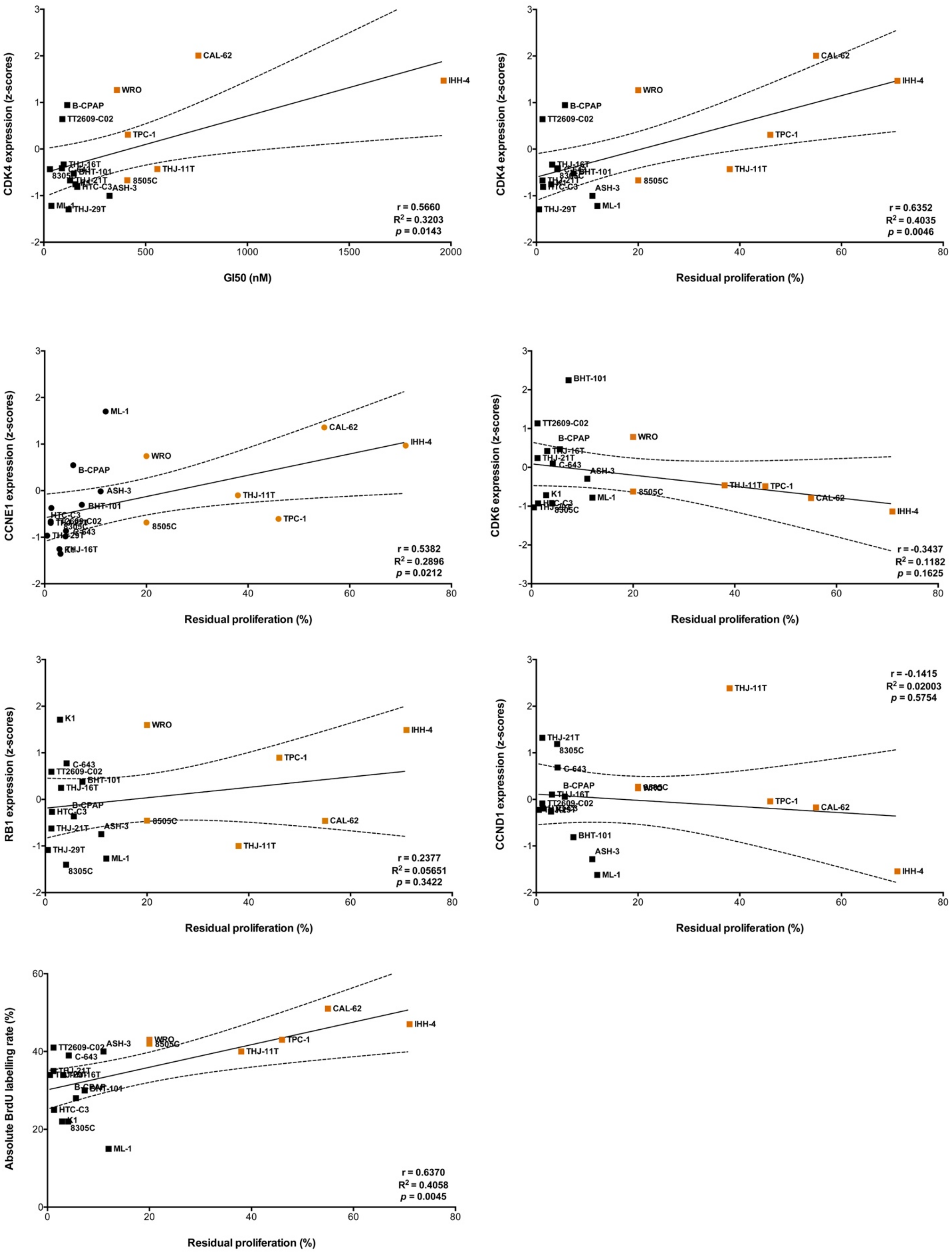


**Supplementary Figure S7. Correlation in thyroid cancer cell lines between palbociclib response and expression of putative markers of response.** Residual proliferation was defined as average relative BrdU labelling rate after 24h treatment with 1μM palbociclib. Half-maximal inhibition of cell proliferation (GI50) is shown as best-fit value. Expression levels of *CDK4*, *CCNE1* (cyclin E1), *CDK6*, *RB1* and *CCND1* (cyclin D1) calculated from the RNA-seq data and expressed as z-scores of counts per 20 million reads. Baseline proliferation was defined as average absolute BrdU labelling rate in control conditions. Data were modelled by linear regression (best-fit values as solid line and 95% confidence bands as dashed lines). Pearson correlation coefficient (r), coefficient of determination ($R^2$) and correspondent statistical significance are shown. Cell lines defined as sensitive with residual proliferation (as detailed in Fig. 2) are colored in orange.

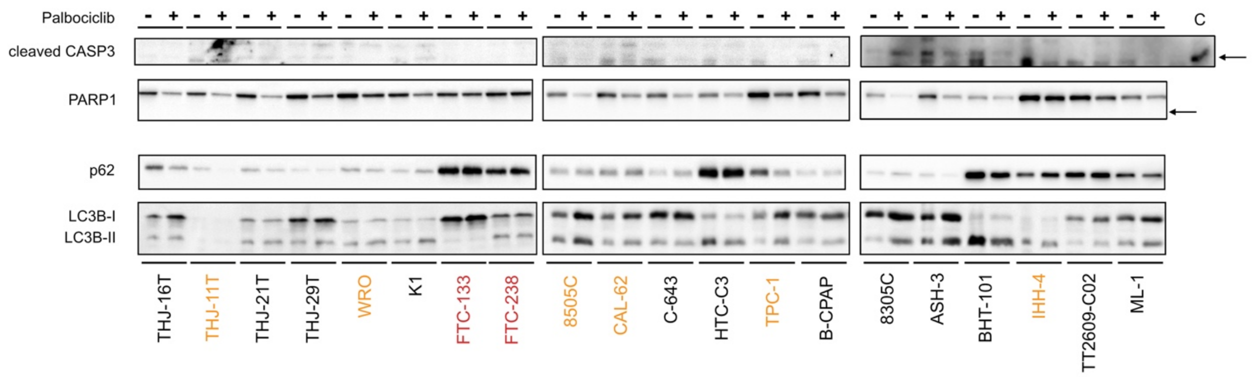


**Supplementary Figure S8. Treatment with palbociclib does not induce cell apoptosis/autophagy.** Indicated proteins were immunodetected after SDS-PAGE of total protein extracts from cells treated for 4 d with vehicle or with 1 µM palbociclib. Cells treated for 20 h with 10 µM etoposide (lane C) were used as a positive control for cleaved caspase 3 detection (black arrow). Cell lines defined as resistant, with residual proliferation or sensitive (as detailed in Fig. 2) are indicated by red, orange or black font colors, respectively.

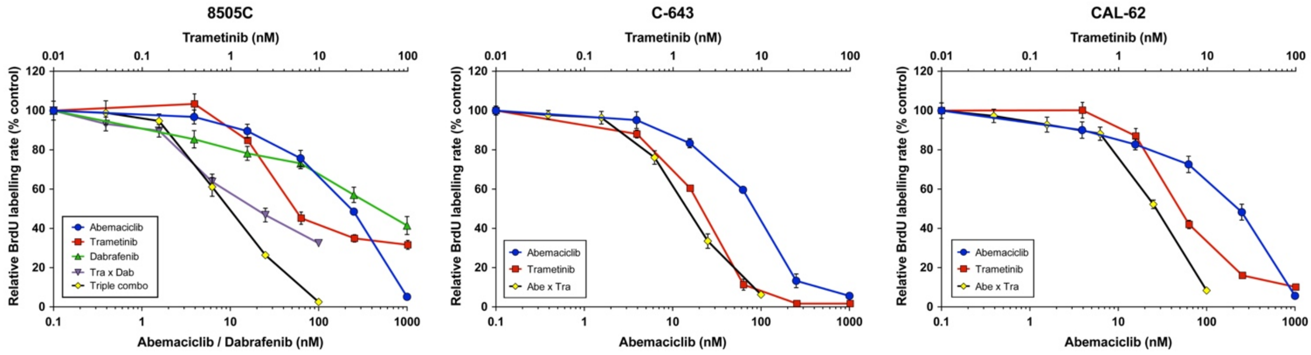


**Supplementary Figure S9. Sensitivity of thyroid cancer cell lines to the combination of abemaciclib with MEK/BRAF inhibitors.** Response curves (measured as BrdU incorporation during a 1 h pulse) in three ATC-derived cell lines, following 24 h treatment with serial dilutions of dabrafenib (dab, BRAF inhibitor), trametinib (tra, MEK inhibitor) or abemaciclib (abe) either alone, in combination of two or in combination of three drugs (triple combo). Error bars: mean ± SD (of triplicates, n = 1).

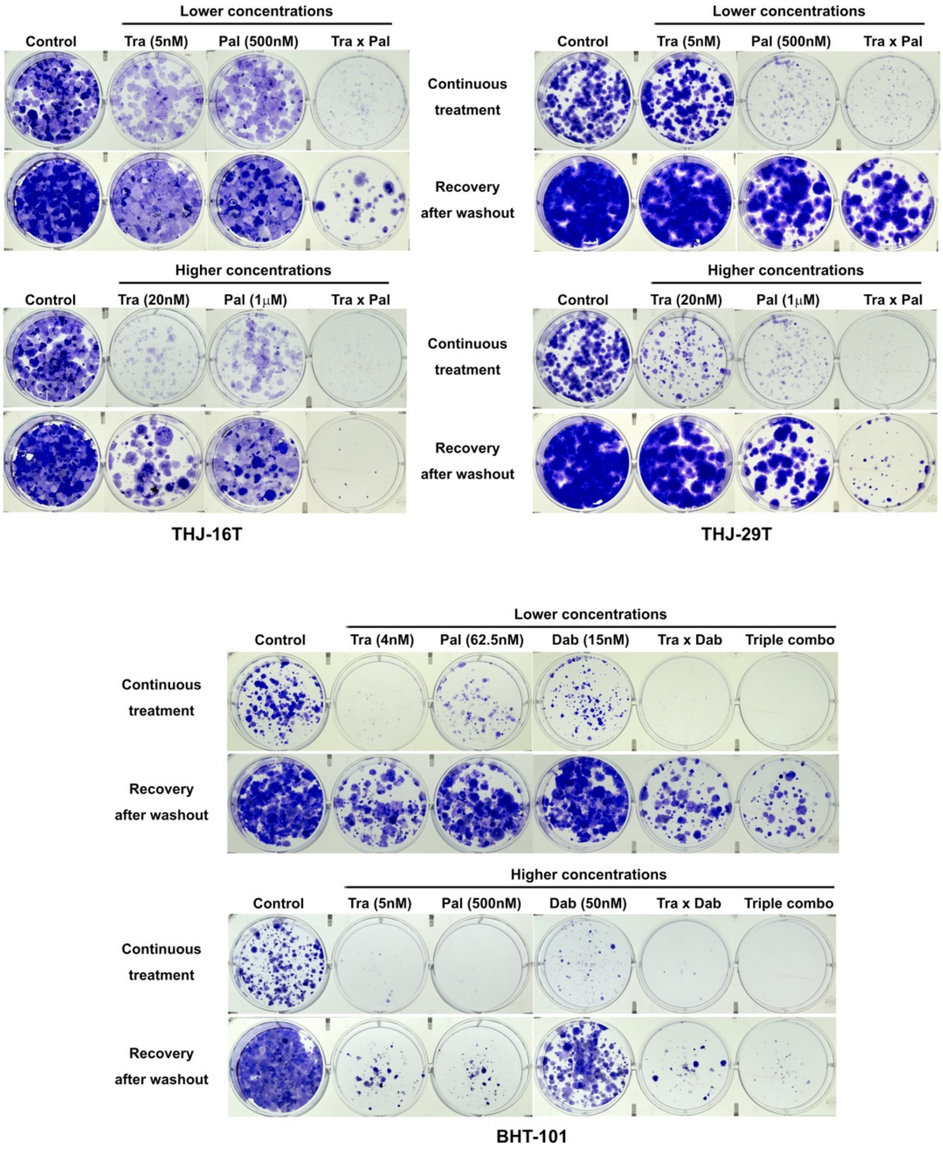


**Supplementary Figure S10. Long-term effect of the combined CDK4/6 and MEK/BRAF inhibition in additional thyroid cancer cell lines.** Clonogenic potentials of ATC-derived cell lines treated for 10 d at indicated concentrations with vehicle, dabrafenib (dab, BRAF inhibitor), trametinib (tra, MEK inhibitor) or palbociclib (pal) either alone, in combination of two or of three drugs (triple combo), followed or not by a 10 d recovery period (drug withdrawal). Representative photographic images of two independent experiments.

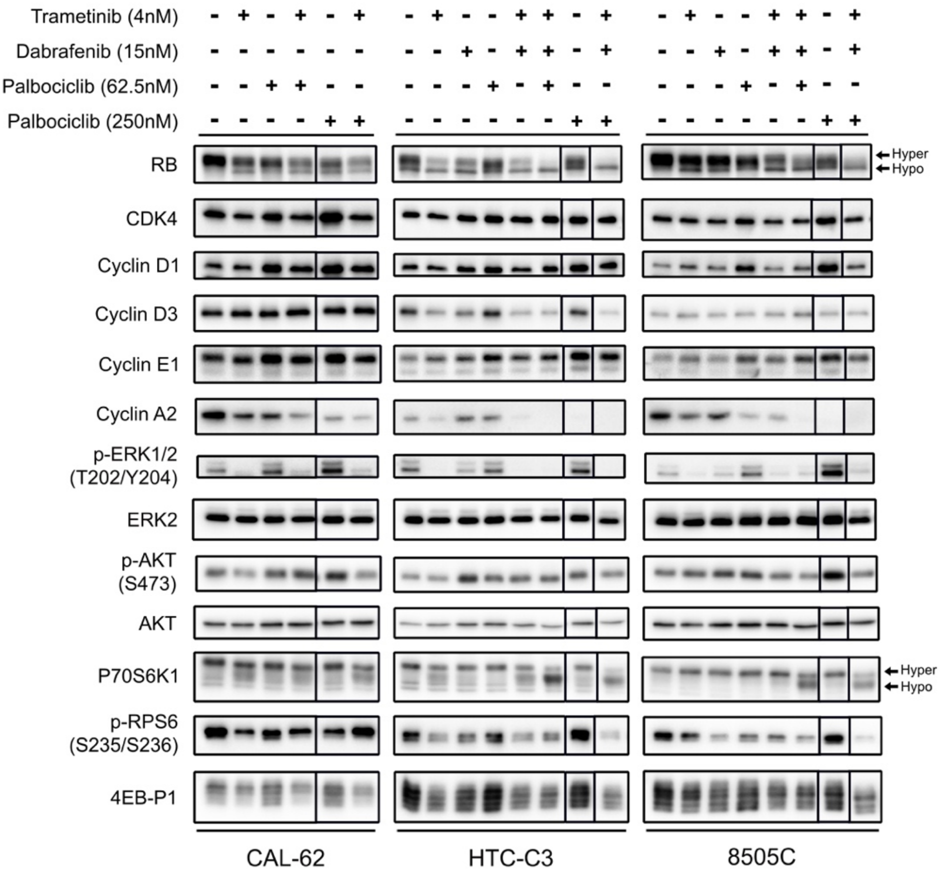


**Supplementary Figure S11. Impact of the CDK4/6, MEK and BRAF inhibitors on cell cycle- and signaling pathways-related protein levels.** Indicated proteins were immunodetected after SDS-PAGE of total protein extracts from cells treated for 3 d with vehicle or with indicated drugs. Solid lines separate parts of the same blot detection that were re-assembled. Hyper- and hypo-phosphorylated forms of RB and p70S6K1 are indicated.
